# Phylogenetic and Spatial Distribution of Evolutionary Isolation and Threat in Turtles and Crocodilians (Non-Avian Archosauromorphs)

**DOI:** 10.1101/607796

**Authors:** Timothy J. Colston, Pallavi Kulkarni, Walter Jetz, R. Alexander Pyron

## Abstract

The origin of turtles and crocodiles and their easily recognized body forms dates to the Triassic. Despite their long-term success, extant species diversity is low, and endangerment is extremely high compared to other terrestrial vertebrate groups, with ~ 65% of ~25 crocodilian and ~360 turtle species now threatened by exploitation and habitat loss. Here, we combine available molecular and morphological evidence with machine learning algorithms to present a phylogenetically-informed, comprehensive assessment of diversification, threat status, and evolutionary distinctiveness of all extant species. In contrast to other terrestrial vertebrates and their own diversity in the fossil record, extant turtles and crocodilians have not experienced any mass extinctions or shifts in diversification rate, or any significant jumps in rates of body-size evolution over time. We predict threat for 114 as-yet unassessed or data-deficient species and identify a concentration of threatened crocodile and turtle species in South and Southeast Asia, western Africa, and the eastern Amazon. We find that unlike other terrestrial vertebrate groups, extinction risk increases with evolutionary distinctiveness: a disproportionate amount of phylogenetic diversity is concentrated in evolutionarily isolated, at-risk taxa, particularly those with small geographic ranges. Our findings highlight the important role of geographic determinants of extinction risk, particularly those resulting from anthropogenic habitat-disturbance, which affect species across body sizes and ecologies.

## Introduction

Turtles and crocodilians (non-avian Archosauromorpha) are among the most charismatic and widely recognized of all living things (1). Both groups date to the Triassic, and exhibit highly derived, yet highly conserved body forms. Crocodilians are famous for their extraordinary size (up to 6m and 1,000kg), long snouts and tails, and bony armor under the skin; and turtles for their bony or cartilaginous shell, and the size of some marine and terrestrial species (1.4m and 400kg on land, 2m and 1,000kg in the sea). Both groups are well-represented in the fossil record and previously attained even more massive sizes, up to 12m and 8,000kg for the extinct crocodilian *Sarcosuchus* (2) and 5m and 2,200kg for the extinct turtle *Archelon* (3). In comparison to their terrestrial vertebrate relatives (birds, mammals, reptiles, and amphibians), both groups have very few species (~360 turtles and ~25 crocodilians), and a large proportion of extant species are both highly evolutionarily distinct (4) and highly endangered (5, 6), with the Madagascan Big-Headed Turtle (*Erymnochelys*) having the highest EDGE (Evolutionarily Distinct, Globally Endangered) score of any terrestrial vertebrate (4).

Turtles and crocodilians present several interesting hypotheses regarding historical diversification. First, the group exhibits relative constancy in the fossil record during the Cretaceous-Paleogene extinction (7–12). Therefore, we test if there is any signature of a K-Pg mass-extinction event or shift in diversification rates either prior or subsequent to the K-Pg boundary. Such a pattern may be relevant if extinction risk is concentrated in older or younger taxa, or specific lineages crossing the boundary. Second, the general body-form of each group (bony-shelled turtles, armored lizard-like crocodilians) dates to the Triassic, but morphological disparity is much higher among extinct taxa for both, with a greater diversity of body forms in fossil species than observed today (13, 14). Thus, we test if significant shifts in body-size evolution are detectable among extant lineages of turtles and crocodilians as found in some previous studies (15, 16), in comparison to the drastic variation seen in their fossil relatives (13, 14). We hypothesize that such shifts may offer partial explanations for the present-day apportionment of ED and thus extinction risk among species, particularly if body size is related to threat status as in amphibians (17), birds (18), mammals (19, 20), and squamates (21).

Despite their visibility as wildlife and study organisms, to date there have been no taxonomically-complete, dated phylogenies produced for turtles and crocodilians. Such trees are invaluable for quantifying the evolutionary distinctiveness of extant species (22, 23), testing hypotheses about historical rates of diversification and body-size evolution (24, 25), predicting threat status for data-deficient (DD) and unassessed (UA) species (17, 26), and quantifying the spatial and phylogenetic distribution of extinction risk (27, 28). The latter two aims are particularly crucial given the higher proportion (65% of all species) of threatened turtles and crocodilians (29) compared to groups such as amphibians with ~33-50% threatened (17), mammals ~25%, birds ~12%, and squamate reptiles ~12% based on the most recent assessments (30). A recent study estimated and imputed ED and EDGE scores based on available phylogenetic datasets (4), but the scanty taxonomic coverage of the trees and the imputation methods employed by those authors leaves a wide margin for error.

Here, we use taxonomically-complete, dated phylogenetic estimates to calculate ED from “fair proportion,” a measure of the evolutionary isolation of each species and correlate of net diversification rates (22). Species with high ED represent relict lineages on long branches, whereas groups of low-ED species represent more recent, rapid radiations; ED is thus correlated with and reflects historical diversification patterns, in the branch lengths of extant species (31). These data are invaluable for identifying conservation priorities (32), given the necessity of triage when allocating limited resources for management and further research (33). Indeed, at least nine species of turtles are thought to have gone extinct in modern times (34), a higher percentage (2.5%) than birds (1.9%), mammals (1.7%), amphibians (0.4%), or squamate reptiles (0.2%) among terrestrial vertebrates which have been assessed (30), while dozens more turtles and crocodilians are critically endangered throughout the world (35).

Second, we gather spatial and trait data for all species to identify the key factors related to extinction risk and build predictive models to estimate threat status for DD and UA species (26, 36). While 23/27 crocodilians have been assessed (85%) of which 11 are threatened (48%), only 258/357 turtles (72%) have been assessed, of which 182 are threatened (71%). Thus, half to two-thirds of all assessed crocodilian and turtle species are threatened, but a quarter of all species are without robust assessment or prediction (35). Many crocodilians and turtles have traits making them particularly susceptible to anthropogenic threats such as small geographic ranges (37), long generation times (38), exploitation for food and commerce (39), reliance on fragile habitats such as coastlines (40), and occurrence near human-population centers such as South and Southeast Asia, the eastern Nearctic, and coastal Australia (35). We use a variety of statistical and machine-learning methods to identify the strongest correlates of known extinction risk and predict threat statuses for the remaining DD and UA species (17, 41).

These data allow us to answer several pressing questions regarding extinction risk at the global scale. Is evolutionary distinctiveness related to threat status? No such pattern exists for birds (23), mammals (42), amphibians (28), or squamate reptiles (43), albeit with limited data for some of these groups. However, many lineages of turtles such as the Fly River or Pig-Nosed Turtle (*Carettochelys*) and the Big-Headed Amazon River Turtle (*Peltocephalus*) are known to be threatened and occupy isolated phylogenetic positions (35, 44). In contrast, the most ancient crocodilian species (45) are the alligators (*Alligator*), for which the American Alligator is Least Concern, while the Chinese Alligator is Critically Endangered (30). Which are the most distinct and endangered species when combining ED and threat status (4)? In terms of spatial distribution, are there geographic hotspots of concentrated threat (37), or geographic “arks” with high diversity of non-threatened species and improving human footprint (46)?

Here, we produce a posterior distribution of 10,000 taxonomically complete Bayesian time-trees representing all 357 recent species of turtles and 27 species-level crocodilian lineages. We test for significant temporal variation in speciation and extinction rates related to the K-Pg boundary, as well as among-lineage variation in body-size evolution. We then estimate threat statuses for 114 DD and UA species based on models incorporating spatial, phylogenetic, and trait data, and relate extinction risk to ED across taxa and in different areas. In contrast to the massive variation seen in the fossil record, the phylogeny of extant species shows little evidence for mass extinctions, changes in diversification rate, or shifts in body-size evolution. Unlike all other terrestrial vertebrates, we find that extinction risk is concentrated non-randomly among the most distinct and unique species, a worrying pattern that suggests current trends will erase a disproportionate amount of evolutionary history. These risks are concentrated both in specific lineages and particular geographic areas, while other clades and regions represent relative havens for diversity. Together, these data paint a picture of urgency for the conservation of select species of high distinctiveness and high extinction risk.

## Materials & Methods

### Phylogeny Construction

Detailed materials and methods are available in the online supplement. We used well-established methods for supermatrix estimation and taxonomic imputation to create a posterior distribution of trees containing full taxonomic sampling for turtles and crocodilians (25, 28, 43, 47). Using the 8^th^ edition of Turtles of the World (34), we recognized 357 species of extant or recently extinct turtle. Based on several recent references, we recognized 26 described crocodilian species plus one undescribed *Osteolaemus* from Western Africa (48). We sampled 14 mitochondrial loci; 12S, 16S, ATP6, ATP8, COI, COII, COIII, CYTB, DLOOP, ND1, ND2, ND3, ND4, and ND5; and six nuclear loci; BDNF, CMOS, GADPH, ODC, RAG1, and RAG2, for all taxa. We sampled R35 for turtles only, as the intron has not been widely sequenced in other groups. This yielded 7 total nuclear loci, and 21 genes overall. Sequence data were available for 340 of 357 turtles, all crocodilians, and the two outgroups. The matrix was 24,798 bp in total, ranging from 405 to 20,929 bp per species, and averaging 8,891 bp for 369 terminals. We included *Gallus gallus* (Aves) and *Sphenodon punctatus* (Lepidosauromorpha) as outgroups.

From this sparsely-sampled supermatrix, we estimated a Maximum Likelihood topology in RAxML (49), which we enforced as a constraint for a PASTIS (47) run in MrBayes (50) to estimate node ages and impute the placement of the seventeen missing turtle species. Specific taxonomic information on the placement of these imputed taxa is provided in the supplemental information. We placed constraints on the ages of seven key nodes, using consensus age-estimates from the Time Tree of Life database (see supplement). We estimated the clock-rate prior using standard approaches (51) and used a pure-birth model for the branch-length priors to remain conservative with respect to the historical signature of extinction (25).

This approach yielded a posterior distribution of 10,000 taxonomically-complete, dated phylogenies. While some authors caution against using phylogenies with imputed taxa to measure rates of character evolution (52), we note that i) the number of imputed taxa is extremely small (17/384; ~4%), which should reduce any bias in estimated rates; and ii) those authors also found that estimated diversification rates provided stable inferences (53). Thus, these phylogenies can thus be used for myriad downstream comparative analyses, such as estimating diversification regimes, predicting threat status, and testing for phylogenetic signal. We do caution that issues of taxonomic non-equivalence in species-level taxa among different turtle groups may hamper some estimates of diversification (see below & supplement).

### Diversification Analyses

To evaluate temporal variation in diversification rate, we used algorithms implemented in the TESS package (54), which estimate speciation and extinction rates through time across a clade while allowing for shifts, as well as testing for mass extinctions. We do not specify the hypothesized location of any rate shifts or mass extinctions beforehand, and the model is thus agnostic to their temporal location. However, we generally hypothesize that if significant shifts or extinction periods are observed, they would be localized near the K-Pg boundary. Specifically, we inferred rate estimates for speciation and extinction with a Bayesian variable ‘birth–death’ model (55), conditioned on turtles and crocodilians separately. Jointly with this analysis, we also searched for potential evidence of past mass extinctions using the ‘compound Poisson process on mass-extinction analysis’ or ‘CoMet’ model (56).

We sampled ten trees from the posterior of each of the two clades and modelled a variable ‘birth–death processes’ with explicit mass-extinction events following the guidelines in (54), with the sampling fraction set to 1 and the expected survival of a mass extinction equal to 0.05 (95% extinction). We generated empirical hyperpriors through an initial Bayesian Markov chain Monte Carlo analysis under a constant-rate birth–death-process model to determine reasonable hyperparameter values for the diversification priors. We allowed up to two mass-extinction events and two rate change events and set the shape parameter for the mass-extinction prior to 100; varying these parameters yielded qualitatively similar results. After burnin, we ran final analyses for 100,000 generations and diagnosed models using effective sample size (>200) and the Geweke statistic. We evaluated significance of rate shifts and mass extinctions using Bayes factors, mentioning instances >6 and considering >10 strong support.

### Body-Size Evolution

We used the program *levolution*, which models the rate, strength, and phylogenetic position of evolutionary jumps using a Lévy process (57) to identify saltational changes in Brownian Motion regimes in a trait across a tree (58). This approach has a variety of statistical, theoretical, and computational advantages over existing methods (59). These include computational simplicity and reduced computing time, greater identifiability of parameters, and better theoretical fit to instances of rapid evolutionary bursts punctuating apparent stasis in phenotype. We only evaluated the length data (including the single imputed value for *Osteolaemus* sp.), as analyzing the phylogenetically-imputed masses on the same phylogeny would be too circular for comfort, while the observed data alone would be too sparse for confidence in the results. For details about how these data were gathered see *Trait Data*, below and the supplement.

We first estimated the value of the hyperparameter α using the included dynamic peak-finder algorithm, which permutes α until the highest likelihood is found. We used the recommended settings of 5000 sampled jump-vectors, 1000 burnin samples, and 2nd-generation thinning for the MCMC chains, and a starting value and step value of 0.5 (on a log scale), optimized 5 times. The program thus tested 14 values of α, with 31.6228 being the final value. The LRT was significant (*D* = 175, *P* < 1e-16), suggesting that a jump-diffusion Lévy process best fits the data. We then ran a final analysis with α fixed to this value, starting from the ML estimates of the mean, variance, and jump-rate parameters. For this final analysis, we increased the number of sampled vectors in the MCMC chain to 25000 for both the parameter estimation and the jump inference, accepting jumps with posterior probabilities >0.85.

### Trait, Spatial, and Threat Data

For the purposes of analyzing morphological evolution and ecological correlates of extinction risk, we gathered several traits shown to be relevant in other terrestrial vertebrate groups such as birds, mammals, amphibians, and reptiles (17, 21, 26, 36, 41, 60). Specifically, we measured body length (carapace length [CL] for turtles or total length [TL] in cm for crocodilians), body mass (g), microhabitat (freshwater, terrestrial, or marine), endemicity (island or mainland), biogeographic area (from 11 global ecoregions encompassing extant diversity), range size (km^2^) and proximity (distance matrix of range centroids) from existing spatial datasets (61), human-encroachment index (proportion of urban or cropland in species’ geographic range at 300m resolution (62)), climate (BIOCLIM 36-40 measuring principal components of temperature, precipitation, and radiation at 10-min resolution (63)), and ecosystem function (AET and NPP from the MODIS datasets at 500m resolution (64)).

We obtained maximum length values for all 357 turtle species and 26/27 crocodilians, and for body mass we gathered values for 202 turtle species and 23 crocodilians. All other data were available for all species, with the exception of BIOCLIM 36-40 for three island-endemic turtles. We imputed TL for a single species, BIOCLIM 36-40 for three species, and body mass for 159 species using phylogenetic techniques (65). Spatial data were generated from polygon shapefiles representing the geographic range of all species, derived from published sources (34, 61, 66) and projected globally using a World Cylindrical Equal Area projection (EPSG:54034).

For threat status, we first accessed the 29 November 2018 version of the IUCN RedList. We then changed *Nilssonia nigricans* from EW to CR because wild populations have recently been rediscovered (67), and assigned *Aldabrachelys abrupta* and *A. grandidieri* to EX, based on their known extinction (68). Thus, there are 54 Least Concern, 34 Near Threatened, 72 Vulnerable, 48 Endangered, 53 Critically Endangered, 9 Extinct, and 11 Data Deficient species. These account for 258/357 turtles and 23/27 crocodilians, for a total of 281 assessments (270 with a known status) out of 384 total species. Remaining to impute are therefore 114 total species: 103 unassessed (99 turtles and 4 crocodilians) and 11 Data Deficient (all turtles). No species are currently Extinct in the Wild.

### ED, Status, and Geography

We calculated ED of all 384 species using the ‘evol.distinct (type = “fair.proportion”)’ function in the R package ‘picante’ (69) taking the median value per species across the posterior distribution of 10,000 trees. We then imputed the 114 DD or UA threat statuses using both traditional statistical methods (26) and novel machine-learning approaches (36). First, we generated linear models (17) linking the trait, range, climatic, and ecological data to status in the 270 assessed species using a GLM approach, while including terms for phylogenetic and spatial covariance. Continuous variables were log-transformed, centered, and scaled, while categorical predictors were converted using binary one-hot (1-of-*k*) encoding as presence or absence of each feature. Preliminary variable selection highlighted 15 significant predictors (see supplement). We then used PLGM estimators to impute the missing statuses as a continuous estimate from 1 to 6, corresponding to the six statuses (17, 26). We then estimated the sensitivity and specificity of this model using leave-one-out cross-validation, removing each of the 270 known statuses one-by-one, re-estimating them from the remaining data, and generating a confusion matrix of the known versus predicted statuses.

Second, we used the machine-learning (ML) techniques ‘random forests’ (RF) and ‘artificial neural networks’ (ANN), which have become widespread in ecology (70, 71) and conservation biology (70, 72), as implemented in the R package ‘caret’ (73). Advantages over approaches such as GLMs include higher classification accuracy, greater flexibility for addressing problems such as regression and classification in a single modeling framework, and the intrinsic ability to detect and incorporate complex non-linear interactions between variables without *a priori* specification (70, 71).

For the RF models, we optimized the ‘mtry’ variable over 500 trees including all 28 categorical and continuous predictors, plus 2-dimensional PCoA estimates of phylogenetic and spatial covariance, for a total of 32 predictors. Both ML techniques integrate out uninformative variables and are thus not particularly susceptible to overparameterization compared to classical statistical approaches such as PGLM. The RF methods are also insensitive to the relative scale of input features, which are assessed sequentially and thus do not interact, so the 32 predictors were not transformed prior to analysis. We assessed model accuracy using repeated *k*-fold cross-validation and assessment of relative variable-importance.

For the ANN models, we used feed-forward multi-layer perceptrons with a single hidden layer, trained using back-propagation under a root-mean-squared-error cost function from the 270 species with known status and the 32 categorical variables. Predictor variables were standardized on the interval [0-1] using min-max scaling. We tuned the model using repeated *k*-fold cross-validation to select the number of neurons in the hidden layer and the weight-decay parameter to regularize the weights across neurons. These weights were then used to calculate variable importance for input features.

Estimates from all three methods were then pooled across species, and the cross-correlation and pairwise identity were assessed across methods. For the final estimate of each species, we took the “mean” estimate on the interval [1-6], though most methods agreed for most species (see Results). Pooling the assessed and imputed statuses for all 384 species, we then tested for a relationship between ED and extinction risk across categories using the notch test on the associated bar-plot, and between threatened and non-threatened species using a two-sample *t*-test. Despite the fact that threat status (or at least, many of the underlying predictors) may exhibit strong phylogenetic signal, this is explicitly a non-phylogenetic test, because we are solely interested in whether the correlation exists at time zero given current conditions, for each species independently.

## Results

### Phylogeny and Diversification

All data and results, including the tree topology with support values (Appendix S1), are available in the Supplemental Material and DataDryad repository (to be deposited upon publication; contact TJC or RAP for data). Our phylogeny is robust overall, with monophyly of all families, subfamilies, and genera strongly supported. Of particular note is the continued weak support for relationships among tortoise genera (Testudinidae), as well as broader uncertainty in higher-level relationships among and placement of Cheloniidae, Chelydridae, Emydidae, and Platysternidae (44, 74–76).

We recover a unique topology from these previous four analyses, with successive divergences of Chelydridae+Dermatemydidae+Kinosternidae, Cheloniidae+Dermochelyidae, Platysternidae, Emydidae, and Testudinidae+Geoemydidae (77). This is based on the same or similar underlying datasets and may reflect issues such as mito-nuclear discordance and phylogenetic signal in the available loci. However, our increase in both taxon and characters sampling may indicate convergence on a more robust topology. Future analyses sampling more loci may be able to resolve these relationships with greater support.

Several turtle clades have been identified by previous authors (15, 16, 74) as representing extraordinary instances of evolutionary diversification, including Galapagos tortoises (*Chelonoidis*), and New World emydids (Deirochelyinae). Similar results were recovered in our preliminary analyses. However, we believe strongly that these are artifactual, and represent an inconsistent application of species concepts and delimitation in turtles. At the current juncture, we are constrained by existing taxonomic frameworks, and these issues must thus be left to future studies. However, interpretation of our downstream results (see below) should be colored by knowledge of these issues. We attempt, when possible, to use analyses and interpret models in a way that is resilient to the possibility of low-level taxonomic biases.

Results for turtles and crocodilians showed roughly constant rates of speciation and extinction, with no support for any shifts therein, or any mass-extinction events (Fig. 1a). Turtles exhibit speciation rates of ~0.07 lineages per million years and extinction rates of ~0.03-0.04, for net diversification of ~0.03-0.04 and turnover probabilities of ~43-57% from the Jurassic to the present. For crocodilians, rates are ~0.05 and ~0.02 respectively, for net diversification of ~0.03 and turnover of ~40% from the Cretaceous to the present. For body size, few strongly-supported jumps are estimated by the model (Fig. 1b). Thus, most variation can be attributed to steady drift over time. One jump occurs along the branch leading to extant crocodilians, reflecting the difference in body size between the two groups, crocodilians being longer and heavier on average. A second occurs in the relict species *Carettochelys insculpta*, a large freshwater species, which is the sister lineage of the radiation of softshell turtles (Cyclanorbinae and Trionychinae) that contains both large and medium sized species. Three terminal species are also estimated as jumps, each of which is significantly smaller than its congeners. These are *Pelodiscus parviformis, Pseudemys gorzugi*, and *Trachemys adiutrix*. As noted above, these radiations and *Pelodiscus* have questionable species boundaries (78). Thus, we refrain from interpreting these results further.

**Figure 1.**
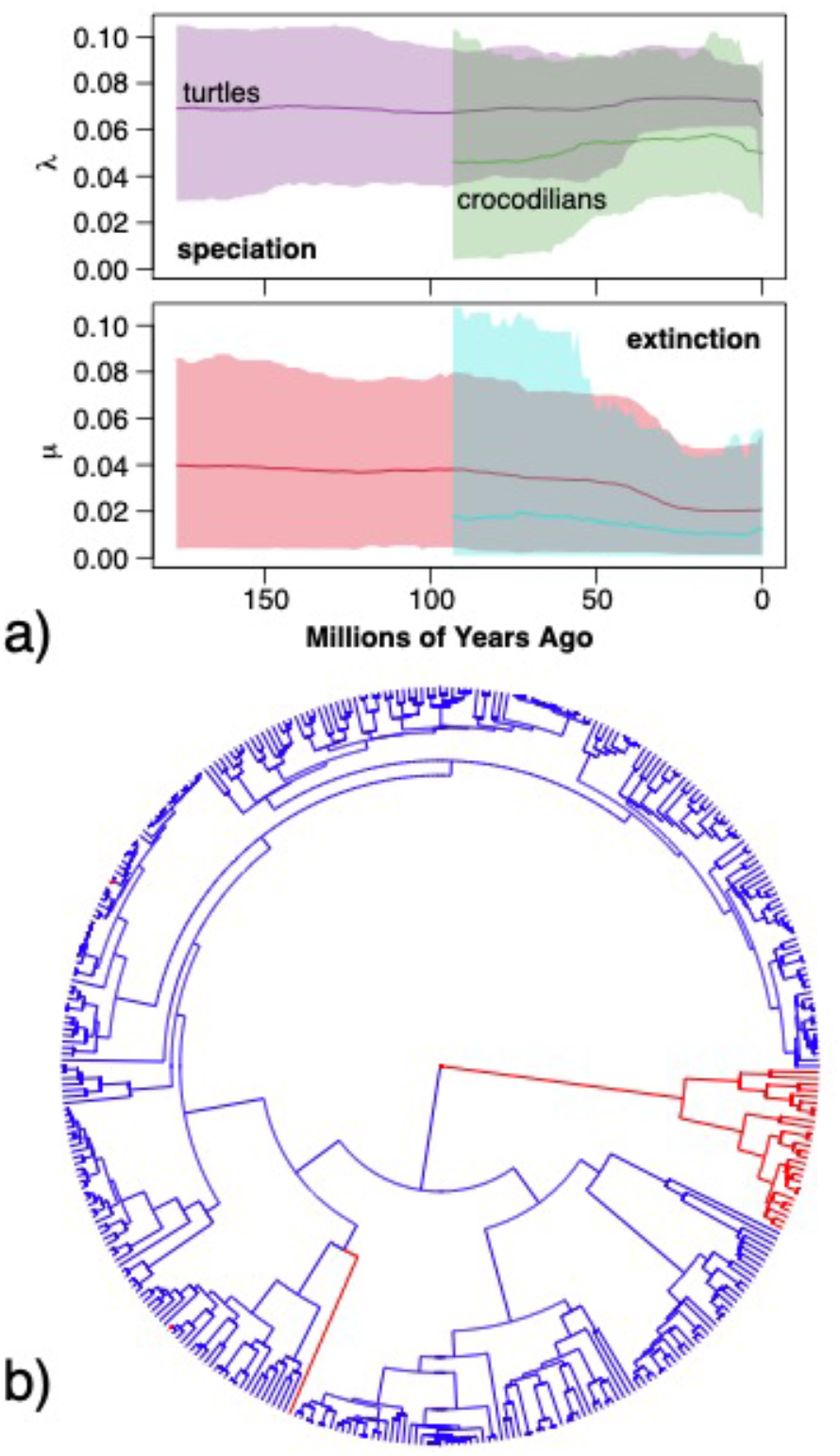
Results from (a) TESS/CoMET analysis showing estimated speciation and extinction rates for turtles and crocodilians across time; and (b) location of significant jumps (red) inferred under a Lévy process in crocodilians, *Carettochelys*, and three other species.

### ED and Threat Status

We find a bi-modal distribution of ED values for extant turtles (Fig. 2). The primary mode is 17Ma, older than amphibians at 16.5Ma (28), squamate reptiles at 11Ma (43), birds at 6.2Ma (23, 25), or mammals at 4.8Ma (79). A secondary mode at 5Ma is dominated by the recent “radiations” in Deirochelyinae and Testudininae (see Discussion), for which multi-locus nuclear datasets generally do not support the higher number of morphologically-delimited lineages. Thus, we refrain from examining among-lineage rate variation for speciation or extinction.

**Figure 2.**
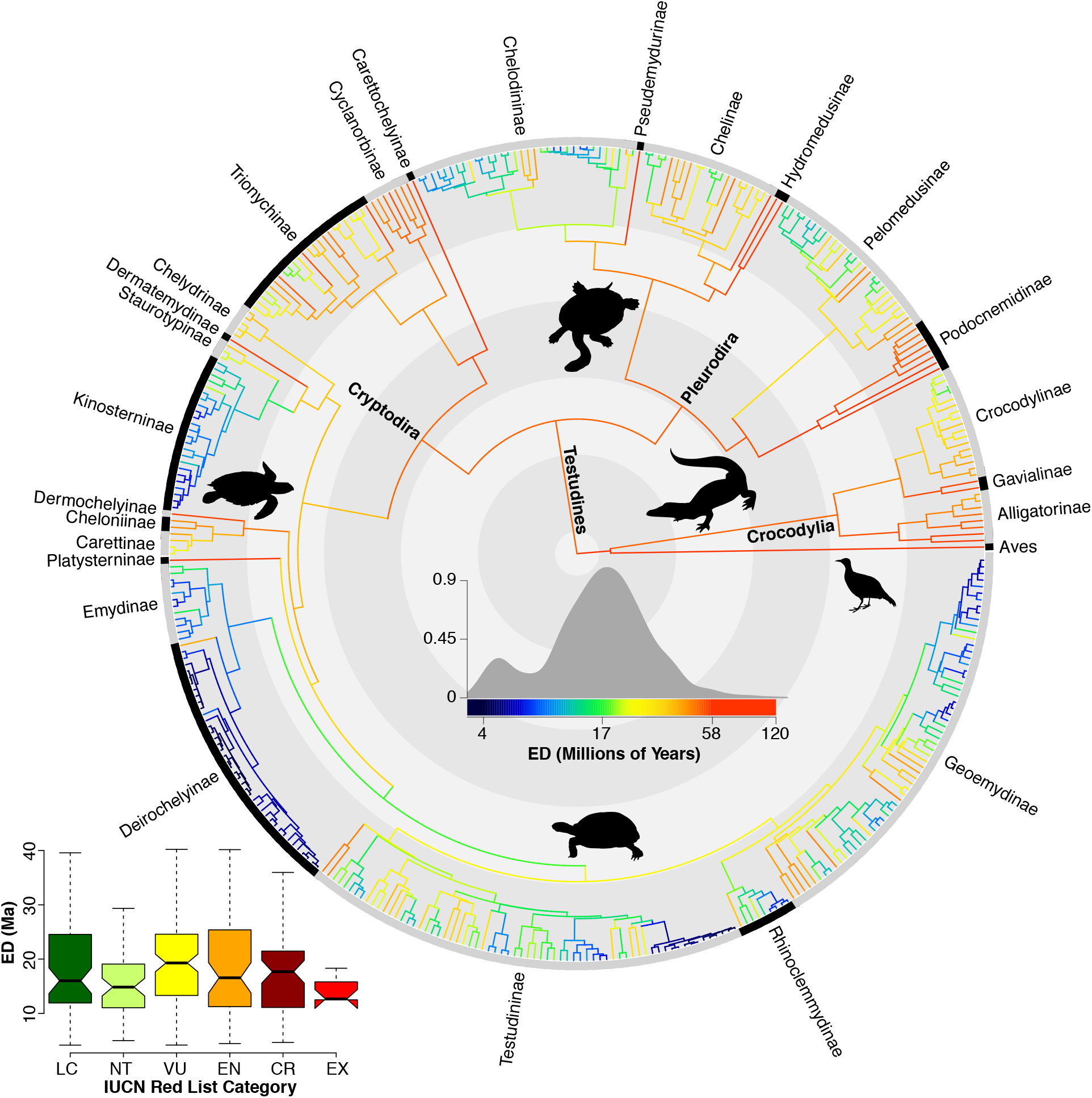
Randomly-selected phylogeny, with branches colored proportional to ED for extant lineages. The internal contrasting rings denote 50Ma intervals. The insets show the overall distribution of ED across 356 turtle and 27 crocodilian species, and the distribution of ED across threat statuses, including both assessed and imputed species. Silhouettes are from http://ww.phylopic.org/; see Supplemental Information for license details.

Across the three methods for threat-status imputation, the final predictions for the 114 species are fairly similar (Appendix S5), with all models identifying area as the single-most important variable by far (Figs. S7-13), as in amphibians (17), birds (80), mammals (81), and squamate reptiles (21). Each method picked out slightly different contributions from secondary predictors. The second-ranked variable for the PGLM model was occurrence in the Oriental ecoregion; for the RF model there is a tie between AET (ecology) and HEI (anthropogenic disturbance); and for the ANN model it was spatial dissimilarity, indicating the geographic clustering of threat status (see below). Body size (length or mass) were not particularly important, unlike for amphibians (17), birds (18), mammals (19, 20), and some squamates (21, 82). Training accuracy for the three models was higher (PGLM = 38%, RF= 49%, ANN = 48%) than random (17%) and confusion matrices showed high sensitivity and specificity (up to 86%), with errors usually involving only a single step in either direction.

Overall, 21 species were predicted identically by all three approaches, and 69 agreed for two out of the three, for ~79% concordance overall. On a pairwise basis for 342 comparisons (114 species times three models), 141 were identical (41%) and an additional 144 (42%) were adjacent (one category different), for a total of 83% identical or adjacent predictions. Given the high level of agreement and lack of apparent bias towards higher or lower categories for any model, we simply used the mean of the three predictions for our final estimate of threat status.

Our final estimate included 17 species classified as LC, 27 NT, 47 VU, 15 EN, and 8 CR. Imputed statuses showed a similar distribution of ED to known species (Fig. S14). Qualitatively, the imputed species were heavily concentrated in tropical pleurodiran lineages, particularly those with many recently-described or resurrected species such as *Pelusios* (83) and *Pelomedusa* (84). Nearly all of the primarily sub-Saharan African pelomedusines, and ~2/3 of the primarily Australasian chelodinines were imputed. No other subfamily with more than 2 species had fewer than 50% assessments. The final breakdown of assessed plus estimated threat statuses used for subsequent analyses was 71 LC, 61 NT, 119 VU, 63 EN, 61 CR, and 9 EX. Thus, 66% of modern turtle and crocodilian species are threatened or extinct.

Comparing ED across threat statuses (Fig. 3), non-threatened species (LC, NT) have significantly lower median ED (18Ma vs. 21Ma) than threatened species (VU, EN, CR), using a two-sample *t*-test (*t*=−2.2, *P*=0.03). This is driven primarily by the higher median ED of VU and CR taxa, while EN covers a broad range of high- and low-ED species (Fig. 2). Thus, imminent extinction of threatened species would represent a disproportionate loss of total evolutionary history across crocodilians and turtles, preferentially removing unique and derived lineages from the Tree of Life, indicating their more-precarious stature in the conservation landscape compared to other terrestrial vertebrates such as amphibians

**Figure 3.**
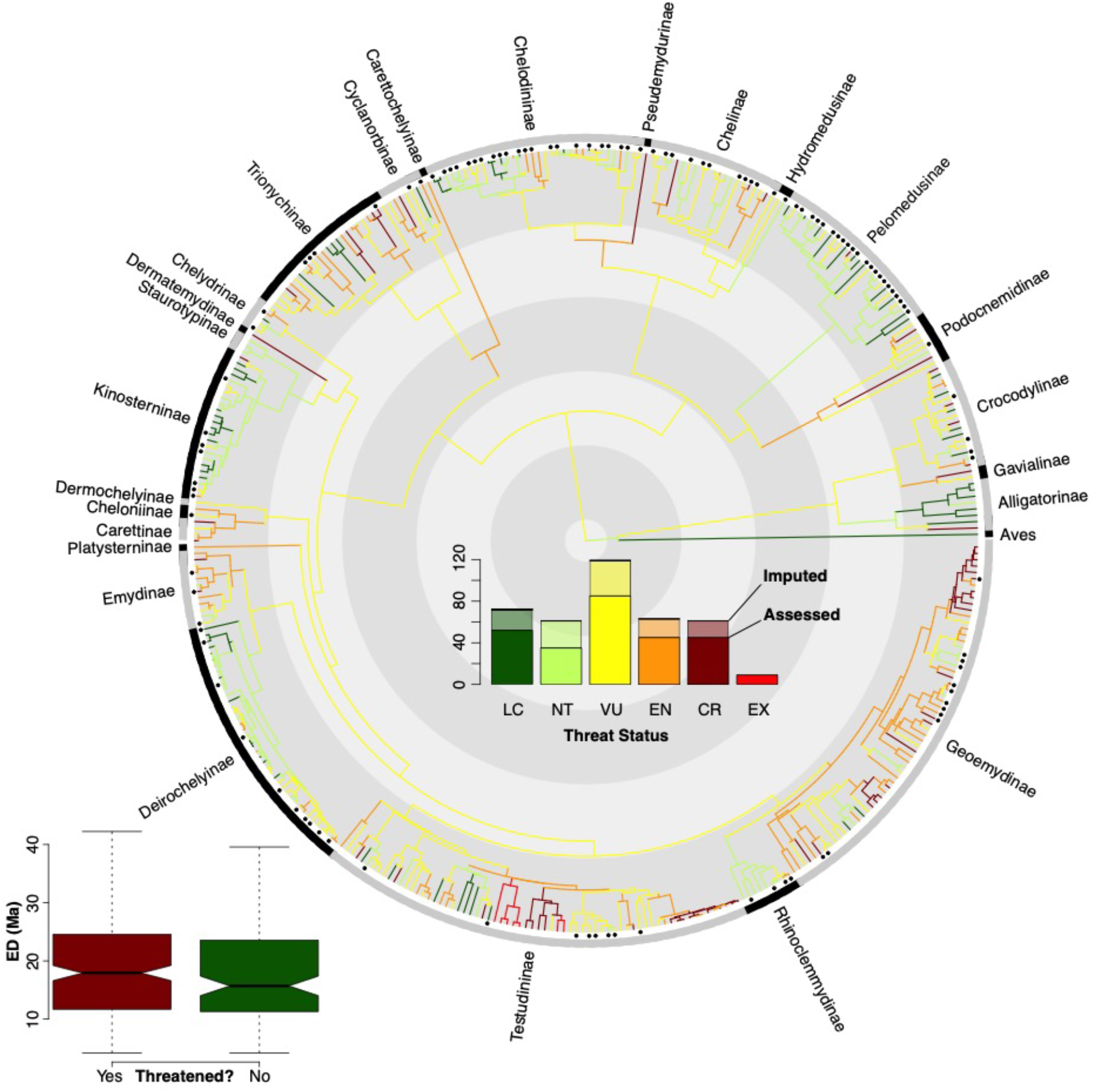
Randomly-selected phylogeny, with branches colored proportional threat status for all species; imputed statuses are indicated with a tip marker. The internal contrasting rings denote 50Ma intervals. The insets show the overall distribution of threat statuses (assessed are solid and imputed are transparent), and the distribution of ED across threatened vs. non-threatened, including both assessed and imputed species.

Combining the ED and threat statuses to calculate EDGE scores indicating the intersection of distinctiveness and risk, we highlight the top-10 at-risk turtles and crocodilians (Fig. 4; Table S1). These data also indicate that the highest-ranked turtles exceed crocodilians in their combination of ED and threat. We find that our estimated ED scores are similar to those from (4) when estimated directly from their phylogenetic dataset, but that a large cluster of their imputed values appear to be inflated (Fig. S15). Estimated EDGE scores are more similar between the two datasets, but variation in ED led previous authors to overemphasize some taxa (such as sea turtles; Cheloninae) and underemphasize others (such as flapshell turtles; Cyclanorbinae). Our dataset thus provides a more robust and presumably more accurate picture of ED and EDGE in the group (see Table S1).

**Figure 4.**
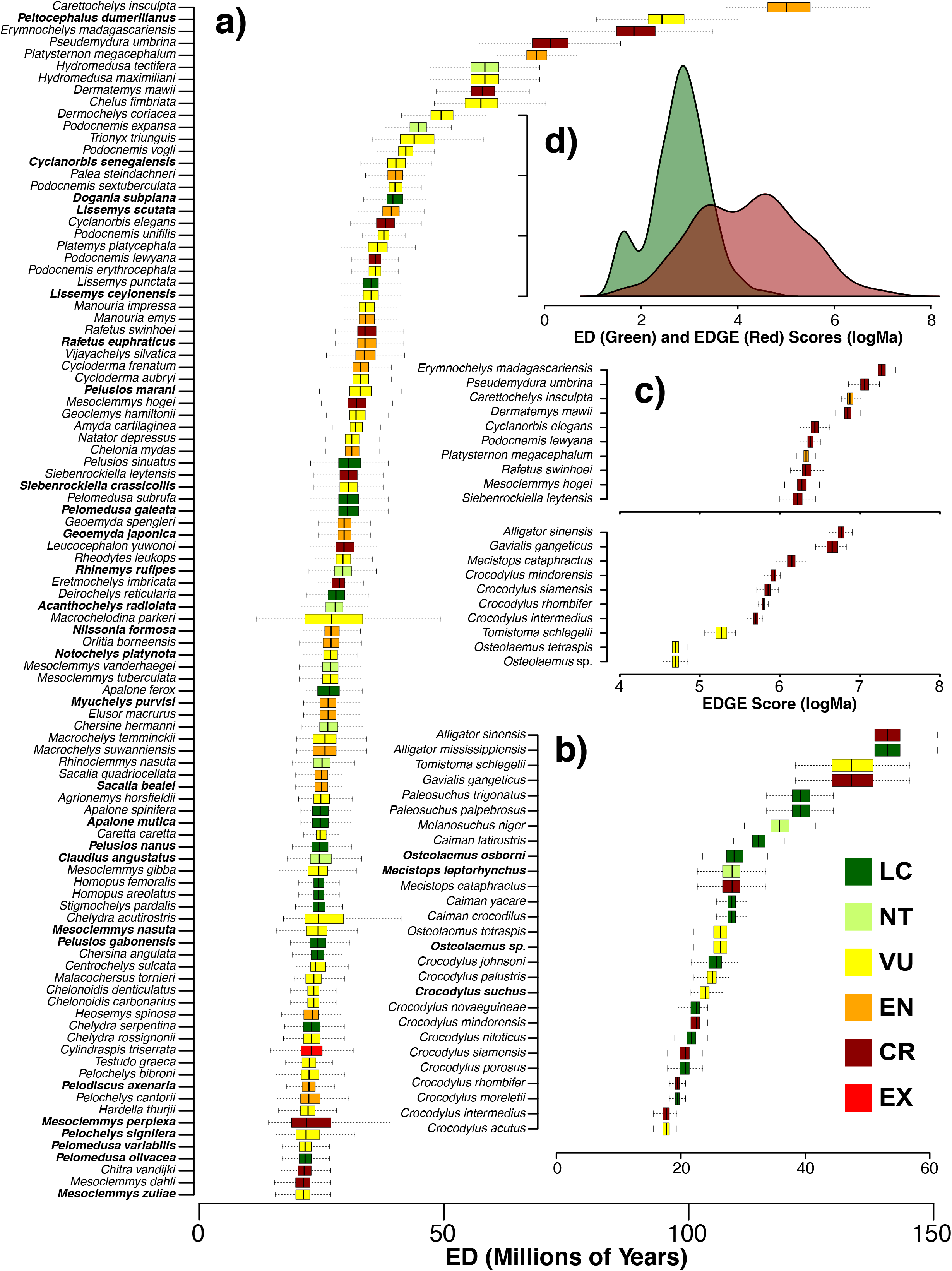
Boxplot of the top-100 ED turtles (a) and all crocodilians (b) with imputed statuses in bold, and insets showing boxplots of the top-10 EDGE species from both groups (c) and the divergence between ED (green) and EDGE (red) scores on a log scale (d). If all threat statuses were LC, ED and EDGE would be equal and the histograms would be identical. Their divergence thus shows the impact of extinction risk.

### Geography of threat

Geographically, threatened taxa are concentrated in a few major sub-areas (35) of the 11 global ecoregions (44). The highest diversity of threatened species occurs in South and Southeast Asia in the Red, Mekong, Irawaddy, and Ganges-Brahmaputra River deltas and Peninsular Malaysia (Fig. 5a). Secondary hotspots occur in tropical western Africa, and the eastern Amazon River basin. In contrast, “arks” of non-threatened species are observed in the eastern Nearctic, Australia-New Guinea, eastern Africa, and the western Amazon River basin (Fig. 5b). We note that the prominence of the eastern Nearctic is reduced when species richness is weighted by EDGE scores (Fig. S4), accounting for the low distinctiveness of the high number of questionably-delimited deirochelyine species in that area. Interestingly, there are also modest concentrations of non-threatened species (~5-10) in the Ganges-Brahmaputra River delta and the Yucatan peninsula, both regions characterized by several high-ED threatened species (Figs. 4, 5). Finally, island endemicity is not a strong predictor of threat status from any of the models, and regions such as Madagascar and the Philippines that have flagship high-ED threatened species (Table S1) nevertheless don’t represent concentrated hotspots of threat.

**Figure 5.**
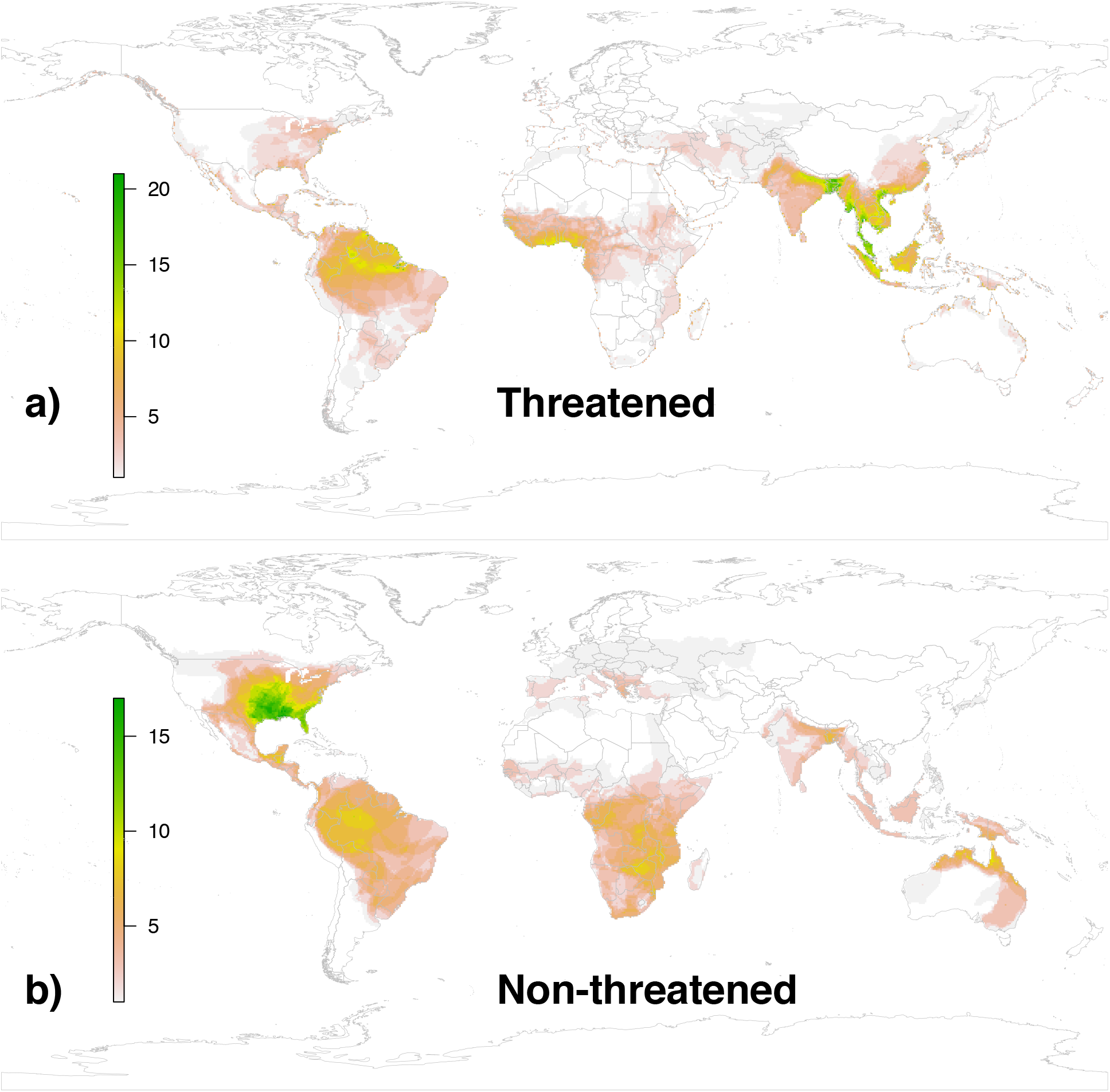
Spatial richness map showing distribution of species assessed or predicted to be threatened (a) or non-threatened (b). Note threat hotspots in the eastern Amazon, west Africa, south Asia, and southeast Asia; compared to relative havens in the eastern Nearctic, western Amazon, eastern Africa, and northern Australia.

## Discussion

### Phylogeny and diversification

Our topological results are closely aligned to previous work on turtles and crocodilians (15, 44, 45, 74–76) but offer a new time-calibrated perspective on the tempo and mode of evolutionary diversification and the phylogenetic and spatial distribution of extinction risk (Figs. 2–5). Consistent with their evolutionary durability and morphological conservatism in the fossil record (7–12), both turtles and crocodilians exhibit roughly constant rates of speciation and extinction across time, with no evidence for any significant mass-extinction events (Fig. 1a). This is unlike birds (25), mammals (79), amphibians (28), and squamate reptiles (85), which show strong evidence for shifts in diversification rate or mass-extinction events coincident with the K-Pg boundary. Similarly, turtles and crocodilians are defined by a single-rate background Brownian motion process of body-size evolution, with little evidence for significant among-lineage variation in rates, a few singleton species excepted (Fig. 1b).

Our estimate of turtle and crocodilian phylogeny is the most complete to date, with 95% of extant species sampled for up to ~21kb (Fig. 2). However, it is far from the last word on the subject, and we suspect that our (and previous) results are beset by confounding factors such as voucher identification (75), poorly defined species boundaries (86, 87), proper identification of targeted sequencing regions (88, 89), and mito-nuclear discordance (90, 91). We attempted to alleviate these issues by selecting well-characterized voucher specimens, but errors may still remain. We are generally confident in strongly supported relationships to the genus level, but caution that some species-level relationships may reflect the problems listed above. Similarly, as only 17 species required imputation, strong support for the monophyly of most genera provides at least preliminary confidence for the placement of imputed species. Overall, we consider the general concordance of our results with previous estimates to provide a sufficient basis for broad-scale analysis of global patterns, particularly with respect to biogeography, diversification, macroecology, and spatial distributions of extinction risk.

However, most turtle genera have not yet been subject to modern systematic revisions in an integrative taxonomic framework that combines morphological and molecular data with explicit criteria for delimiting species (92, 93). For instance, (87) showed that the eight traditionally-recognized species of *Pseudemys* reduced to only three species-level lineages in their molecular analyses. Similarly, (94) showed that the 14 currently-recognized species of *Graptemys*, long noted for their low levels of molecular divergence (95), appeared to represent eight or nine species-level lineages at most. These findings were confirmed (96) by authors who nevertheless continued to recognize 14 species. Similar instances of mismatches in the level of mitochondrial, nuclear, and morphological divergence have been noted in *Trachemys* (97) and *Cuora* (86, 89). In the Galapagos, nearly every island has a “species” of *Chelonoidis* that can putatively be diagnosed morphologically, but these have exceptionally low genetic divergence, and often interbreed and produce viable hybrids (98, 99). Many other apparently valid species of turtle (e.g., *Cuora*) also appear to hybridize naturally, in the wild, with great frequency (100, 101). Drastically reducing the number of recognized turtle species as indicated by molecular data in many lineages would likely decrease rate estimates and the prominence of clades such as *Chelonoidis* and Deirochelyinae in phylogenetic comparative analyses.

### Threat Status

In contrast to diversification rates and body size, ED and threat status show strong associations and non-random patterns in their evolutionary distribution (Figs. 2–4). Groups such as Alligatorinae (alligators and caimans), Podocnemidinae (South American river turtles), and Cyclanorbinae (flapshell turtles) are generally comprised of high-ED (>50Ma) species, while Kinosterninae (mud turtles) and Deirocheylinae (map turtles and sliders) consist almost entirely of low-ED (<17Ma) species. With regard to threat, groups such as Geoemydinae (Asian box turtles) and Testudininae (tortoises) are almost all threatened, while Pelomedusinae (African side-necked turtles) along with Kinosterninae and Deirochelyinae have a low proportion of threatened species. This pattern also yields a significant association between ED and extinction risk (Figs. 2, 3); loss of currently-threatened taxa would represent a disproportionate loss of evolutionary history in the group, far more so than in any other terrestrial vertebrate lineage.

A majority (66%) of turtle and crocodilian species are threatened, resulting from a number of factors including organismal traits and anthropogenic factors. By far the most important variable affecting extinction risk is the total geographic area covered by a species’ range, as in amphibians (17), birds (80), some mammals (81), and squamate reptiles (21). The effect of small ranges on extinction risk is due in large part to smaller population sizes (102, 103) and is likely amplified in turtles and crocodilians by their conspicuous nature (104), long generation times (105), and high degree of human exploitation (106, 107). For instance, amphibians and squamate reptiles with small ranges also tend to be threatened, but primarily due to ecological and demographic effects (17, 21). For squamates, human persecution or exploitation for food or commerce is generally limited to a few larger species (e.g., South American tegus and African bullfrogs) and not a common threat for the majority of taxa.

An additional critical factor facing turtles and crocodilians is the spatial distribution of their diversity; perhaps more so than for any other group of terrestrial vertebrates, “geography is destiny.” Birds, mammals, amphibians, and squamate reptiles exhibit massive species-richness in sparsely-populated tropical areas such as the Amazon and Congo river basins, the Andes mountains, and the Greater Sunda Islands (61). In contrast, while there are some turtle and crocodilian species in those areas, their peak species-richness (Fig. 5) is coincident with the highest population-density and strongest exploitation pressures on the planet, in South and Southeast Asia (37). Concomitantly, our PGLM model identifies occurrence in the Oriental ecoregion (39) as the second-strongest predictor of threat status behind area. Similar effects are seen in other heavily deforested and densely populated tropical regions such as western Africa and the eastern Amazon river basin (108). In contrast, areas such as Australo-Papua, eastern Africa, the western Amazon, and the Nearctic have lower population densities and less deforestation (46) and exhibit higher richness of non-threatened species (Fig. 5b).

A final pattern of great importance is that, despite the high concentration of threat in particular geographic regions such as the Indo-Gangetic plain (37), the distribution of species with the greatest combination of ED and threat (so-called ‘EDGE’ species (4)) is far more dispersed and idiosyncratic (Fig. S4, Table S1). This reflects the ancient history of both groups, across numerous tropical and temperate landmasses and remote oceanic islands (44). The top-10 turtle taxa by EDGE score include species from Madagascar (*Erymnochelys*), Papua New Guinea (*Carettochelys*), and the Philippines (*Siebenrockiella*), along with Africa, Asia, Australia, and South America. Similarly, the top-10 crocodilians by EDGE score are found in South and Southeast Asia, Africa, Indonesia and the Philippines, the Caribbean, and South America. Thus, while the bulk of diversity is concentrated and threatened in particular geographic hotspots (such as South and Southeast Asia), exceptionally distinct and threatened species are found outside of these areas in other global regions of high extinction risk.

Here, we provide a blueprint for guiding conservation management of these threatened taxa, highlighting both specific geographic areas with a high concentration of extinction risk, and exceptionally distinct and threatened species worldwide (Figs. 2–5). On the upside, given the relatively small number of species involved, their charisma, and the attendant amount of attention they receive, robust conservation-management plans and long-term restoration strategies are already in place for many turtles (6, 35) and crocodilians (6, 109). On the downside, while some of the “ark” areas such as the eastern Nearctic, Yucatan, and eastern Africa are improving in terms of human footprint (46), anthropogenic pressures show no signs of slowing or reversing in many of the global hotspots identified here such as South and Southeast Asia, western Africa, and tropical South America (110).

## Acknowledgments

This research was supported by US NSF grants DEB-1441737, DEB-1558568 and NSF DBI-1262600 to W.J. and DEB-1441719 to R.A.P., and by the GWU ColonialOne HPC Center.

