## Supplementary Information for "Phylogenetic and Spatial Distribution of Evolutionary Isolation and Threat in Turtles and Crocodilians (Non-Avian Archosauromorphs)"

**Supplemental Information – Colston et al.**

Here, we provide an outline of our dataset, analytical strategy, and detailed results. All data analyzed and all results obtained (and thus the full information needed to replicate this study) are provided either in this document, or in the supplemental files in DataDryad repository (to be deposited upon publication; contact TJC or RAP for data)..

*Taxonomy & Molecular Sampling*

For turtles, we used the 8th ed. of the Turtles of the World checklist from the IUCN Turtle Taxonomy Working Group (Rhodin et al. 2017), which recognizes 356 modern species in 94 genera in 14 families and 22 subfamilies. We also include *Aldabrachelys abrupta* and *Al. grandidieri,* which are distinct pre-modern species (Austin et al. 2003) but were omitted by the TTWG because they went extinct prior to modern times (~1Ka). However, in terms of geological time these clearly represent "modern" species and are thus retained here as "extant." In contrast, the species boundaries between *Amyda cartilaginea* and *Am. ornata* are poorly defined (see Rhodin et al. 2017), and we thus recognize only the former as a valid taxon encompassing all populations. This yields a total of 357 species in 94 genera, 22 subfamilies, and 14 families in our final classification.

Two new turtle species have been described since the publication of Rhodin et al. (2017). These are *Trachemys medemi* and *Kinosternon vogti* (see Uetz et al. 2018). These taxa were not included in our dataset and may represent extant populations (at least in part) of some species analyzed here. Future analyses should consider them separately; the models for threat status presented below can easily be used to predict their extinction risk based on known factors such as body mass and range size.

Crocodilian taxonomy is somewhat complex despite the low diversity, with at least 24 species in 9 genera in 4 families (see Oaks 2011). We follow Hekkala et al. (2011), Meredith et al. (2011) and Oaks (2011) in recognizing two species of *Crocodylus “niloticus,” C. niloticus* and *C. suchus.* Shirley et al. (2014) recognized two distinct species-level lineages of *Mecistops cataphractus*, the nominal species (Western lineage, type locality Senegal), and a “Central” species (from the Congo basin). Shirley et al. (2018) formally resurrected the name *M. leptorhynchus* for the latter, which we recognize here.

Finally, we follow Eaton et al. (2009), Oaks (2011), Franke et al. (2013), Shirley et al. (2014), and Shirley et al. (2015) in recognizing three species of *Osteolaemus*, *O. osborni* (Congo Basin), *O. tetraspis* (Ogooué Basin), and a third, as-yet unnamed species from West Africa. Franke et al. (2013) demonstrated phylogeographic structure in this West African form, but we follow them in not recognizing the two lineages as full species. For the West African form, the name *Halcrosia afzelli* Lilljeborg, 1867 with the type locality Sierra Leone appears to be available. Thus, for the sake of nomenclatural convenience in the dataset, we refer to the West African form as *O. afzelli*. However, we specifically disclaim the above act as not being a valid nomenclatural change under Article 8.2 of the ICZN, pending taxonomically complete re-description of these species. This yields a total of 27 crocodilian species.

Franke et al. (2013) demonstrated that the identity of captive specimens labeled *"Osteolaemus tetraspis"* varied widely, and all 4 specimens sampled by Oaks (2011) were captive. By comparing the CYTB datasets of Oaks (2011) and Franke et al. (2013), we made the preliminary determination that LSUMZ H-21755 corresponds to *O. tetraspis,* while 21756, 6990, and 6992 are actually *O. afzelli.* The *Osteolamus* sequenced by Gatesy et al. (2003) is also *O. afzelli,* based on 12S and CYTB affinities. Similarly, based on the data from Shirley et al. (2014) and their comments regarding the origin of captive U.S. *Mecistops,* the captive specimens LSUMZ H-6975, 21718, 21719, and 21720 sequenced by Oaks (2011) are the Western nominotypical lineage *M. cataphractus,* as is the individual sequenced by Gatesy et al. (2003).

Furthermore, we determined using the CYTB dataset that the GenBank entries for two complete mitochondrial genomes of *O. tetraspis* (EF551001 and NC_009728) both represent the nominotypical lineage, as do both *M. cataphractus* (EF551000 and NC_010639). We chose the latter in each case to fill in the remaining gaps in our matrix for those taxa. Finally, we were unable to determine the identity of the *Osteolaemus* specimens represented by Ornithine Decarboxylase (ODC) sequences in McAliley et al. (2006), and thus did not include them.

We initially used the alignment of Oaks (2011) as is, including 4 mitochondrial genes and 9 nuclear genes, GenBank accession nos. JF314862–JF315859. We trimmed the taxon sampling to a single terminal from each of the 25 species and excluded the tRNAs. As the matrix of Oaks (2011) is essentially 100% complete, we simply selected the first representative from each species as displayed in Figure 1b of Oaks (2011), starting from the top. We then gathered data from conspecific specimens for the remaining 15 genes. We assiduously determined the identity of each voucher specimen for *Mecistops* and *Osteolaemus* used for the alignments (as described above) based on their identification by Franke et al. (2013) and Shirley et al. (2014, 2015) for proper inclusion in the alignments. Similar care was taken for the recently recognized species *Crocodylus suchus,* which was formerly considered *C. niloticus,* though this was much simpler given their geographic distinctiveness.

We attempted to minimize chimerism by deriving the largest possible number of sequences from the same voucher specimen. Most species were represented by most if not all genes and loci; 21 species were represented in alignments for at least 23 of 27 genes. However, a few lesser-known caiman (*Caiman latirostris, C. yacare,* and *Melanosuchus niger*) were represented in only 14–16 of the 27 total alignments. Additionally, due to our reliance on only well-characterized vouchers, the species *Mecistops leptorhynchus* and *Osteolaemus osborni* were only represented in 6 and 5 alignments, respectively. Finally, *O. afzelli* was represented in 17 alignments, individuals of this species having been sequenced in several studies. Curiously, the data of Shirley et al. (2014) do not appear to have been uploaded to GenBank, only DataDryad repository https://doi.org/10.5061/dryad.sh3m0, and thus accession numbers are not available for these sequences. We used the voucher "CTang01" from Lake Tanganyika as our representative from *M. leptorhynchus*.

For turtles, significant amounts of data both molecular and mitochondrial are available for a significant proportion of extant species. These have been collated by previous researchers including Thomson and Shaffer (2010), Guillon et al. (2012), Rodrigues and Diniz-Filho (2016), and Pereira et al. (2017). Fortunately, these data overlap significantly in terms of locus sampling with the matrix of Oaks (2011). Thus, we sampled 14 mitochondrial genes for all turtles, crocodilians, an avian outgroup (*Gallus*), and a lepidosaurian outgroup (*Sphenodon*). These are 12S, 16S, ATP6, ATP8, COI, COII, COIII, CYTB, DLOOP, ND1, ND2, ND3, ND4, and ND5. For nuclear loci, we sampled BDNF, CMOS, GADPH, ODC, RAG1, and RAG2 for all taxa. We sampled R35 for turtles only, as the intron has not been widely sequenced in other groups. We omitted ACHR, ACTB, ACTC, ATROP, LDHA, LDHB, and RHO from the Oaks (2011) matrix, as they were not available from many turtle species. This yielded 7 total nuclear loci, 14 mitochondrial genes, and 21 genes overall. Sequence data were available for 340 of 357 turtles, all crocodilians, and the two outgroups. The matrix was 24,798 bp in total length, ranging from 405 to 20,929 bp per species, and averaging 8,891 bp for each of the 369 terminals. GenBank accession numbers are given in Dryad.

In general, protein-coding exons were aligned via translation in Geneious except ND3, which has a functional frameshift mutation in many turtle species (Russell and Beckenbach 2008). The control region/d-loop proved particularly difficult to align using traditional methods, though regions of homology were clearly present. Thus, we employed the PASTA algorithm (Mirarab et al. 2015), which uses an iterative strategy of tree-building, multiple alignment within subclades, and merging of subclade alignments to arrive at an optimal global alignment. We ran 20 iterations under the default settings, using FastTree (Price et al. 2010) and MAFFT (Katoh et al. 2002) for tree building, and alignment, respectively. Additional iterations did not result in improvement in alignment scores.

Two nuclear loci, GAPDH and ODC, consisted of both exons and introns, with the majority of informative sites occurring in the introns. These intronic regions showed very little subjective similarity between turtles and crocodilians when aligned, but high similarity within groups, and high similarity in the exons. We chose to retain them, as the resulting phylogenies showed good apparent resolution within each group. However, the alignments may reflect low homology among lineages across sites, potentially inflating estimated rates or branch lengths. We anticipate this effect to be minor, though, when integrated across the dataset under the relaxed-clock model. Future studies may wish to expand exon-based sampling for these loci to alleviate this potential issue.

*Phylogenetic Inference & Topology*

We developed an optimal partitioning strategy for both ML and BI to facilitate the downstream PASTIS analyses. Of the 21 genes, 14 could be modeled by codon position, while 7 were either structural (12S, 16S, DLOOP), primarily introns (GAPDH, ODC, R35), or had internal frame-shift mutations that rendered codon homology ambiguous (ND3). Thus, we had 49 potential partitions.

We used PartitionFinder 2 (Lanfear et al. 2016) to estimate the best partitioning strategy. For the RAxML analysis, we used the --raxml (Stamatakis 2006) and search = 'rcluster' (Lanfear et al. 2014) flags to decrease computational intensity, given the size of the matrix. We restricted the model set to GTR+G as we used RAxML version 8 (Stamatakis 2014), which only supports GTR and recommends using the GAMMA approximation for very large matrices. Thus, we were essentially addressing the number of distinct partitions less than or equal to 49 that needed to be modeled using GTRGAMMA in RAxML. To determine the optimal BI models for MrBayes, we ran a similar analysis using the the 'mrbayes' model set.

The RAxML-specific analysis yielded 41 distinct partitions using the GTRGAMMA model, and the MrBayes-specific analysis yielded 42 partitions taking the GTR, GTR+G, or GTR+I+G models. Use of the 'rcluster' algorithm, necessitated by the number of partitions, requires the use of RAxML to calculate likelihoods and limits model selection to the GTR family. Given the degree and complexity of molecular divergence within and among lineages, it seems unlikely that simpler models would be needed.

For initial inference of ML topology, we used RAxML version 8.2.9, with the rapid-bootstrapping option ("-f a"), 1000 non-parametric bootstrap replicates, and a final round of ML searches starting from every $5^{th}$ bootstrap replicate, for a total of 200 indepdent ML estimates to find a globally optimum result. We used the partitioning strategy defined above from Partition Finder 2, applying the GTRGAMMA model to the optimized character-sets. This analysis included the 340 turtles, 27 crocodilians, one bird (*Gallus*), and a lepidosaurian (*Sphenodon*) as the outgroup. The topology with support values is given in Dryad. We used this topology as the primary constraint for the PASTIS-based MrBayes analysis, which simultaneously imputes the missing species into the phylogeny using taxonomic constraints (see below) and estimates divergence times using temporal constraints.

Our estimate of turtle phylogeny is the most complete to date (Appendix S1), with 95% of extant species sampled for up to ~21kb. However, it is far from the last word on the subject, and we suspect that our current results (as well as those of previous researchers using similarly constructed datasets) are beset by numerous issues such as voucher identification (see Guillon et al. 2012), poorly defined species boundaries (Stuart and Parham 2004; Spinks et al. 2013), proper identification of targeted sequencing regions (Spinks and Shaffer 2007; Fritz et al. 2010), and nuclear-mitochondrial discordance (Wiens et al. 2010; Spinks et al. 2015). We attempted to alleviate these issues to the greatest degree possible by selecting well-characterized voucher specimens, but many issues likely still remain.

We are generally confident in strongly supported relationships to the genus level, but caution that some species-level relationships may reflect the issues listed above. Similarly, as only 17 species required imputation, the strongly supported monophyly of most affected genera (see below) provides at least preliminary confidence in their low-level placement. Regardless, we consider the general concordance of our results with previous estimates to provide a sufficient basis for broad-scale analysis of global patterns, particularly with respect to biogeography, diversification, macroecological associations, and spatial distributions of threat and risk.

Generally speaking, our results are highly concordant with those of most recent crocodilian (e.g., Oaks 2011) and turtle (e.g., Guillon et al. 2012; Rodrigues and Diniz-Filho 2016; Pereira et al. 2017) studies, as expected since they are based on the same or highly-similar underlying data. As stated above, our tree thus reflects several notable inconsistencies or features that influence either taxonomic placement (e.g., for PASTIS), or comparative analyses (e.g., BAMM). We review these here as a guide both to interpreting our downstream results, and to highlight areas of future research.

Overall, the backbone and most relationships at the genus and species level are strongly supported (e.g., >90% BS). Similarly, the vast majority of genera are estimated to be monophyletic, and the tree is almost fully-sampled (367/384 turtles and crocodilians). Of particular note is the continued weak support for relationships among tortoise genera (Testudinidae), as well as broader uncertainty in higher-level relationships among and placement of major groups such as Cheloniidae, Chelydridae, and Platysternidae (see Thomson and Shaffer 2010; Guillon et al. 2012; Rodrigues and Diniz-Filho 2016; Pereira et al. 2017). We recover a unique topology from these previous four analyses, with successive divergences of Chelydridae+Dermatemydidae+Kinosternidae, Cheloniidae+Dermochelyidae, Platysternidae, Emydidae, and Testudinidae+Geoemydidae. However, this is based on similar underlying datasets, and likely reflects issues such as mito-nuclear discordance and weak phylogenetic signal in the available loci, rather than an increase in phylogenetic accuracy over any previous study. Hopefully, future analyses sampling more characters will be able to resolve these relationships with greater support.

The genus *Mesoclemmys* is paraphyletic with respect to *Phrynops, Acanthochelys, Platemys,* and *Rhinemys,* reflecting the results and data of Reid et al. (2011). The species *M. zuliae, M. dahli, M. nasuta,* and *M. gibba* form a strongly supported clade that is the sister lineage of the remaining taxa. The species *M. hogei* is strongly supported as the sister lineage of *Phrynops.* Finally, *M. tuberculata* + *M. vanderhaegei* are weakly supported as the sister lineage of *Rhinemys,* and this group of three species forms a strongly supported clade with *M. raniceps* + *M. heliostemma.* The only species missing from our dataset, to be imputed using PASTIS, is *M. perplexa,* which the original description (Bour and Zaher 2005) refers to the clade including *M. gibba* and *M. nasuta;* the Amazonian species. As *M. gibba* is also the type species of the genus, we restrict the placement of *M. perplexa* to the clade formed by the species *M. zuliae, M. dahli, M. nasuta,* and *M. gibba,* including the stem. The species *P. tuberosus* also required imputation and was placed with the other strongly-supported members of this genus, including the stem.

Also, as in Reid et al. (2011) within Chelodininae, the subgenus *Macrochelodina* recognized by Rhodin et al. (2017) is paraphyletic with respect to *Macrodiremys* and *Chelodina.* The former is a monotypic subgenus recognized by Rhodin et al. (2017), while the sampled species of the latter are strongly supported as monophyletic. As there are three unsampled species each for both *Chelodina* and *Macrochelodina,* we chose to allocate all of them conservatively to the entire clade formed by *Macrochelodina, Macrodiremys,* and *Chelodina,* including the stem lineage. Most authorities consider these as a single genus (*Chelodina*) regardless, and it seems clear that the subgeneric taxonomy will need to be revised.

Additionally, within Chelodininae, Le et al. (2013) erected the genus *Flaviemys* for the former *Myuchelys purvisi.* Spinks et al. (2015) later showed that the paraphyly of *Myuchelys* stemmed from mito-nuclear discordance, and that nuclear data supported a monophyletic *Myuchelys.* As our dataset includes both mitochondrial and nuclear markers, we recover a paraphyletic *Myuchelys.* While this is likely an erroneous topology, it does not affect placement of the remaining unsampled species in Chelodininae. These are *Elseya flaviventralis* (monophyly of *Elseya* is strongly supported) and *Hanwarachelys rhodini* and *H. schultzei,* which we restrict to the single branch subtending *H. novaeguineae,* the only species of *Hanwarachelys* sampled in our dataset. The subgenus *Pelocomastes* is paraphyletic (as in Rodrigues and Diniz-Filho 2016; Pereira et al. 2017), with *P. albagula* forming the sister lineage to the remaining *Pelocomastes, Hanwarachelys,* and *Elseya*. This does not affect the placement of any missing taxa.

For the remaining unsampled species in the tree, all were members of genera that are represented by multiple sampled species, and are estimated to be monophyletic, easing consideration of their placement for PASTIS. These were *Chelydra acutirostris, Cuora cyclornata, Aldabrachelys abrupta, Chelonoidis guntheri, Chelo. microphyes,* and *Pelochelys signifera.* These species were all constrained to occur with their respective congeners, including the stem lineage of the genus. The two *Chelonoidis* species were limited to the Galapagos lineage, excluding the continental South American species.

Issues of mito-nuclear discordance, such as those affecting *Myuchelys,* have also been observed in New World emydids (see Wiens et al. 2010), but these do not seem to be affecting generic monophyly in the current dataset, possibly due to the increased volume of characters from nuclear loci. Thus, *Trachemys nebulosa* is recovered within a monophyletic *Trachemys.* In Kinosternidae, we recover a monophyletic *Kinosternon* (like Spinks et al. 2014), contrary to Iverson et al. (2013) who found *Sternotherus* nested within *Kinosternon* and erected a new genus *Cryptochelys* for *K. acutum, K. angustipons, K. dunni, K. creaseri, K. herrerai,* and *K. leucostomum*. These alternative topologies also appear to stem from mito-nuclear discordance, though not in all mitochondrial loci (see Spinks et al. 2014).

Several clades have been identified by previous authors (see Rodrigues and Diniz-Filho 2016) as representing extraordinary instances of evolutionary diversification, including Galapagos tortoises (*Chelonoidis*), and New World emydids (Deirochelyinae). Similar results are recovered in our analyses (see below). However, we strongly believe that these are artifactual, and represent an inconsistent application of species concepts and species delimitation in turtles. This is further confounded by inconsistent application of taxonomic ranks such as subgenera and subspecies. This is not a criticism of any researcher, but the product of historical biases and inertia in systematic practices.

For instance, Spinks et al. (2013) showed that the seven to nine traditionally-recognized species of *Pseudemys* (we recognize eight here) reduced to only three species-level lineages in their molecular analyses. Similarly, Praschag et al. (2017) showed that the 14 currently-recognized species of *Graptemys,* long noted for their low levels of molecular divergence (see Lamb et al. 1994), appeared to represent eight or nine species-level lineages at most. Thomson et al. (2018) generally confirmed these findings but re-affirmed the geographic and morphological distinctiveness of the traditionally-recognized taxa. Similar instances of apparent mismatches in the level of mitochondrial, nuclear, and morphological divergence have been noted in *Trachemys* (Fritz et al. 2011), as well as other genera such as *Cuora* (Stuart and Parham 2004; Spinks and Shaffer 2007). In the Galapagos, nearly every island has a “species” of *Chelonoidis* that can be diagnosed morphologically (at least in theory), but these have exceptionally low genetic divergence, and many freely interbreed and produce viable hybrids (see Edwards et al. 2014; Poulakakis et al. 2015). Many other apparently valid species of turtle (e.g., Cuora) also appear to hybridize naturally, in the wild, with great frequency (see Parham et al. 2001; Strujik 2016).

In short, most turtle groups have not yet been subject to modern systematic revisions in an integrative taxonomic framework that combines morphological and molecular data with explicit criteria for delimiting species (see Fujita et al. 2012; Solis-Lemus et al. 2015). Indeed, many turtle lineages do not appear to conform to many of the commonly-used operational or theoretical criteria of biologically-oriented species concepts (such as consistent reproductive isolation with concordant morphological and molecular divergence) that we might like to apply in such revisions. This is not to say that these clades do not exhibit the remarkably dramatic and rapid ecomorphological diversification.

However, it is clear that what we diagnose, define, and delimit as “species” clearly varies across the Turtle Tree of Life, in a way that confounds species delimitation and accurate phylogenetic inference, at least for a bifurcating tree. Drastically reducing the number of species, as apparently indicated by molecular data in many lineages, would likely decrease rate estimates, and the prominence of clades such as *Chelonoidis* and Deirochelyinae in phylogenetic analyses of diversification rate. At the current juncture, we are constrained by existing taxonomic and phylogenetic frameworks, and these issues must thus be left to future researchers. However, interpretation of our downstream comparative results (see below) should be colored by knowledge of these issues. We attempt, when possible, to use analytical strategies and interpret statistical models in a way that is agnostic to the possibility of low-level taxonomic biases.

*Divergence-Time Estimation and Taxonomic Imputation*

Three decades of molecular divergence-time estimation has resulted in a large dataset of estimated node-ages, concordance among which now provides a convenient meta-prior on dates for major line-ages in the Tree of Life (Hedges et al. 2006). These reflect a wide variety of analytical methods and un-derlying fossil data, and thus integrate over uncertainty in variables such as molecular rates, fossil rela-tionships, and character and taxon sampling. Thus, rather than generate our own idiosyncratic estimate of proper node-age calibrations, we choose to leverage these existing data to parameterize node-ages within appropriate bounds that are congruent with the bulk of existing datasets. The website Timetree: The Timescale of Life (http://www.timetree.org/; accessed 1 June 2018) aggregates these data conven-iently, providing a summary of the published literature. For each node of interest, we queried Timetree to gather the studies addressing this node, their estimated ages, and the Timetree consensus age.

Similar to previous studies (Tonini et al. 2016; Jetz and Pyron 2018), we placed minimum and maximum ages on seven key higher-level nodes, using uniform densities. The first is the root node Sauria, subtend-ing *Sphenodon* and *Gallus.* Time tree records at least 24 studies calculating the age of this node, ranging from 245-300Ma. The estimated age is 280Ma, with a range from 273-286Ma, which we used as the cali-bration. The second is Archosauromorpha, subtending *Gallus* and *Cuora,* for which 22 studies range from 152-300Ma. The estimated age is 254Ma, with a range from 240-268Ma. The third is Archosauria, subtending *Gallus* and *Alligator,* for which 28 studies range from 183-259MA. The estimated age is 237Ma, with a range from 229-244Ma. The fourth is Crocodylia, subtending *Alligator* and *Crocodylus,* for which 15 studies range from 25-151Ma. The estimated age is 80Ma, with a range from 63-98Ma. The fifth is Testudines, subtending *Peltocephalus* and *Cuora,* for which 19 studies range from 88-215Ma. The estimated age is 184Ma, with a range from 161-206Ma. The sixth is Pleurodira, subtending *Peltocephalus* and *Emydura,* for which 10 studies range from 52-177Ma. The estimated age is 132Ma, with a range from 103-160Ma.The seventh is Cryptodira, subtending *Carettochelys* and *Cuora*, for which 15 studies range from 97-250Ma. The estimated age is 172Ma, with a range from 138-185Ma.

To impute the missing taxa, we used the PASTIS approach (Thomas et al. 2013), which implements the relatively simple strategy of constraining the topology for species with sequence data (we used the ML topology from RAxML), and allowing the placement of unsampled species to vary, with branch lengths drawn from a birth-death process. This method has been used previously for birds (Jetz et al. 2012), squamates (Tonini et al. 2016), and amphibians (Jetz and Pyron 2018), and we generally follow their recommendations. We produced the constraint file using PASTIS 0.1-2, based on the ML topology and the hypothesized placement of the 17 missing species as described above. For analysis in MrBayes, we used the 42-partition scheme estimated by Partition Finder 2, and the temporal constraints described above. For the branch lengths, we used the recommended default priors of zero extinction and an exp(1) prior on speciation, for a conservative approach to estimating tree shape (see Jetz et al. 2012).

To estimate prior values for the independent branch-rates model, we followed the recommendations of Ronquist et al. (2012) and conducted two runs of 13 million generations (for 10 million post burn-in), one under a strict clock, and one with branch lengths measured in substitutions per site. These quickly reached stationarity with ESS>200 for all or nearly all parameters. The slope *b* of the relationship between the variance of the clock-like and non-clock-like branches (0.01439174) was used as the exponential mean for igrvarpr, 1/*b**log(2) = 48.16. Using a similar approach to Ronquist et al. (2012), the median tree height from the strict clock analysis (0.9006) divided by the mean estimated root age (279.5Ma) gives an estimate of the mean clock-rate, $m$ = 0.00322 subst./site/Ma. An approximation of the standard deviation is given by the upper range of the 95% Highest Posterior Density of tree height (0.9284) divided by the minimum root-age (273Ma) = 0.0034 subst./site/Ma minus $m$, for an estimate of sqrt($v$) = 0.00018. These were then converted to log space, with the lognormal mean equal to $\ln(m/\mathrm{sqrt}(1+(v/m^2)))$ = -5.74, and lognormal standard deviation equal to $\mathrm{sqrt}(\ln(1+(v/m^2)))$ = 0.06.

Combined, this yields a MrBayes input file that accomplishes three major goals simultaneously. First, the placement of the 17 missing taxa is imputed according to our best estimate of overall topology for the sampled species and previous taxonomic classifications. Second, branch-lengths are re-estimated according to the best-estimate partitioning scheme from the sampled loci. Third, branch lengths are estimated proportional to time under a birth-death process parameterized from the fossil-record and previous divergence-time estimates, as well as the observed substitution process in the molecular sequence-data gathered here. We ran 8 runs of 4 chains for 55 million generations, sampled every 10,000$^{th}$.

Of these, two runs did not achieve convergence and were discarded. For the remaining six runs, we increased burnin by one-million-generation increments until the log-likelihoods exhibited ESS>100. The post burnin results were then combined for a total of 275 million post-burnin generations. All parameters except tree height and clock rate had ESS>100 in the combined results. We hypothesize that the combination of a highly parameterized by sparsely sampled supermatrix with strict priors on both topology and dates resulted in low posterior signal and difficulty achieving convergence. However, given the stringency of the taxonomic and temporal constraints, we anticipate this having a very minor impact on the estimated branch lengths or imputed position of missing taxa.

From the 275 million post-burnin generations, we randomly sampled a posterior distribution of 10,000 taxonomically-complete, dated phylogenies estimated using multiple independent lines of evidence to approximate the globally-optimal tree. While some authors caution against using phylogenies with imputed taxa to measure rates of character evolution (Rabosky 2015), we note that i) the number of imputed taxa is extremely small, which should reduce any bias in estimated rates; and ii) those authors also found that estimated diversification rates did not suffer from obvious bias (Title and Rabosky 2016). These phylogenies can thus be used for myriad downstream comparative analyses, such as estimating diversification regimes and biogeographic reconstructions. However, we temper this with mindfulness of the potential non-equivalence of taxonomic units at the species level, as described above.

*Trait Data*

For the purposes of analyzing morphological evolution and ecological correlates of extinction risk, we gathered several traits shown to be relevant in other terrestrial vertebrate groups such as birds, mammals, amphibians, and reptiles (Davidson et al. 2009; Bland et al. 2014, 2015; Jetz and Freckleton 2015). Specifically, we measured body length (carapace length [CL] or total length [TL] in cm), body mass (g), microhabitat (freshwater, terrestrial, or marine), environment (aquatic, semi-aquatic, or terrestrial), endemicity (island or mainland), biogeographic region (11 ecoregions), range size ($km^2$), human-encroachment index (proportion of urban or cropland in range), and spatial proximity (distance of range centroids). For more detail on the spatial data, see the next section. All data and sources are given in the supplement and Dryad.

For body length, we used existing datasets (e.g., Itescu et al. 2014) and a comprehensive literature search to obtain maximum CL or TL values for all 357 turtle species and 26/27 crocodilians (Fig. S1). A reliable maximum length was unavailable for *Osteolaemus afzelli.* For body mass, we built on datasets such as Regis and Meik (2017) to obtain maximum mass for 202 turtle species and 23 crocodilians (Fig. S2). For habitat and endemicity, we followed Jaffe et al. (2011) for the initial classification scheme, and categorized all remaining species using the range maps in Rhodin et al. (2017) and additional sources.

To ensure comprehensive coverage of trait data for predicting threat status, we imputed the missing values of length (for *Osteolaemus afzelli;* Appendix S2) and body mass (for 159 species; Appendix S3) using phylogenetic methods accounting for trait covariance (Goolsby et al. 2017). Specifically, we used Rphylopars, incorporating tree structure and the data for length and mass. Using the default algorithm, we imputed the missing values across 10,000 trees, used the mean as our best estimate (Fig. S3).

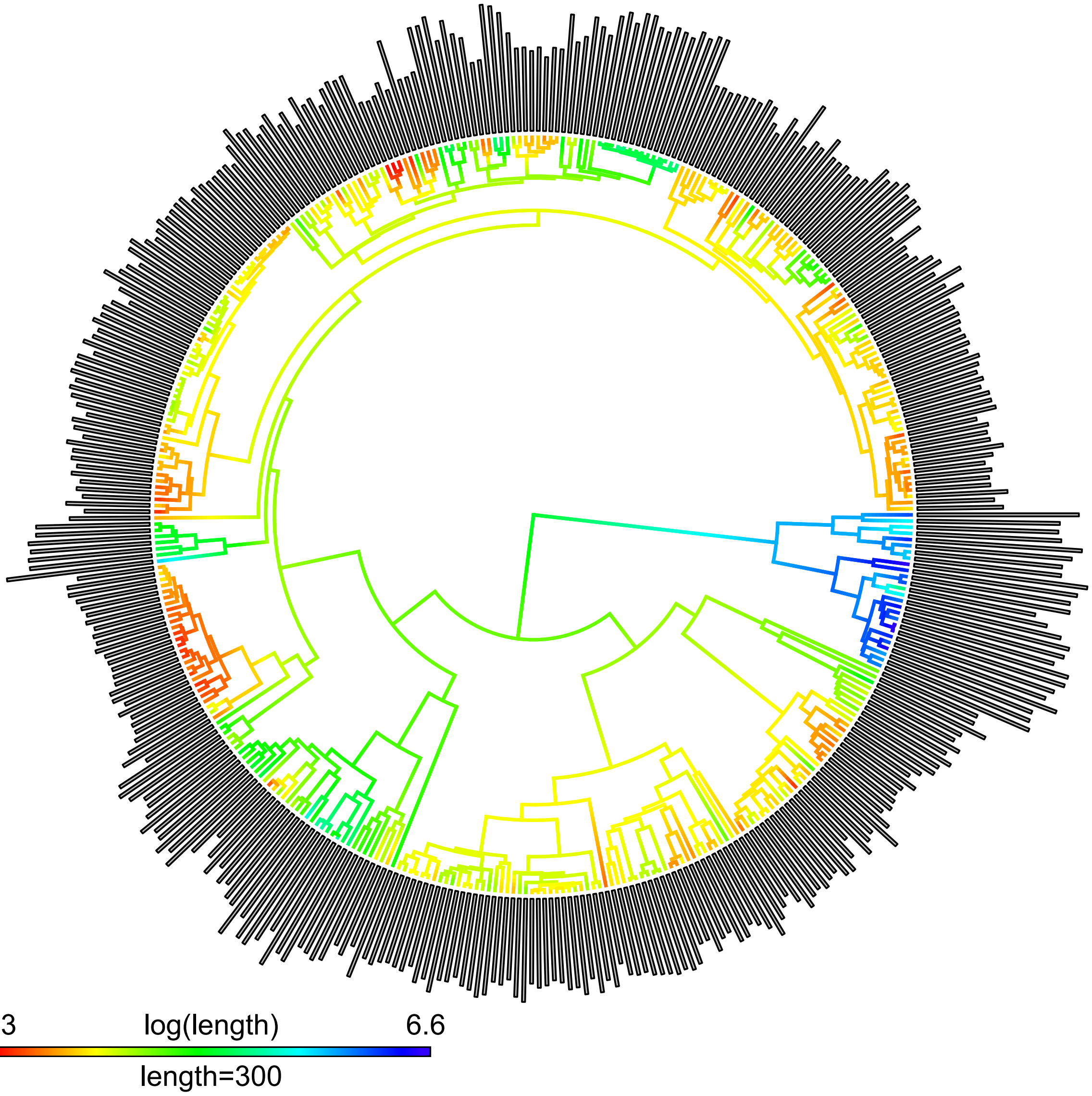


**Fig. S1.** Ancestral-state reconstruction and trait values for total length in cm.

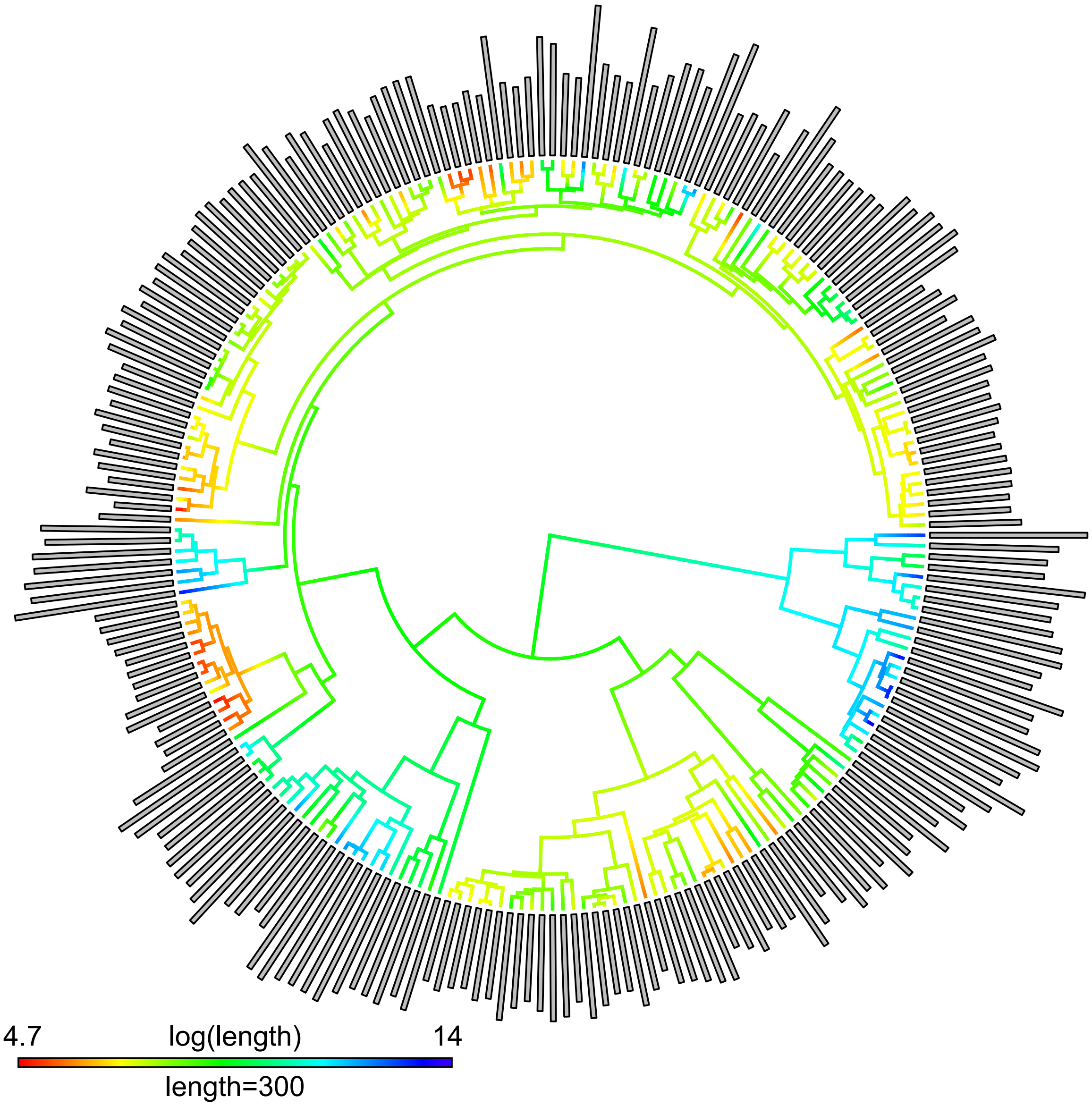


**Fig. S2.** Ancestral-state reconstruction and trait values for body mass in g.

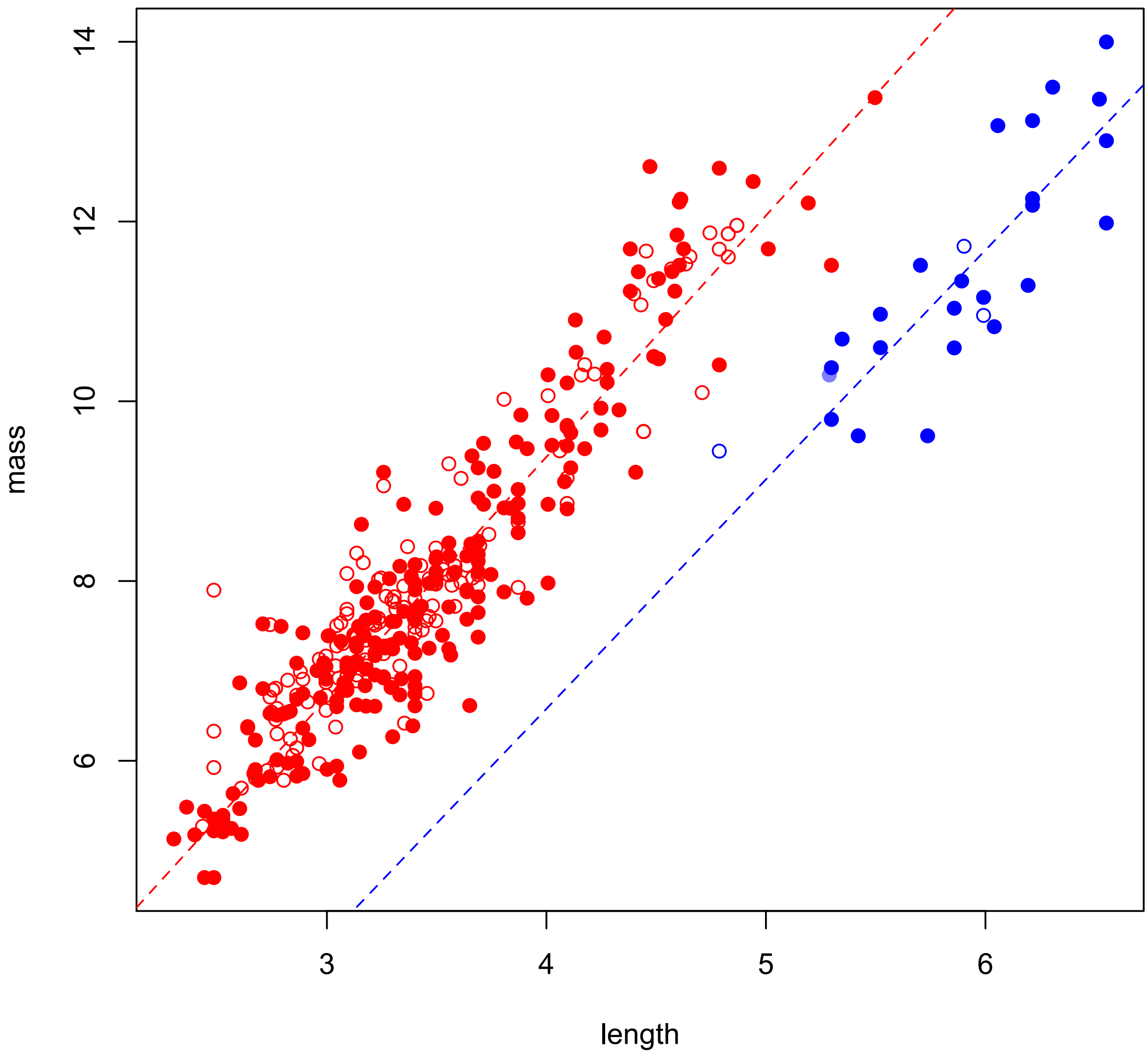


**Fig. S3.** Linear relationship between log(length) and log(mass) for turtles (red) and crocodilians (blue). Filled circles are the empirical data (graphed in Figures SX and SX), open circles are the mean of the 10,000 imputed values of mass for the missing species, and the transparent blue circle is the mean of the 10,000 imputed values of length and mass for *Osteolaemus afzelli*.

*Spatial Data*

Due to biogeographic history and differences in anthropogenic pressure, the distribution of phylogenetic diversity and extinction risk tend to vary drastically among regions (e.g., Jetz et al. 2014; Rosauer et al. 2017). For biogeographic area, we followed Pereira et al. (2017) in classifying species into one or more of the following areas based on range maps in Rhodin et al. (2017): Afrotropical (AF), Australia (AU), Eastern Palearctic (EP), Indian Subcontinent (IS), Madagascar (MG), Nearctic (NA), Neotropical Central America (NCA), Neotropical South America (NSA), Oriental (OR), West Indies (WI), and Western Palearctic (WP). These areas are broadly concordant both with global patterns of turtle endemicity, as well as other terrestrial vertebrate groups (e.g., Pyron 2014).

For range size, we began with the dataset of Roll et al. (2017), which provides global range maps for reptiles and area ($km^2$) calculated from a Berhmann equal-area projection. Their data include polygons for 322 turtles and 24 crococodilians. To match this to our reference taxonomy, we merged *Trachemys grayi* and *T. emolli,* and replaced *Mecistops cataphractus, Osteolaemus tetraspis,* and *Pelomedusa subrufa* with redrawn maps for those species after their recent taxonomic subdivisions.

For the remaining species, we drew new maps projected to World Cylindrical Equal Area (EPSG:54034), with range data for land turtles from Rhodin et al. (2017); *Mecistops* from Shirley et al. (2018); *Osteolaemus* from Shirley et al. (2014); *Aldabrachelys* from Gerlach and Canning (1998); and sea turtles from the SWOT (Kot et al. 2015) and OBIS-SEAMAP (Halpin et al. 2009) datasets, specifically the regional management units (RMU) from Wallace et al. (2010), with the RMU files accessed from SWOT/OBIS-SEAMAP on 8 December 2018. Our estimate of global species-richness (Fig. S5a) is thus highly similar to previous projections (e.g., Rhodin et al. 2017).

The sea turtle data required additional processing. Species such as *Dermochelys coriacea* have been observed throughout nearly the entirety of the world’s oceans, though breeding ranges are substantially restricted, and neither are entirely comparable to the range maps of terrestrial species. As a compromise, we converted the oceanic-range polygons to lines, buffered the lines to 0.1 degrees (~11km at the equator), clipped the buffered lines to the 10m coastline polygons from Natural Earth (naturalearthdata.com), and used these coast-intersection polygons as estimates of the potential land-area of each species. The range data thus represent the vicinity of the shoreline in which each species can potentially be found, with near-shore urbanization or agriculture representing a proxy of disturbance.

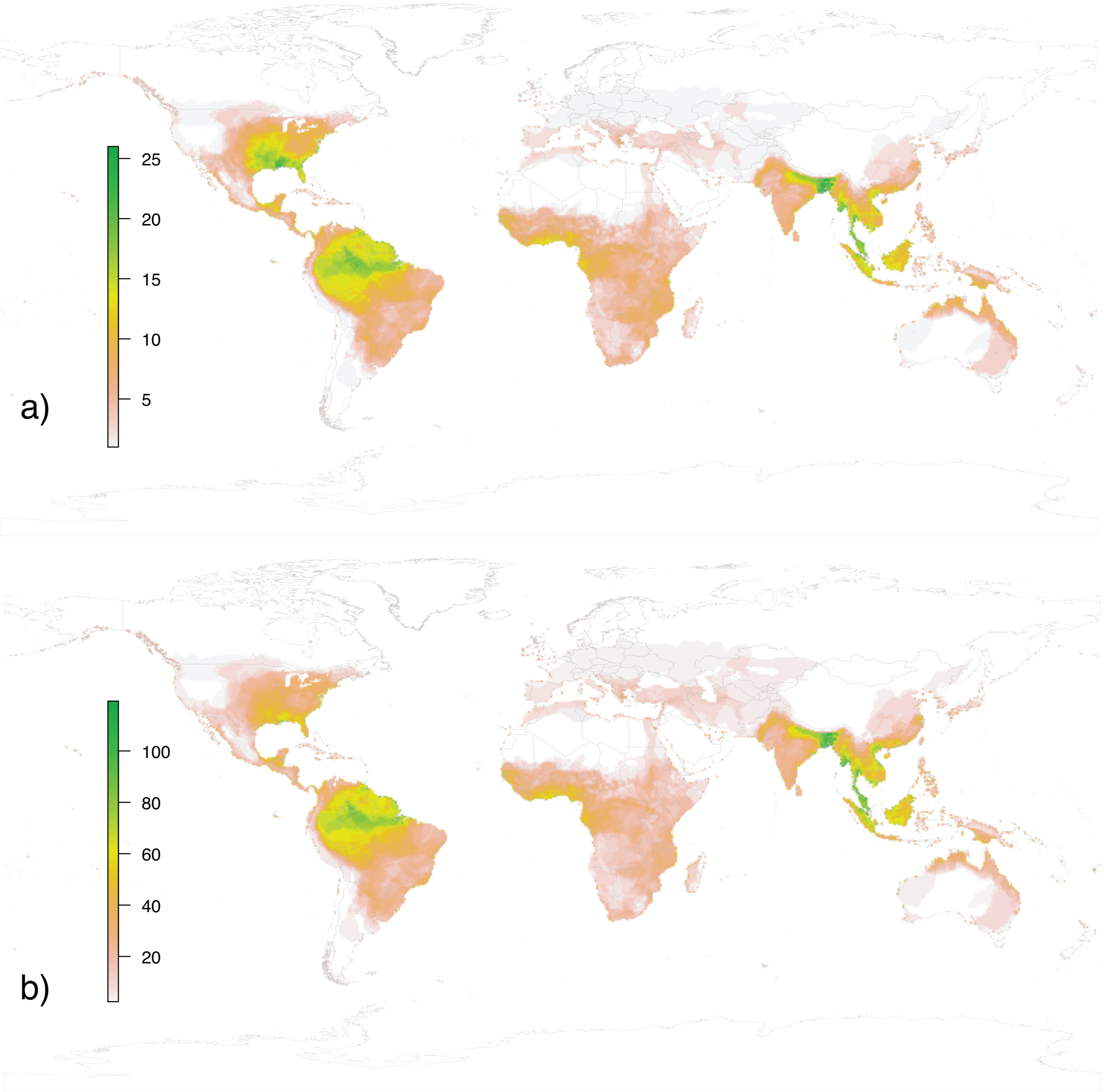

412

**Fig. S4.** Overall richness (a) and EDGE richness (b) of turtles (357 species) and crocodilians (27 species) in 1-degree grid cells. 413

To calculate the human-encroachment index (HEI; see Lee and Jetz 2011), we used the European Space Agency 300m landcover map for 2015 (http://www.esa-landcover-cci.org/), reclassified with urban (code 190) and cropland (codes 10-40) considered “modified.” We then intersected the range polygons for all species with this layer and calculated the proportion (ranging from 0 to 1) of each species’ range that is under human modification. This gives an initial estimate of anthropogenic pressure on each species due to habitat destruction. Finally, we calculated the centroid of each species’ range polygon, and estimated a distance matrix between all centroids as a measure of spatial proximity for potential autocorrelation of extinction risk.

As a final set of variables potentially influencing threat status in turtles, we measured several common axes of climate and ecosystem function. Climate has clear impacts on extinction risk due to anthropogenic change (Thomas et al. 2004), particularly for turtles (Ihlow et al. 2012) which have phylogenetically conserved climatic niches (Rodrigues et al. 2018). For climatic niche, we used the BIOCLIM dataset (Hijmans et al. 2005; Fick and Hijmans 2017), as re-interpolated by Kriticos et al. (2012) at 10-minute resolution as part of the CliMond dataset (Fig. S5). Specifically, they performed a spatial principal-components analysis on the 35 BIOCLIM layers to produce 5 synthetic layers (BIO36-40), capturing >90% of the total variation in the original dataset (Kriticos et al. 2014).

Based on variable loadings, these layers provide indices of temperature, wetness, dryness, rainy-season radiation, and warm-season radiation, respectively (see Kriticos et al. 2014). As with HEI, we calculated the mean value for each species by intersecting the range-map polygons with the climatic layers and calculating zonal statistics. Four island species (*Aldabrachelys gigantea, Chelonoidis abingdonii, C. duncanensis,* and *C. hoodensis*) were omitted by the spatial resolution of the climatic layers. Therefore, we imputed values for these species for BIO36-40 using the Rphylopars methodology described above (Appendix S4), including the observed values of AET and NPP (see below) for those species in the phylogenetic-trait covariance matrix (Fig. S6).

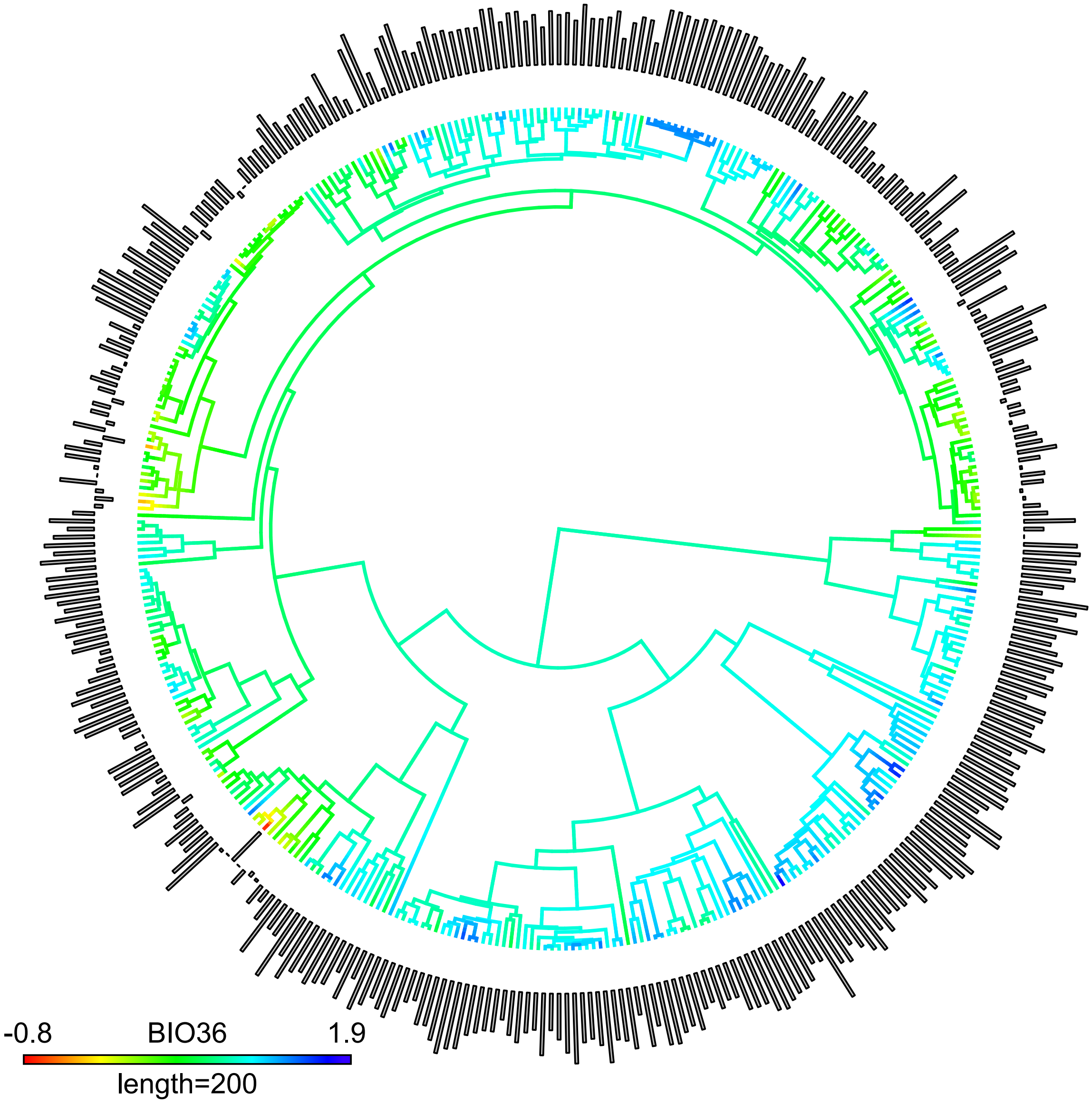


**Fig. S5.** Ancestral-state reconstruction and trait values for BIO36, the first principal-component axis of the 35 BIOCLIM variables, primarily measuring variation in temperature.

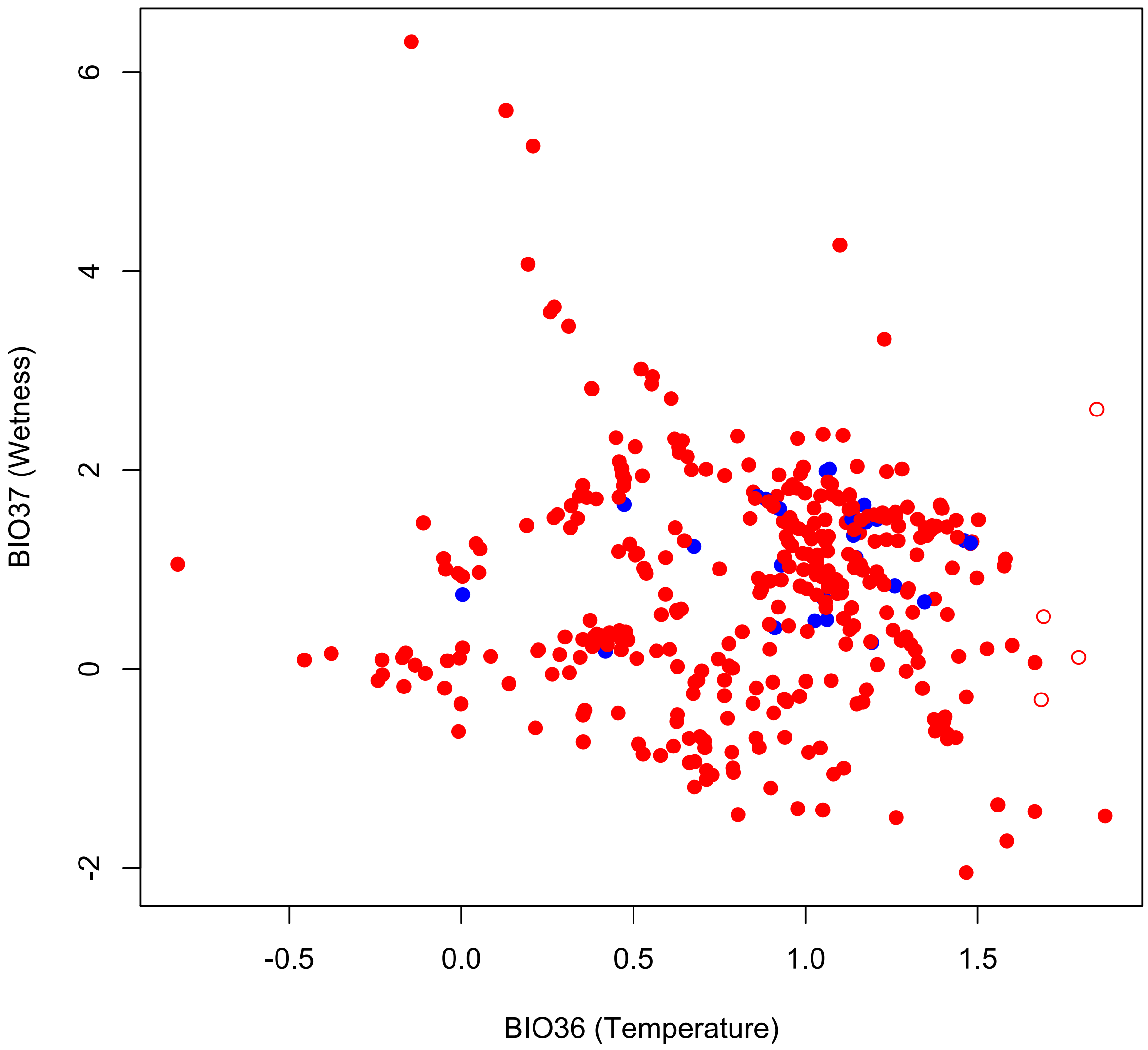


**Fig, S6.** Two uncorrelated principal-component axes (BIO36 & BIO37) estimated by Kriticos et al. (2014) for the 35 BIOCLIM variables (Hijmans et al. 2005). Red points are turtles (unfilled are imputed), while blue are crocodilians. Together, the first five axes (BIO36-40) account for ~90% of the variation in the BIOCLIM dataset, making them a robust proxy for climatic niche.

Finally, we used Actual Evapotranspiration (AET) and Net Primary Productivity (NPP) as measures of ecosystem function. These variables show strong correlations with historical diversification and present-day species-richness in amphibians, reptiles, birds, and mammals (Pyron and Wiens 2013; Coops et al. 2018). There are also large-scale correlations between these variables and anthropogenic disturbance, leading to significant conservation implications (Luck 2006). We used the MODIS products measured annually at 500m resolution, specifically MOD16A3 (AET, mean from 2000-2013; Running et al. 2017) and MOD17A3 (NPP, mean from 2000-2015; Running et al. 2015). As with the other spatial datasets, we used zonal statistics to measure the mean values within each species' range-map polygon intersected with the underlying climatic layer. All spatial analyses were performed in QGIS3.4 Madeira (https://qgis.org/en/site/).

*Threat Status Imputation*

For threat status, we first accessed the 29 November 2018 version of the IUCN RedList. We then changed *Nilssonia nigricans* from EW to CR because wild populations have recently been rediscovered (Praschag et al. 2007), and assigned *Aldabrachelys abrupta* and *A. grandidieri* to EX, based on their known status (Gerlach and Canning 1998). Thus, there are 54 Least Concern, 34 Near Threatened, 72 Vulnerable, 48 Endangered, 53 Critically Endangered, 9 Extinct, and 11 Data Deficient species. Overall, accounted for 258/357 turtles and 23/27 crocodilians, for a total of 281 assessments (270 with status) out of 384 total species. Remaining to impute are therefore 114 total species: 103 unassessed (99 turtles and 4 crocodilians) and 11 Data Deficient (all turtles). No species are currently Extinct in the Wild.

Multiple strategies for imputing threat status have been employed by previous authors, of which we chose PGLM estimation (Jetz and Freckleton 2015) and machine-learning classification and prediction (Davidson et al. 2009). Given multiple predictor variables and similarity matrices describing phylogenetic and spatial distance, we evaluated three primary methods for predicting extinction risk, which were all highly concordant. We then used the consensus as our best estimate for further analyses.

First, we used classical statistical models from the GLM framework, considering threat status as a continuous variable ranging from 1 (Least Concern) to 6 (Extinct). Our predictor variables (as described above) were a mixture of categorical and continuous. The binary categorical variables were clade (crocodilian or turtle), habitat (presence/absence in Freshwater, Marine, Terrestrial, Island, or Mainland), and ecoregion (presence/absence in the 11 biogeographic regions). The continuous variables were traits (length and mass), spatial attributes (area, HEI), and climate (BIO36-40, AET, and NPP). The continuous data were centered and scaled, except for HEI (which ranges from 0-1) and the BIOCLIM variables, which are already centered as a result of PCA transformation. This yielded a total of 28 predictors.

The full GLM containing all 28 predictors as main effects without interactions yielded a multiple-$R^2$ of 0.57, though only 11 of the predictors had significant effects. These were clade, length, Marine, Island, 4 ecoregions (MG, NEA, NCA, and OR), area, BIO36, and BIO 39. To avoid overfitting, we evaluated preliminary variable-importance using step-wise evaluation of AIC values under forward and backward addition of predictors. Furthermore, we assessed relative variable importance (Fig. S7) as the contribution of each predictor as a percentage of the multiple-$R^2$ partitioned by averaging over orders (Lindeman et al. 1980), conducted in the R package ‘relaimpo.’

This yielded a best-fit model of 15 predictors with a multiple-$R^2$ of 0.55, containing clade, length, mass, Marine, Terrestrial, Island, 5 ecoregions (IS, MG, NEA, NCA, and OR), area, HEI, BIO36, and BIO39. Interestingly, Terrestrial and IS were not independently significant in this model. The most important variable was area by far, followed by occurrence in the Oriental ecoregion. The remaining 13 variables were of modest importance overall in terms of variance partitioning. We used this set of 15 predictors for subsequent PGLM estimation following Jetz and Freckleton (2015).

We first fit a linear mixed-effects kinship model in the R package ‘lmekin,’ which allows species in a phylogeny to represent a random effect from a phylogenetic variance-covariance matrix representing their expected similarity, a number of fixed-effect predictors (the 15 described above), and an additional similarity matrix describing spatial proximity as a random effect. The phylogenetic and spatial matrices were standardized on the interval from 0-1. The random-effects matrices thus return a parameter describing the variance component attributable to evolutionary and geographic effects. This model was fit to the 270 species for which threat status was known. Given the estimated evolutionary and geographic variance components, we estimated a phylo-spatial matrix by multiplying each underlying matrix by its constituent variance component and adding them together. This model thus accounts for the 15 highest-importance predictors, as well as the phylogenetic relationships and spatial proximity of all species.

To estimate the 114 missing threat statuses, we then used the PGLM approach described by Jetz and Freckleton (2015) in the R package ‘PGLM.’ This method takes the 15-predictor input model and the phylo-spatial matrix to predict the unobserved values in the input data. Predictions are returned as continuous estimates, which were then rounded to the nearest integer and matched to the six IUCN threat-status categories. The estimation is not bounded on the interval from 1-6, and species could thus conceivably estimate values such as 0 or 7, which would not be inherently meaningful. However, all 114 estimates rounded to an integer from 1-6 and could thus be assigned a meaningful threat status.

We then estimated the sensitivity and specificity of this model using leave-one-out cross-validation, removing each of the 270 known statuses one-by-one, re-estimating them from the remaining data, and generating a confusion matrix of the known versus predicted statuses. The PGLM model had an overall accuracy of 38%, which is approximately twice as high as random; given a 6x6 matrix of potential outcomes from 6 categories, a random classification would yield 6/36 or 17% accuracy. The class-specific accuracy balanced by class prevalence (sensitivity + specificity/2) ranged from 57% for LC to 86% for EX.

Unlike the analyses in TESS/CoMet and Ievolution (see below) or Rphylopars (see above), this PGLM approach does not estimate parameters directly from the phylogeny. Because the number of imputed species in the tree is low and the phylogenetic variance-covariance matrix only appears as a down-weighted error term, the effect of phylogenetic uncertainty should thus be very low. Therefore, we did not replicate these analyses across a sample of trees, instead using a randomly-selected tree.

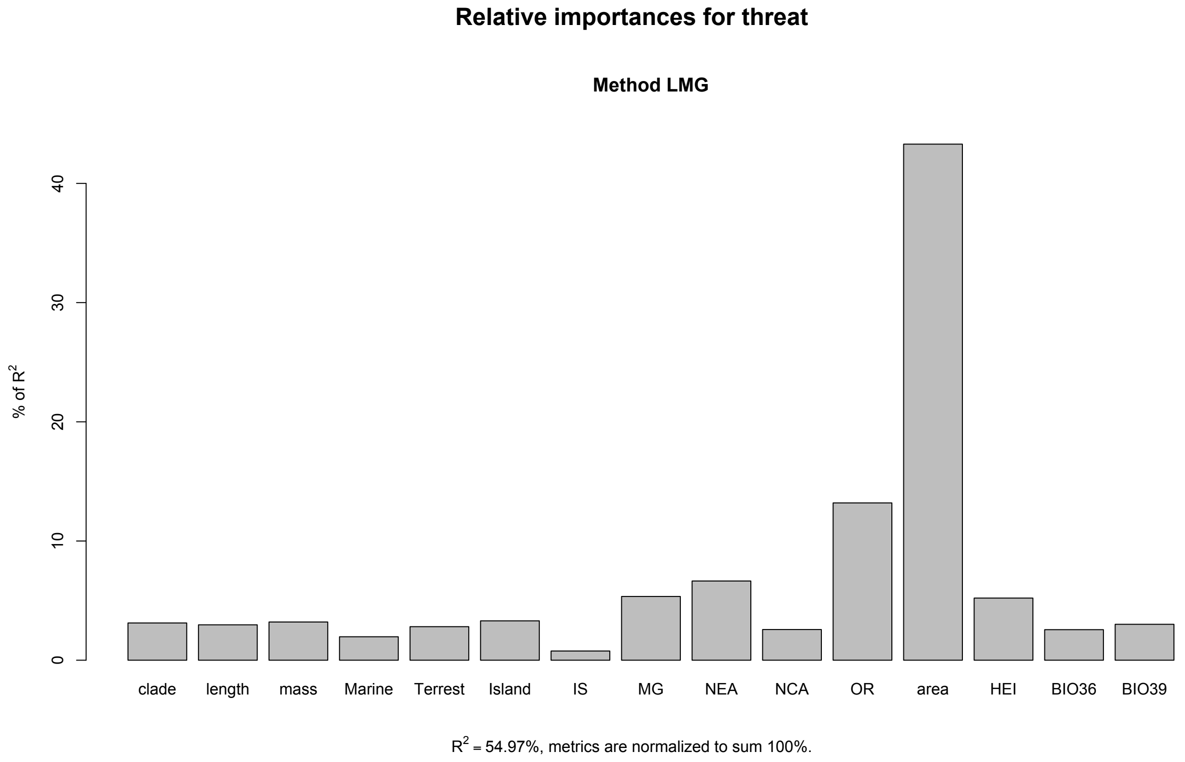


**Fig. S7.** Variable importance for the GLM consisting of the 15 top-ranked predictors, assessed as a percentage of the multiple $R^2$ partitioned by averaging over orders (Lindemann et al. 1980).

Subsequent to the PGLM methods representing the application of classical statistical techniques, we implemented to recently-developed machine-learning (ML) techniques (random forests and artificial neural networks) which are becoming widely adopted in ecological analyses (Cutler et al. 2007; Olden et al. 2008) and for conservation in particular (Bielby et al. 2009; Bland et al. 2015). The advantages of these techniques over approaches such as GLMs include higher classification accuracy, greater flexibility for addressing numerous problems such as regression and classification in a single modeling framework, and the intrinsic ability to detect and incorporate complex non-linear interactions between variables without a priori specification. All ML techniques were implemented in the R package 'caret.'

The first technique we employed was random forests (Breiman 2001; Cutler et al. 2007). In this algorithm, a forest of size 'ntree' is grown, consisting of decision trees which randomly sample a set number of predictor variables 'mtry' at each node, and continue branching until the Gini coefficient stabilizes. Predictions are then combined from all the trees in the forest. To assess accuracy, we performed repeated *k*-fold cross-validation, splitting the training dataset of 270 observations into 10 groups. Each group is then removed from the model (out-of-bag observations) and predicted by the remaining data. This procedure was repeated three times, across a range of 'mtry' values for 'ntree'=500. We selected the value of 'mtry' that maximized accuracy (Fig. S8). This procedure generates a measure of variable importance (scaled from 0-100) indicating the contribution of each factor (Fig. S9).

We included the 28 categorical and continuous predictors described for the PGLM approach, plus estimates of evolutionary and geographic covariance. Because the ML techniques do not easily accommodate full phylogenetic variance-covariance or spatial-proximity matrices, we summarized these matrices using principal-components analysis. We retained the first two axes (accounting for >90% of the variance) from each matrix and included them as continuous variables in the random forest analyses, for a total of 32 predictors. Continuous variables measuring real quantities (e.g., mass, area, distance) were log-transformed but not standardized, as random forests are invariant to scale since variables are considered sequentially. This allows decision trees to be interpreted in the units of the original variables.

The best value of ‘mtry’ was 5, yielding overall classification-accuracy of 49%, nearly three times the randomly-expected rate of 17% and 29% higher than the PGLM results. The out-of-bag estimate of error rate was 53%, with class-specific rates ranging from 30% (LC) to 75% (EN). However, these errors were generally over-estimates of extinction risk, yielding classifications which are thus beneficially conservative for our purposes. The variable-importance measure identified area as far and away the most important predictor (100) of threat status. A second group of 14 predictors range from 55-67, including traits (length and mass), the 4 phylogenetic and spatial components, climate (BIO36-40, AET, and NPP), and human encroachment. Occurrence in the Oriental ecoregion has a modest impact (22), and the remainder of the ecoregion, habitat, and clade variables have minimal impacts. An example tree shows the overwhelming influence of area, encroachment, and climate on the first three nodes (Fig. S10).

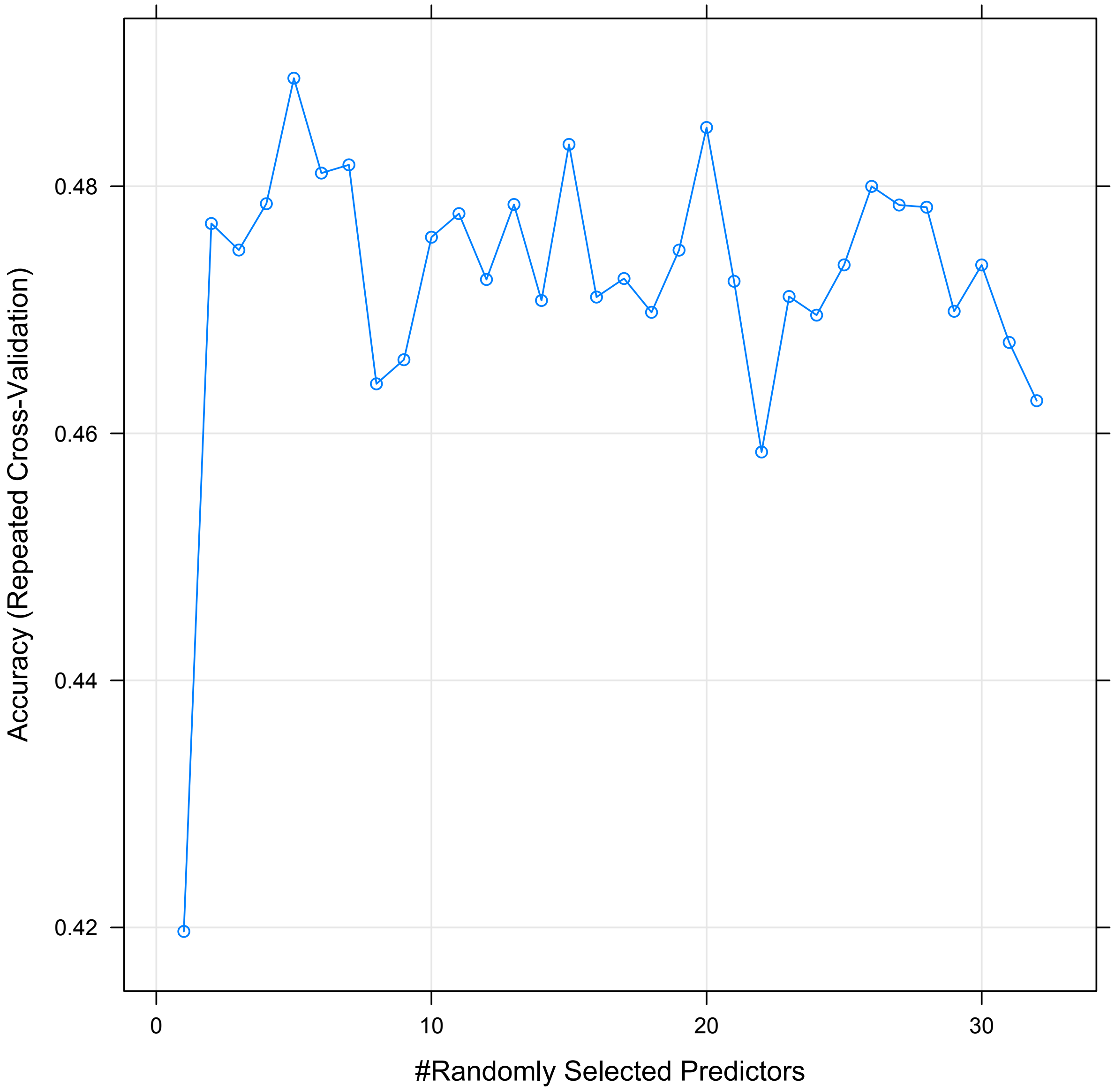


**Fig. S8.** Accuracy (repeated *k*-fold cross-validation) of random-forest models across values of mtry at each node.

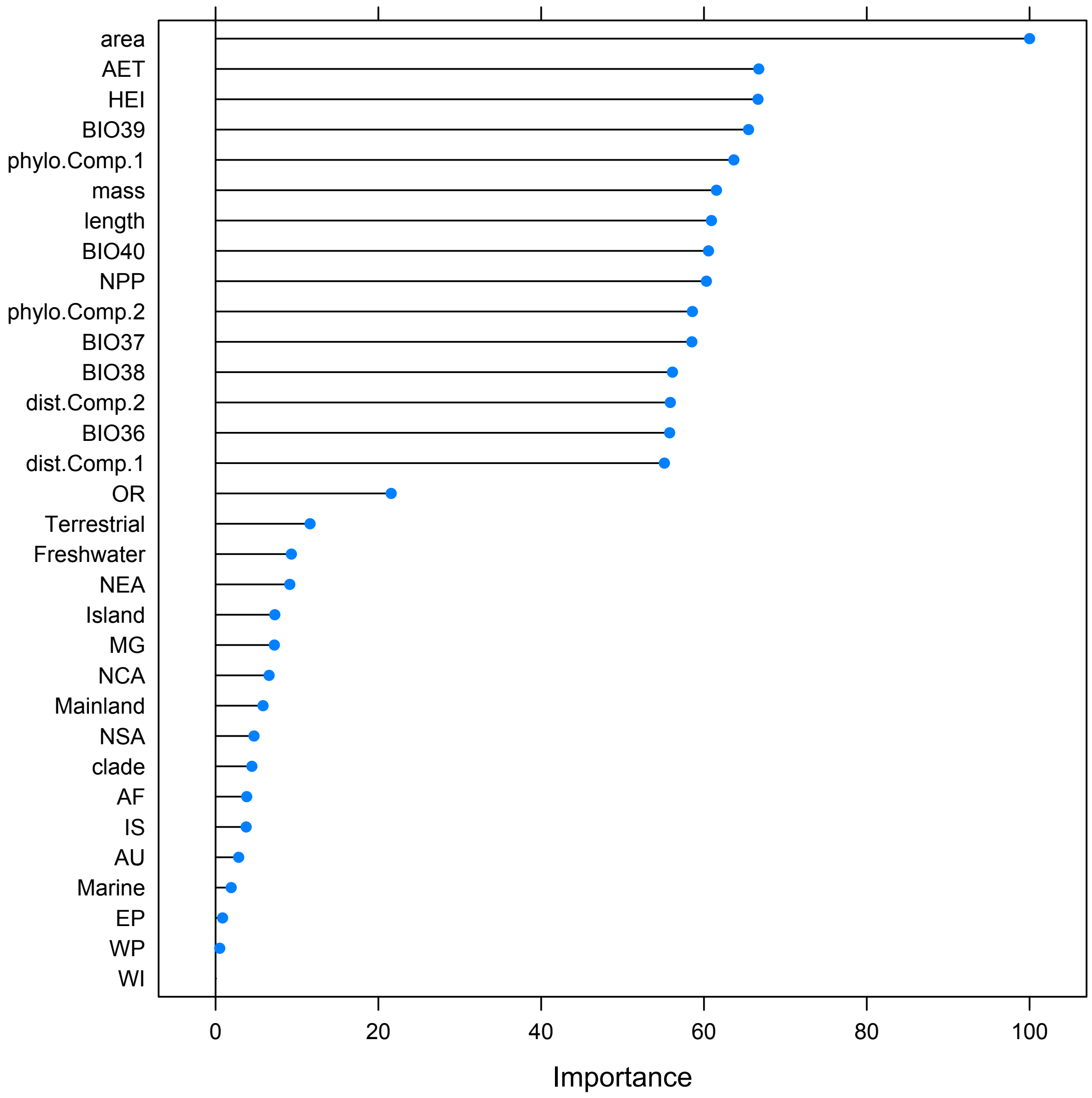


**Fig. S9.** Scaled variable importance for predictors in the random-forest model.

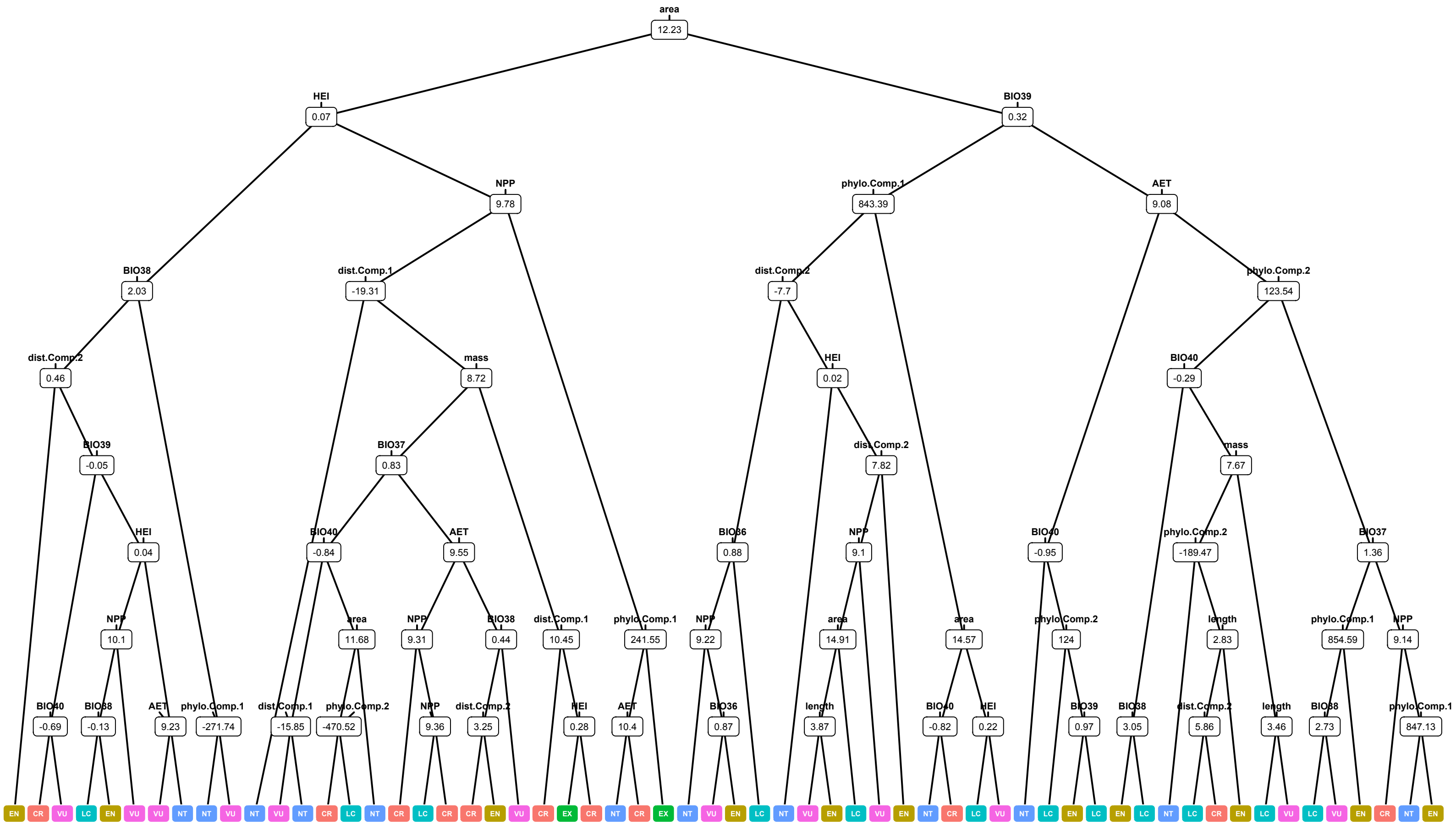


**Fig, S10.** Example decision-tree from the RF model. For illustrative purposes, this tree is the smallest representative (in terms of number of nodes) in the random forest generated with the reduced subset of variables having >50 relative importance in the main model.

Finally, we used a simple implementation of artificial neural networks, the multi-layer perceptron (see Olden et al. 2008). The classic MLP algorithm is a supervised, fully-connected, feed-forward network, trained by back-propagation. Each feature class (predictor variable) is connected to a single neuron in the input layer. Each input neuron is connected to each of a given number of processing neurons in the hidden layer. Each hidden neuron is then connected to every neuron in the output layer; in a classification problem such as this, there are as many output neurons as there are states. The hidden and output layers are each also connected to a bias neuron, which adds a constant output. The connections (axons) between each neuron each have a unique weight (analogous to a partial regression coefficient) that is multiplied by the input signal, and these summed as the input to the next layer. The input signal to each neuron in the hidden layer is then fed into an activation function (typically sigmoid), which determines whether that neuron "fires" a weighted signal towards the output layer. The sum of the weighted input signals to the output layer and their subsequent activation determines the final output of the model; a categorical determination in the case of a classification problem such as this one.

The learning stage consists of estimating the weights and biases using a back-propagation algorithm on a training dataset with known inputs and outputs, by minimizing a root-mean-squared-error cost function. This yields a fully-trained network which can then accept an input dataset and predict the output variables with high accuracy. As with the PGLM and random forest models, we trained the network using the 270 species with known statuses, tuning the model using repeated *k*-fold cross validation to select the number of hidden neurons and the weight decay (a regularization parameter for the weights) yielding the highest training accuracy (Fig. S11). The estimated weights can then be compared across input features to assess variable importance (Garson 1991).

We used the same 32 input features as in the random forest model. However, neural networks generally require standardization to the interval [0-1], which we achieved using min-max scaling for the continuous variables. As the evolutionary and geographic matrices now represented scaled Euclidean distances, we thus principal coordinates (rather than principal components) analysis for dimensionality reduction, though this is approximately equivalent mathematically to the PCA performed on the original matrices. We again retained the first two axes for each matrix.

The variable importance measures again identified area as the most important predictor, but with a slightly different set of secondary predictors (Fig. S12). The upper 50% of these include the phylogenetic and spatial components; occurrence in terrestrial, freshwater, and island habitats; length and mass; temperature and radiation (BIO36 and BIO39); and occurrence in the Oriental, Nearctic, Neotropical South America, and Neotropical Central America ecoregions. The best-fit model contained 22 hidden neurons with a weight decay of 0.1, yielding 48% training accuracy (Fig. S13).

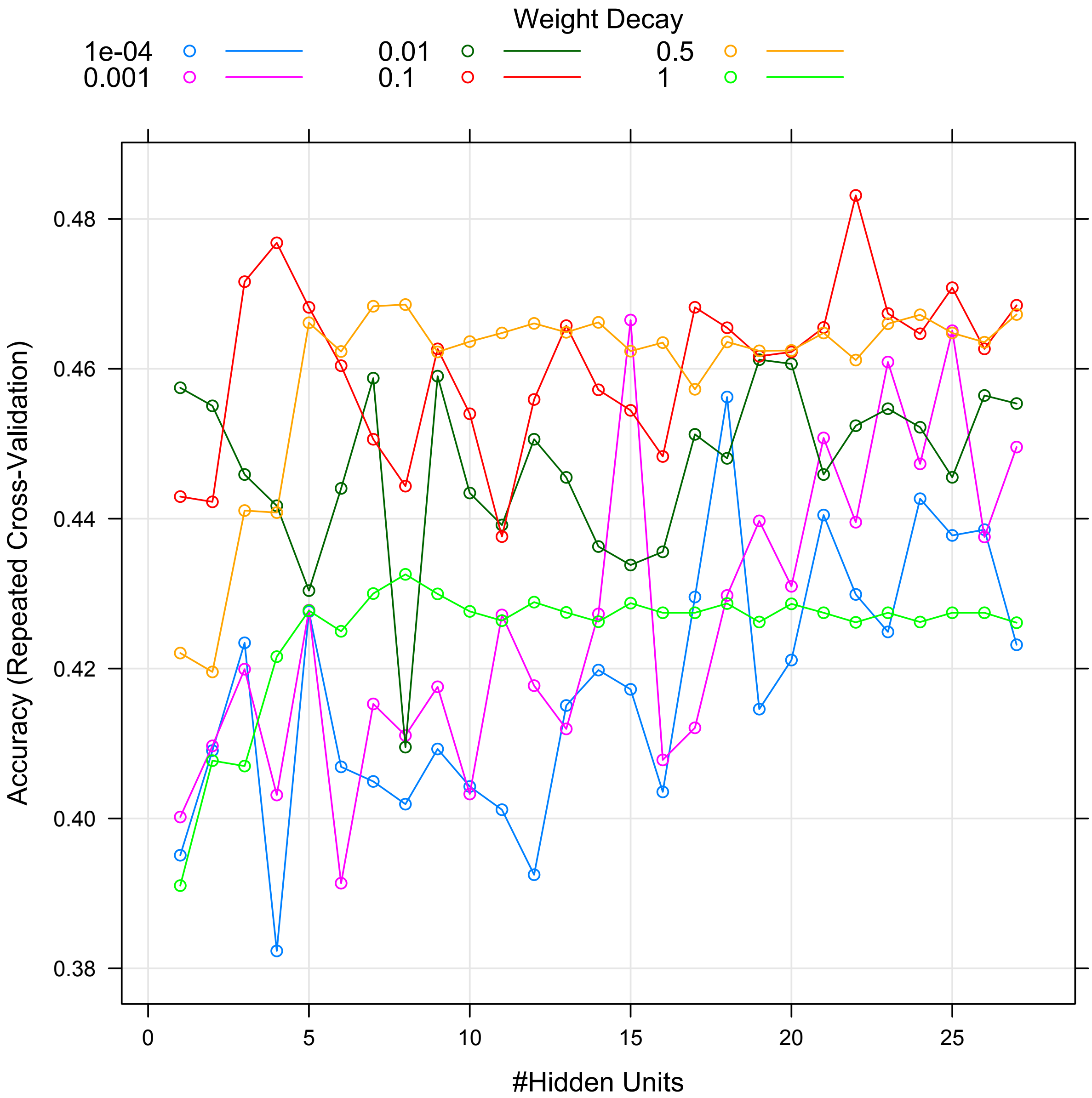


**Fig. S11.** Plot of accuracy (repeated *k*-fold cross-validation) across different parameter values for network size (hidden units) and regularization (weight decay) for the MLP neural-network model.

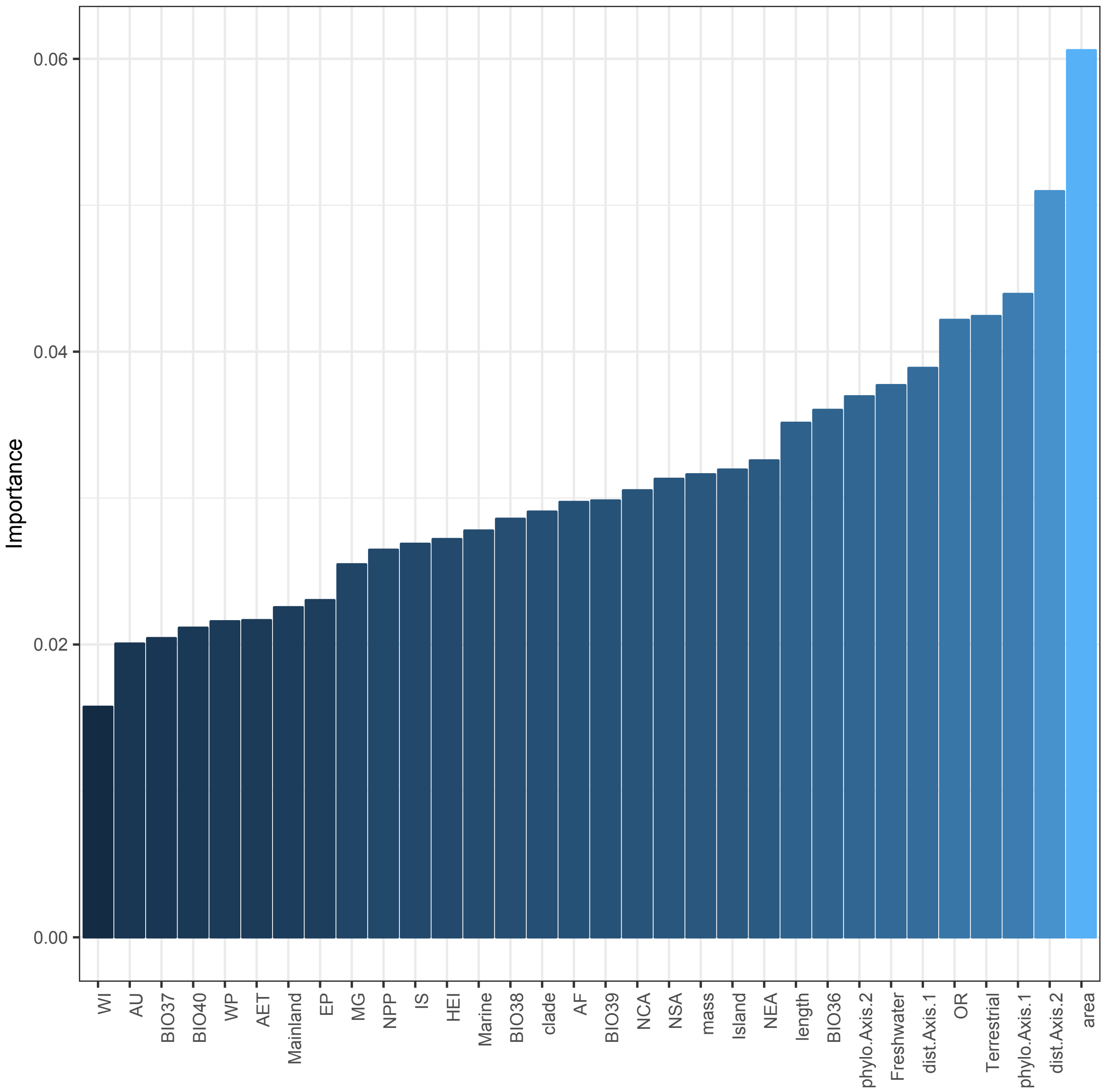


**Fig. S12.** Plot of variable importance (Garson 1991) for the MLP neural-network model.

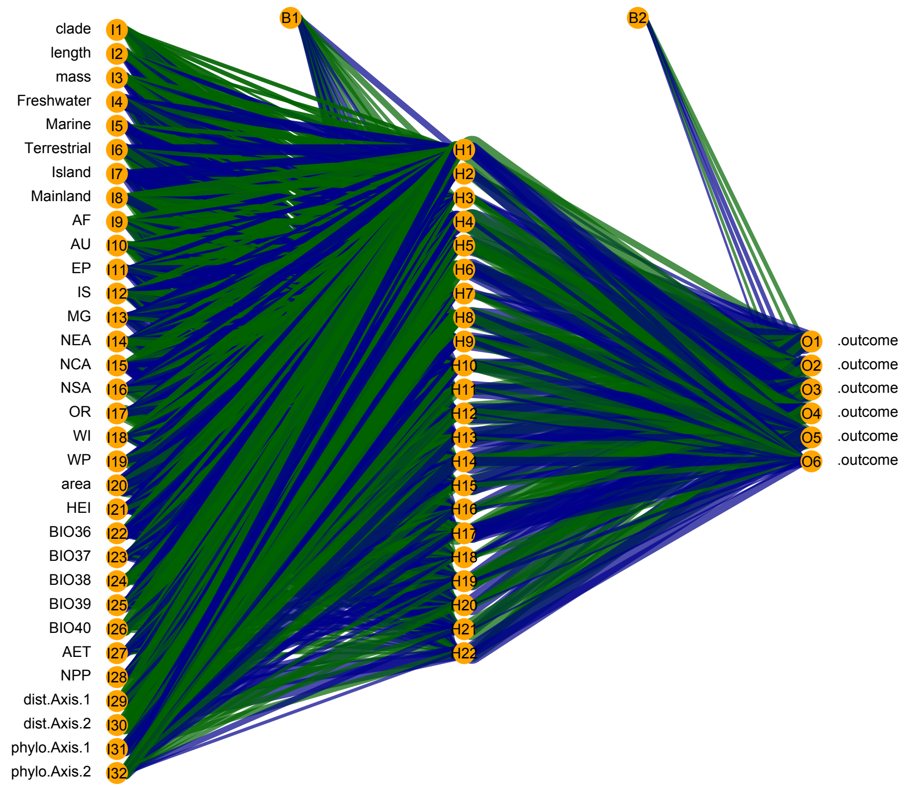


**Fig. S13.** Plot of the structure of the best-fit MLP ANN model. Positively-weighted connections are green, negative are blue. From left to right, orange cirlces labeled with “I” are input nodes (predictor variables), “H” are hidden nodes (processing neurons), and “O” are the outcome states (six IUCN threat-status categories). The biases are labeled B1 and B2, connecting to the hidden and output layers. Axons are scaled relative to their weights, indicating the strength of feature importance across layers.

The final predictions for the 114 species are fairly similar (Appendix S5), with all models identifying area as the single-most important variable, though each method picked out slightly different contributions from secondary predictors. Overall, 21 species were predicted identically by all three approaches, and 69 agreed for two out of the three, for ~79% concordance overall. On a pairwise basis for 342 comparisons (114 species times three models), 141 were identical (41%) and an additional 144 (42%) were adjacent (one category different), for a total of 83% identical or adjacent predictions. Given the high level of agreement and lack of apparent bias towards higher or lower categories for any model, we simply used the mean of the three predictions for our final estimate of threat status.

Our final estimate included 17 species classified as LC, 27 NT, 47 VU, 15 EN, and 8 CR. Imputed statuses showed a similar distribution of ED to known species (Fig. S14). Qualitatively, imputed species were heavily concentrated in tropical pleurodiran lineages, particularly those with many recently-described or resurrected species such as *Pelusios* (see Fritz et al. 2011) and *Pelomedusa* (see Petzold et al. 2014). Nearly all of the sub-Saharan African pelomedusines, and ~2/3 of the Australasian chelodinines were imputed. No other subfamily with more than 2 species had fewer than 50% assessments.

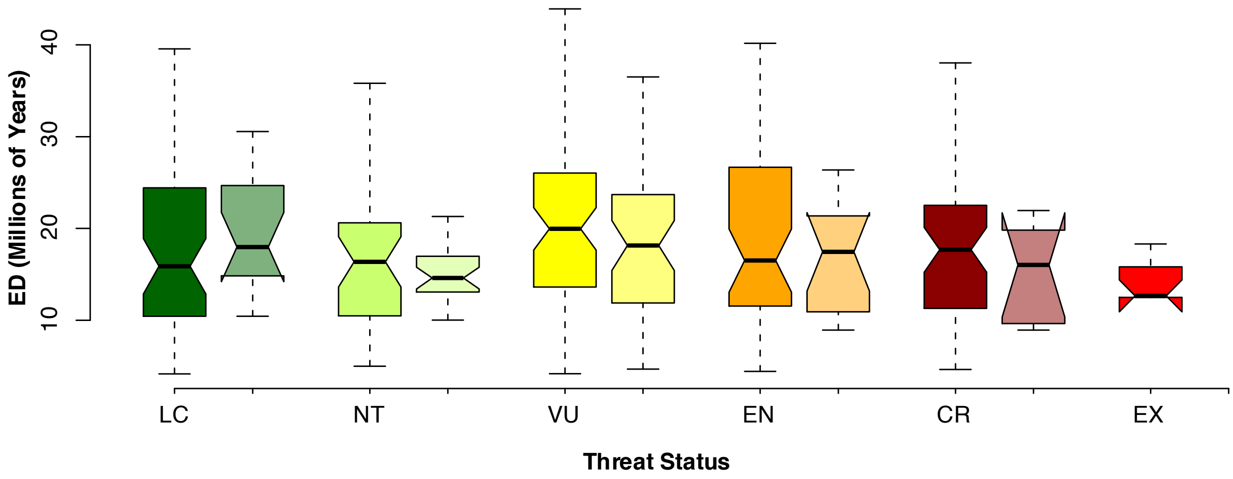


**Fig. S14.** Comparison of ED for known (solid) and imputed (transparent) threat statuses.

The final breakdown of assessed plus estimated threat statuses used for subsequent analyses was 71 LC, 61 NT, 119 VU, 63 EN, 61 CR, and 9 EX. Thus, 66% of recent turtle and crocodilian species are imperiled. We then calculated the so-called 'EDGE' scores (Evolutionarily Distinct, Globally Endangered using the approach of Isaac et al. (2007), as EDGE = ln(1 + ED) + GE * ln(2), where GE is threat status on the scale of 0-5. When a species is not threatened, ED is equal to EDGE; scores diverge the more threatened a species is, and divergence in their histograms is an indicator of overall threat status of a lineage.

We also produced an estimate of global EDGE-richness (Fig. S5b), wherein species richness in each 1-degree grid cell is weighted by the sum of EDGE scores for the species in that cell. In this estimate, areas such as the eastern Nearctic are significantly down-weighted in prominence. This results from the profusion of low-ED species in lineages such as Deirochelyinae that we suggest may be the result of taxonomic overinflation (see above). In contrast, other major hotspots such as South and Southeast Asia, Australo-Papua, western Africa, and the Amazon Basin display similar prominence.

Previous estimates of EDGE scores for turtles and crocodilians were produced by Gumbs et al. (2018) based on the scanty available phylogenetic coverage for the group. Those authors used the existing incomplete IUCN assessments for threat status and phylogenetic imputation methods to estimate ED for unsampled species. Thus, their estimates of ED and EDGE may be over- or under-estimated compared to our taxonomically complete dataset, and thus the rankings of some species for conservation priorities may change. We compare our results in terms of both overall correlation (to identify systemic differences) and in terms of the top-10 list of most distinct and endangered species for further attention.

Comparison of ED values (Fig. S15) reveals fairly high similarity in taxa measured by direct estimation from a dated phylogeny. However, a large cluster of imputed ED values are significant over-estimates compared to our dataset. In contrast, EDGE scores were highly similar between the two methods (Fig. S16). However, due to differences in ED estimation or imputation or threat assessments, they heavily under-rank some species such as *Cyclanorbis elegans,* while over-ranking species such as *Chelodina novaeguineae.* As Gumbs et al. (2018) recognized the potential for error when working from a limited phylogenetic dataset, they presented only 'robust' EDGE scores, where robustness was assumed when a genus was fully-sampled in their phylogenetic tree. Thus, our top-10 list differs from theirs due to changes in threat status, differences in ED calculation, and variation in taxonomic completeness in the phylogenetic dataset which led to their inclusion of lower-ranked but 'robust' species in their top 10. We suggest that our estimates provide a more accurate picture of the EDGE ranking of turtles and crocodilians.

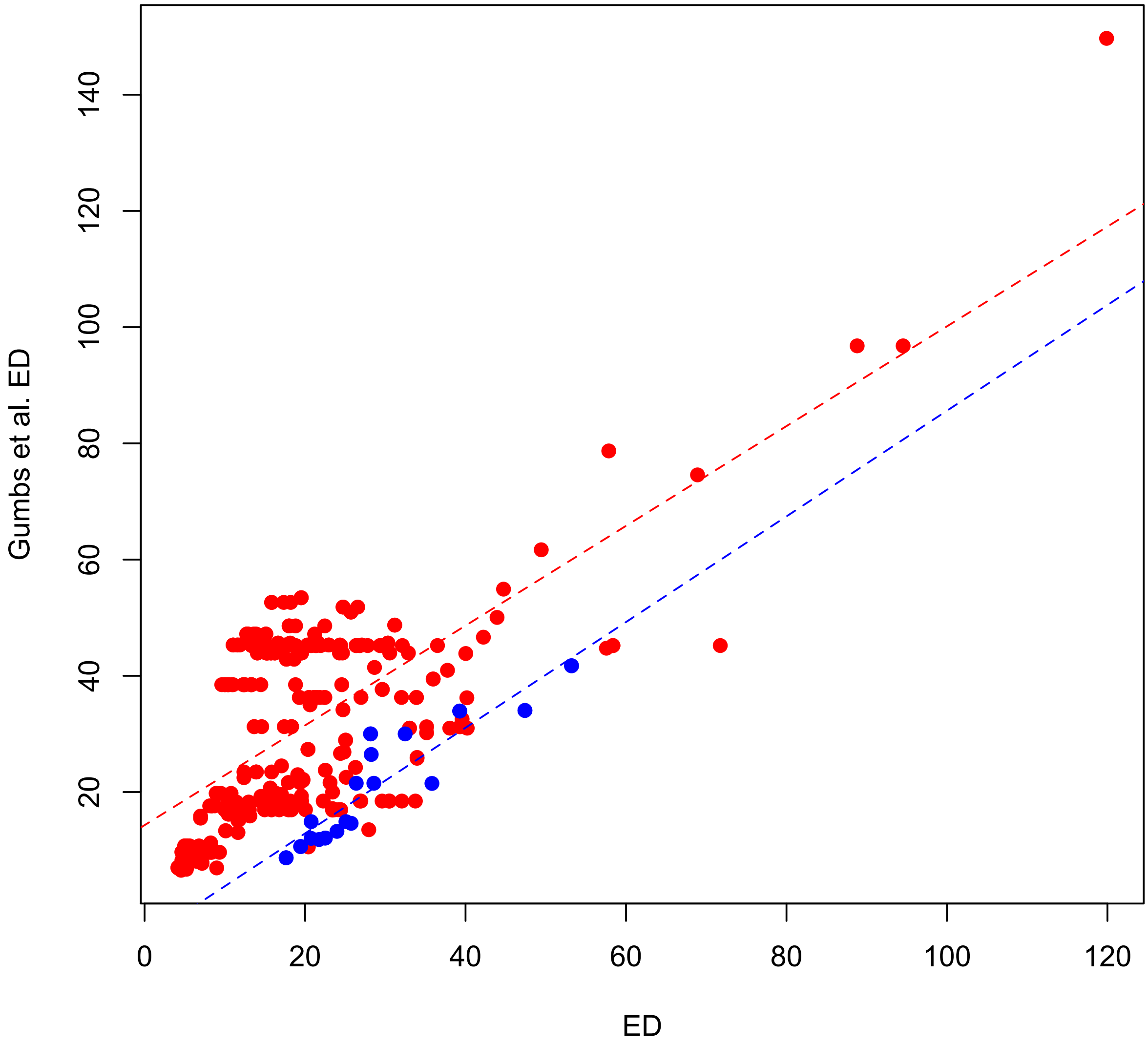


**Fig. S15.** Comparison of median ED from our phylogenetic estimate to the estimated + imputed ED values presented by Gumbs et al. (2018), with turtles in red and crocodilians in blue. For turtles, a cluster of Gumbs et al.'s values between 20 and 60Ma represent imputed ED values that are overestimates compared to our dataset, inflating EDGE scores for these taxa, which were not sampled in their phylogenetic trees. Estimates for crocodilians are generally very similar between both datasets.

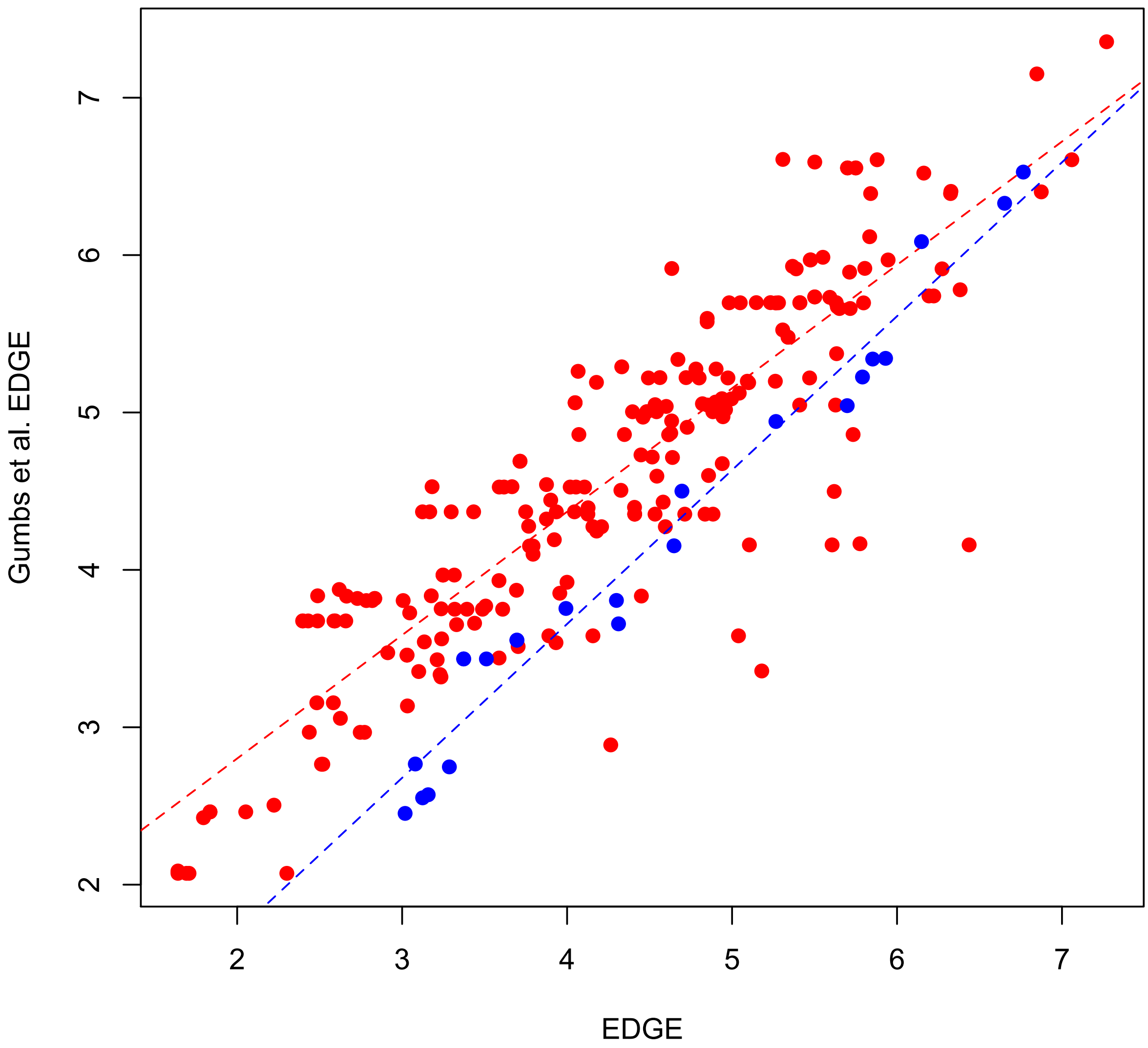


**Fig. S16.** Comparison of EDGE from our phylogenetic estimate to the estimated + imputed EDGE values presented by Gumbs et al. (2018). In contrast to the ED values (Fig. S15), the EDGE values are highly similar, with minor differences reflected in the top-10 lists generated from each dataset.

*Diversification Rates*

Numerous, varied approaches exist to estimate diversification rates (see Pyron & Burbrink 2013), including constant rates, or variation through time, or across lineages (Alfaro et al. 2009; Morlon et al. 2011; Rabosky 2014; May et al. 2016). Models for estimating significant shifts in among-lineage diversification have been applied to turtles, which estimated significant increases in the Galapagos tortoises and deirochelyine pond turtles, while all other species were part of the same background regime (Rodrigues and Diniz-Filho 2016). As noted above, these two clades exhibit high levels of apparent taxonomic overinflation, and rate estimates dependent on the total number of species and their associated branch lengths are thus suspect. Few if any similar analyses have been performed on crocodilians, but both groups have a rich Mesozoic and Cenozoic fossil record (Gaffney and Meylan 1988; Markwick 1998).

To avoid potential issues with low-level taxonomic inaccuracies in turtles, we instead focus on temporal variation in whole-lineage speciation and extinction rates, with specific reference to testing for mass extinctions at the K-Pg boundary. Both turtles and crocodilians showed high (50-80%) survival across the K-Pg boundary (Macleod et al. 1997; Novacek 1999; Brochu 2004; Ferreira et al. 2018), as well as strong diversification in the Jurassic, Cretaceous, and Paleocene (Markwick 1998; Nicholson et al. 2015).

Thus, we used the CoMet model in the TESS package (Hohna et al. 2016), which estimates speciation and extinction rates through time across a clade while allowing for shifts, as well as testing for mass extinctions. We do not specify the hypothesized location of any rate shifts or mass extinctions beforehand, and the model is thus agnostic to their temporal location. However, we generally hypothesize that if significant shifts or extinction periods are observed, they would be localized near the K-Pg boundary. Specifically, we inferred rate estimates for speciation and extinction with a Bayesian variable 'birth–death' model (Hohna 2013), conditioned on turtles and crocodilians separately. Jointly with this analysis, we also searched for potential evidence of past mass extinctions using the ‘compound Poisson process on mass-extinction analysis’ (CoMet; May et al. 2016).

These models are implemented in the TESS package (Hohna et al. 2016). We sampled ten trees from the posterior of each of the two clades and modelled a variable ‘birth–death processes’ with explicit mass-extinction events following the guidelines in Hohna et al. (2016), with the sampling fraction set to 1 and the expected survival of a mass extinction equal to 0.05 (95% extinction). We generated empirical hyper-priors through an initial Bayesian Markov chain Monte Carlo analysis under a constant-rate birth–death-process model to determine reasonable hyperparameter values for the diversification priors. We allowed up to two mass-extinction events and two rate change events and set the shape parameter for the mass-extinction prior to 100; varying these parameters yielded qualitatively similar results. After burn in, we ran final analyses for 100,000 generations and diagnosed models using effective sample size (>200) and the Geweke statistic. We evaluated significance of rate shifts and mass extinctions using Bayes factors, mentioning instances >6 and considering >10 strong support.

Results for both turtles (Fig. S17) and crocodilians (Fig. S18) showed roughly constant rates of speciation and extinction, with no support for any shifts, and no support for any mass-extinction events. Turtles exhibit speciation rates of ~0.07 lineages per million years and extinction rates of ~0.03-0.04, for net diversification rates of ~0.03-0.04 and turnover of ~43-57% from the Jurassic to the present. For crocodilians, these rates are ~0.05 and ~0.02 respectively, for net diversification of ~0.03 and turnover of ~40% from the Cretaceous to the present. Because of the low number of imputed taxa, fixed backbone, robust age-constraints, and lack of significance in the preliminary runs, we present results from a single tree; additional posterior samples do not support substantially different results. This is a conservative approach, given the limitations of these trees and models (Rabosky 2010, 2015; Title and Rabosky 2016).

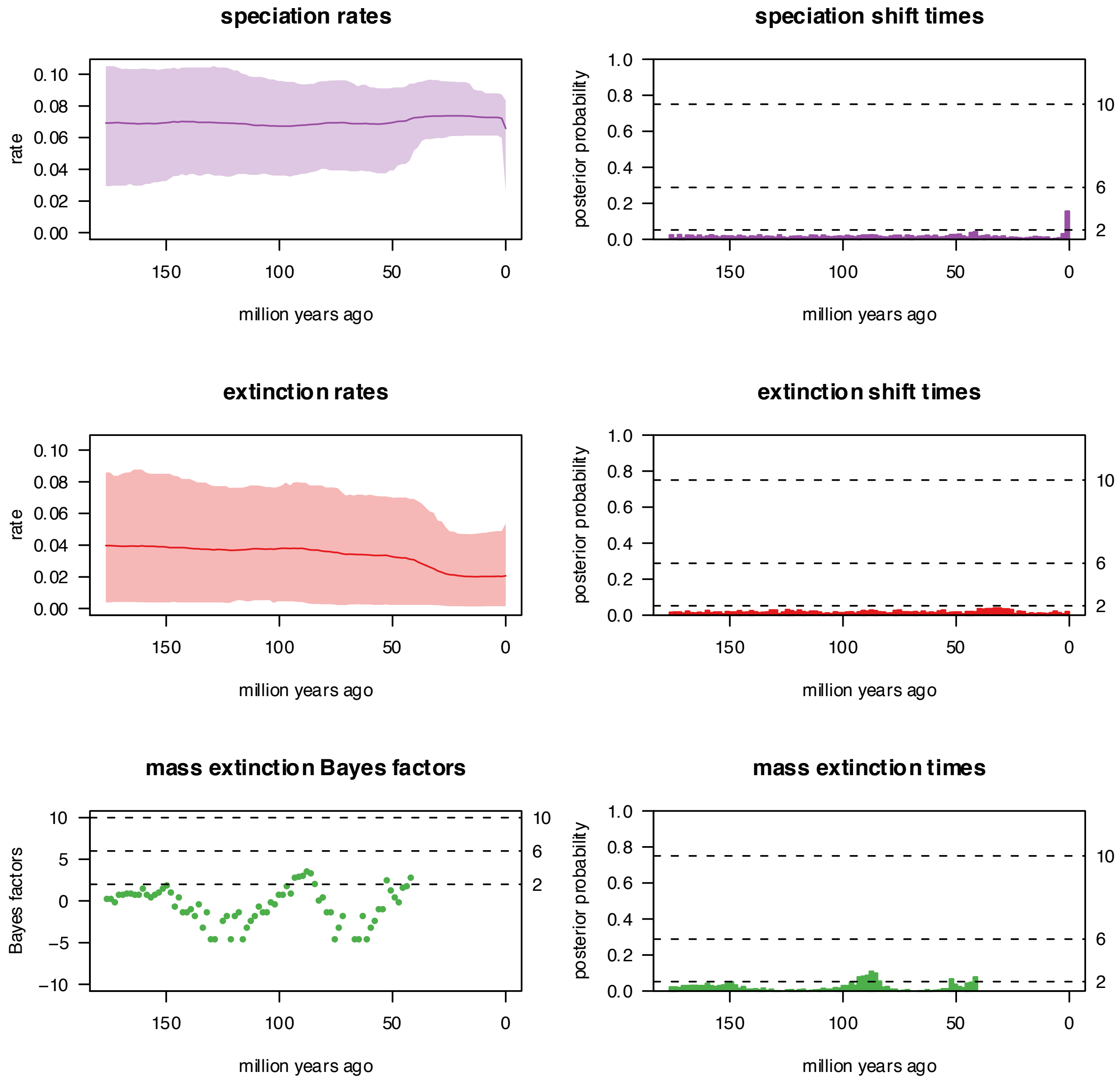


**Fig. S17.** Results from a single TESS/CoMET analysis for turtles, showing constant speciation and extinction rates, with no support for shifts therein, or for any mass-extinction events.

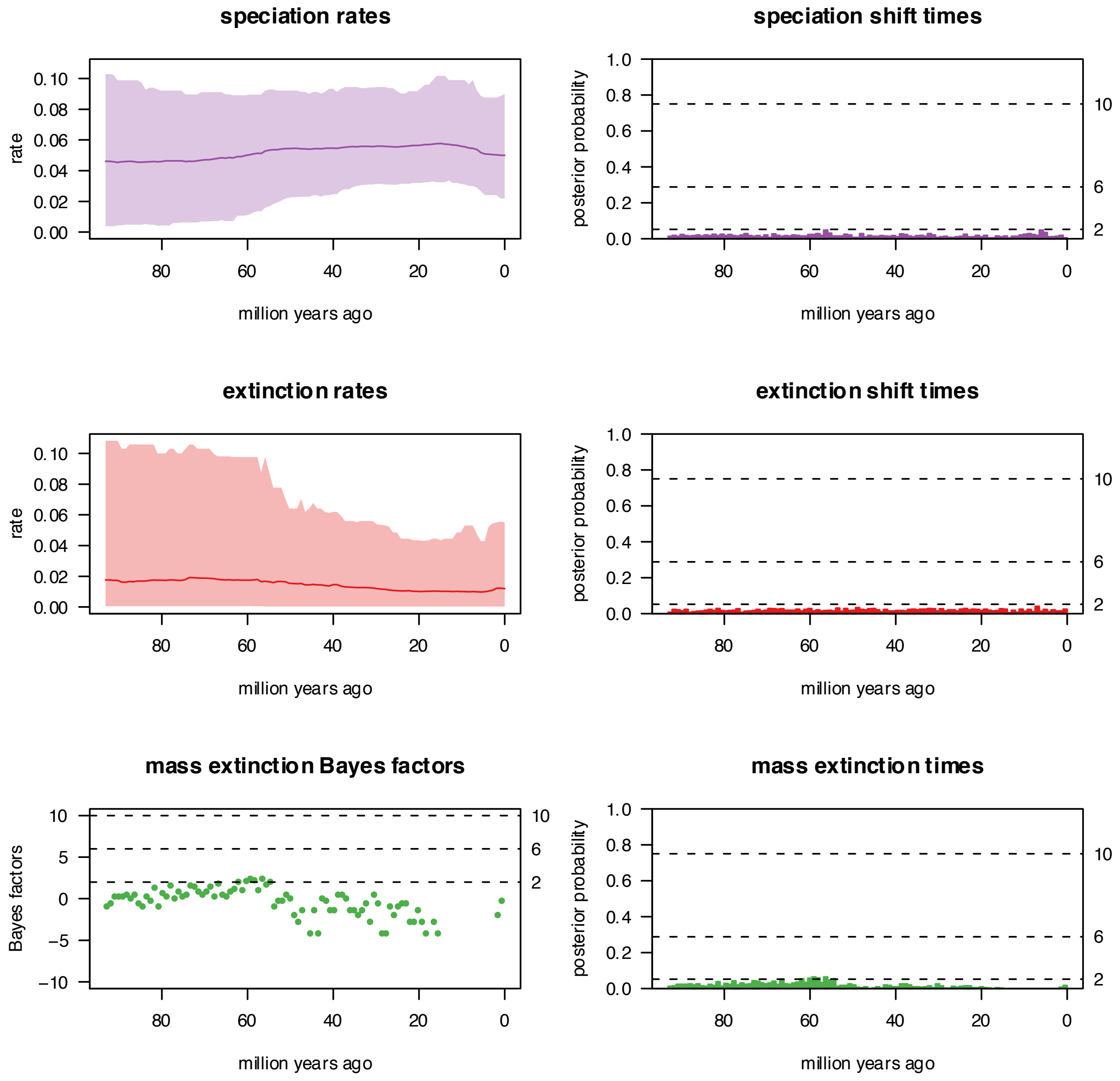


**Fig. S18.** Results from a single TESS/CoMET analysis for crocodilians, showing constant speciation and extinction rates, with no support for shifts therein, or for any mass-extinction events.

*Body-Size Evolution*

Numerous methods exist for analyzing the evolutionary history of traits on phylogenetic trees (e.g., Butler and King 2004; Eastman et al. 2011; Uyeda and Harmon 2014), allowing for a great degree of flexibility in specifying historical scenarios and testing hypotheses. Caution is also required, as many of these methods are limited by statistical power and identifiability of parameters (Boettiger et al. 2012). Methods evaluating quantitative shifts in characters across branches have been applied to turtles to study body size, indicating different selective optima for marine turtles, island tortoises, freshwater turtles, and mainland turtles and tortoises (Jaffe et al. 2011). A subsequent study estimated decreased rates of body-size evolution in geoemydid turtles and increases in emydids and testudinids (Eastman et al. 2011). However, these latter two clades include the questionably-delimited sliders and map turtles (*Pseudemys* and *Graptemys*) and Galapagos tortoises (*Chelonoidis*) described above, and the methods may thus be partially affected by an artifactual collection of short terminal branches.

Cognizant of the statistical issues intrinsic to the methods (Ho and Ané 2014), empirical issues related to potential taxonomic inflation (see above), and computational burdens associated with more complex methods (Uyeda and Harmon 2014), we limited our analyses to a small set of simple, tractable hypotheses. Specifically, given the strong conservatism in body plan but massive variation in body size in turtles and crocodilians, we ask whether any significant shifts in body size evolution are evident in the phylogeny of extant taxa. We acknowledge that this omits the huge amount of diversity observed in the fossil record of the group which would likely improve estimates (Slater et al. 2012). However, previous analyses suggest the signal is still present in extant taxa (e.g., Eastman et al. 2011; Jaffe et al. 2011).

We use the program *levolution,* which models the rate, strength, and phylogenetic position of evolutionary jumps using a Levy process (Landis et al. 2013) to model saltational changes in Brownian Motion regimes in a trait across a tree (Duchen et al. 2017). This approach has a variety of statistical, theoretical, and computational advantages over existing methods (Landis and Schraiber 2017). These include computational simplicity and reduced computing time, greater identifiability of parameters, and better theoretical fit to instances of rapid evolutionary bursts punctuating apparent stasis in phenotype. We only evaluated the length data (including the single imputed value for *Osteolaemus afzelli*), as analyzing the phylogenetically-imputed mass data on the same phylogeny would be circular.

We used a multi-stage analytical strategy to optimize the relevant parameters and infer the likelihood of a jump model (over a null tree-wide Brownian Motion process) and the location and posterior probability of shifts. The program *levolution* employs up 100 rounds of Expectation-Maximization to estimate parameters (the mean, variance, and jump rate of the process) using Markov Chain Monte Carlo sampling, and then uses a second MCMC analysis with those parameters to infer the topological position of jumps. This is then compared to the null model by likelihood-ratio test. Given the fixed nature of most of the topology and the computational burden, we ran these analyses on a single randomly-selected tree.

We first estimated the value of the hyperparameter $\alpha$ using the included dynamic peak-finder algorithm, which permutes $\alpha$ until the highest likelihood is found. We used the recommended settings of 5000 sampled jump-vectors, 1000 burnin samples, and $2^{nd}$-generation thinning for the MCMC chains, and a starting value and step value of 0.5 (on a log scale), optimized 5 times. The program thus tested 14 values of $\alpha$, with 31.6228 being the final value. The LRT was significant ($D$=175, $P$<1e-16), suggesting that a jump-diffusion Lévy process best fits the data. We then ran a final analysis with $\alpha$ fixed to this value, starting from the ML estimates of the mean, variance, and jump-rate parameters. For this final analysis, we increased the number of sampled vectors in the MCMC chain to 25000 for both the parameter estimation and the jump inference, accepting jumps with posterior probabilities >0.85.

Consistent with the relative stasis observed in the fossil record, relatively few strongly-supported jumps are estimated by the model. Thus, despite the massive variation in size observed in both extinct and living species, most of this variation can be attributed to steady drift over time. Two major and three minor jumps are significant in our final analyses. The first occurs along the branch leading to extant crocodilians, reflecting the fundamental difference in body size between the two groups. The second occurs in the highly-distinct relict species *Carettochelys insculpta,* which is the sister lineage of (and generally much larger than) the radiation of trionychid softshell turtles. Three terminal species are also estimated as jumps, each of which is significantly smaller than its congeners. These are *Pelodiscus parviformis, Pseudemys gorzugi,* and *Trachemys adiutrix.* As noted above, the latter two radiations have highly questionable species boundaries, as does *Pelodiscus* (Yang et al. 2011). Thus, we refrain from making any interpretation of these estimates due to their potential ambiguity.

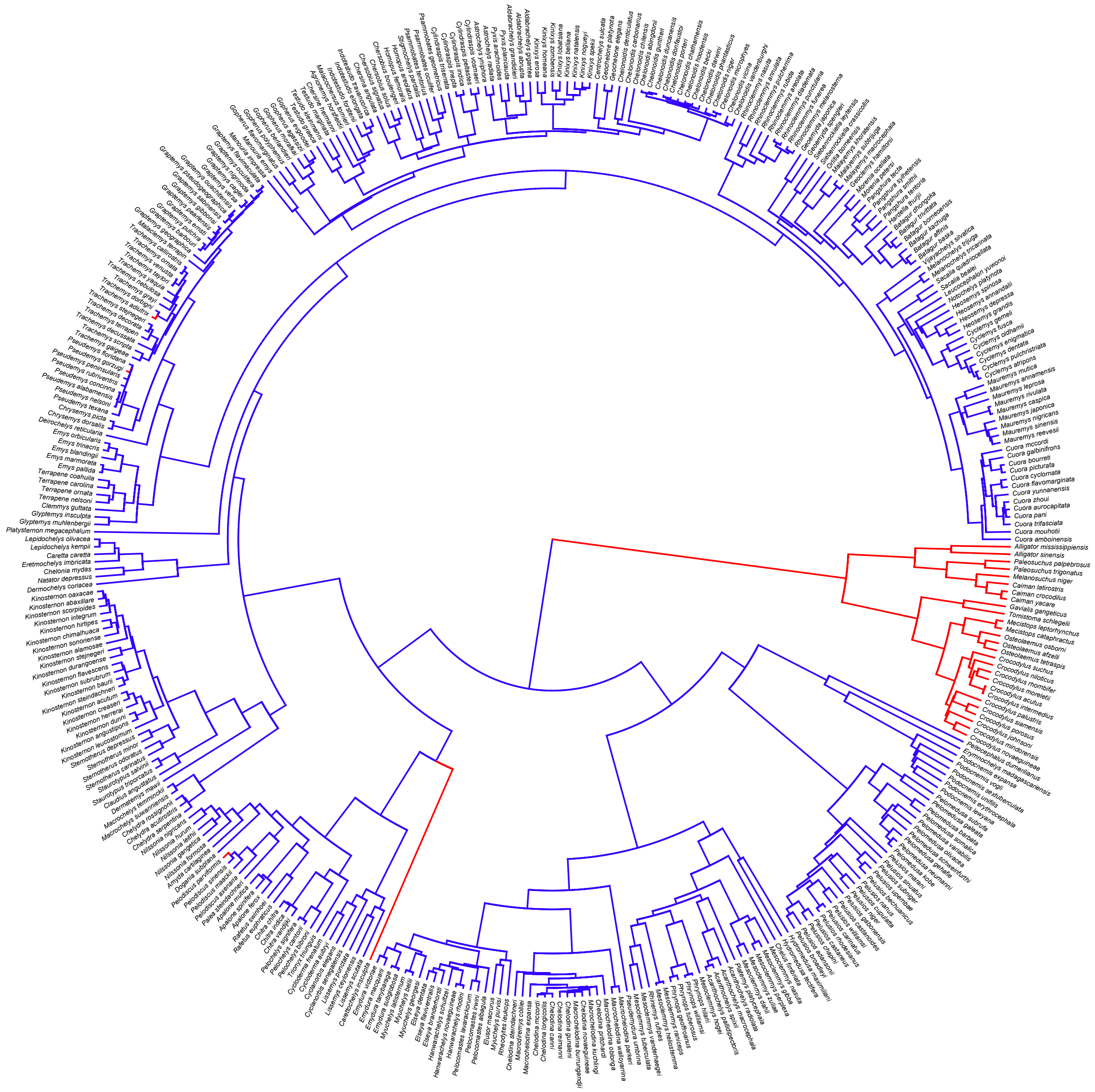

**Fig. S19.** Results from levolution analysis, showing location of jumps (Pp>0.85) under a Lévy process.

**Acknowledgements**

The silhouettes of *Araripemys* and *Gopherus* are reproduced here under a Creative Commons Attribution-NonCommercial 3.0 Unported license (https://creativecommons.org/licenses/by-nc/3.0/), and were generated by Aline M. Ghilardi, and Andrew A. Farke and Yan Wong, respectively. The remaining figures are freely distributed. All silhouettes are hosted on phylopic.org.

**Tables**

| **Turtles** | **Common Name** | **Historical Range** | **ED** | **EDGE** |
|---|---|---|---|---|
| *Erymnochelys madagascariensis* | Madagascan Big-Headed turtle | Madagascar | 88.81 | 7.27 |
| *Pseudemydura umbrina* | Western Swamp Turtle | Western Australia | 71.75 | 7.06 |
| *Carettochelys insculpta* | Fly River Pig-Nosed Turtle | Australia-New Guinea | 119.89 | 6.87 |
| *Dermatemys mawii* | Central American River Turtle | Southern Mexico to Guatemala | 57.85 | 6.85 |
| *Cyclanorbis elegans* | Nubian Flapshell Turtle | Western and Central Africa | 38.04 | 6.44 |
| *Podocnemis lewyana* | Magdalena River Turtle | Northern Colombia | 35.97 | 6.38 |
| *Platysternon megacephalum* | Big-Headed Turtle | Southeast Asia (Eastern Indochina) | 68.91 | 6.33 |
| *Rafetus swinhoei* | Yangtze Giant Softshell Turtle | Southeast Asia (Red and Yangtze Rivers) | 33.88 | 6.32 |
| *Mesoclemmys hogei* | Hoge's Side-Necked Turtle | Southeastern Brazil | 32.13 | 6.27 |
| *Siebenrockiella leytensis* | Philippine Forest Turtle | Philippines (Palawan) | 30.50 | 6.22 |
| **Crocodilians** | | | | |
| *Alligator sinensis* | Chinese Alligator | Southeast Asia (Yangtze River) | 53.21 | 6.77 |
| *Gavialis gangeticus* | Gharial | South Asia (Indus to Irawaddy Rivers) | 47.40 | 6.65 |
| *Mecistops cataphractus* | West African Slender-Snouted Crocodile | Western Africa | 28.25 | 6.15 |
| *Crocodylus mindorensis* | Philippine Crocodile | Philippines | 22.52 | 5.93 |
| *Crocodylus siamensis* | Siamese Crocodile | Southeast Asia | 20.76 | 5.85 |
| *Crocodylus rhombifer* | Cuban Crocodile | Cuba | 19.45 | 5.79 |
| *Crocodylus intermedius* | Orinoco Crocodile | Colombia and Venezuela | 17.64 | 5.70 |
| *Tomistoma schlegelii* | False Gharial | Southeast Asia | 47.40 | 5.27 |
| *Osteolaemus* sp. | Undescribed Dwarf Crocodile | Western Africa (west of Benin) | 26.38 | 4.70 |
| *Osteolaemus tetraspis* | Dwarf Crocodile | Western Africa (Benin to Gabon) | 26.38 | 4.70 |

**Table 1.** Top-10 EDGE species for turtles and crocodilians, showing scientific and common names, ranges, ED (Millions of Years), and EDGE scores. Compared to recent results (4), we do not estimate any sea turtles in the top 10 and place the Chinese Alligator (*Alligator sinensis*) ahead of the Gharial (*Gavialis gangeticus*) and the West African Slender-Snouted Crocodile (*Mecistops cataphractus*).

### Appendices

**Appendix S1.** Maximum-likelihood topology and support (1000 non-parametric bootstrap replicates) from the RAxML analysis of our supermatrix of up to 7 nuclear and 14 mitochondrial genes per species, containing up to 20,929bp for 340 turtles, 27 crocodilians, and 2 outgroups.

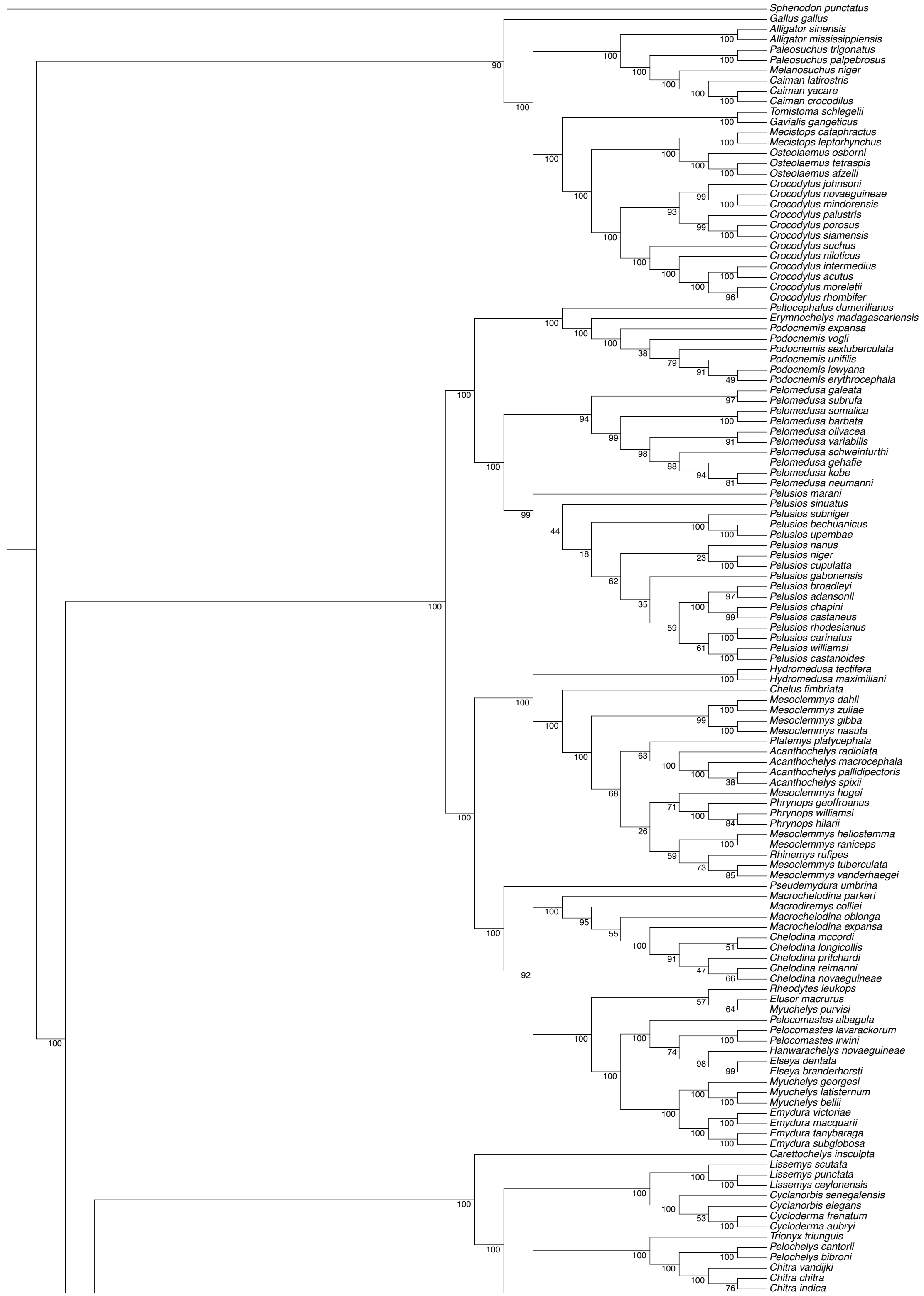
Sphenodon punctatus
Gallus gallus
Alligator sinensis
Alligator mississippiensis
Paleosuchus trigonatus
Paleosuchus palpebrosus
Melanosuchus niger
Caiman latirostris
Caiman yacare
Caiman crocodilus
Tomistoma schlegelii
Gavialis gangeticus
Mecistops cataphractus
Mecistops leptorhynchus
Osteolaemus osborni
Osteolaemus tetraspis
Osteolaemus afzelli
Crocodylus johnsoni
Crocodylus novaeguineae
Crocodylus mindorensis
Crocodylus palustris
Crocodylus porosus
Crocodylus siamensis
Crocodylus suchus
Crocodylus niloticus
Crocodylus intermedius
Crocodylus acutus
Crocodylus moreletii
Crocodylus rhombifer
Peltocephalus dumerilianus
Erymnochelys madagascariensis
Podocnemis expansa
Podocnemis vogli
Podocnemis sextuberculata
Podocnemis unifilis
Podocnemis lewyana
Podocnemis erythrocephala
Pelomedusa galeata
Pelomedusa subrufa
Pelomedusa somalica
Pelomedusa barbata
Pelomedusa olivacea
Pelomedusa variabilis
Pelomedusa schweinfurthi
Pelomedusa gehafie
Pelomedusa kobe
Pelomedusa neumanni
Pelusios marani
Pelusios sinuatus
Pelusios subniger
Pelusios bechuanicus
Pelusios upembae
Pelusios nanus
Pelusios niger
Pelusios cupulatta
Pelusios gabonensis
Pelusios broadleyi
Pelusios adansonii
Pelusios chapini
Pelusios castaneus
Pelusios rhodesianus
Pelusios carinatus
Pelusios williamsi
Pelusios castanoides
Hydromedusa tectifera
Hydromedusa maximiliani
Chelus fimbriata
Mesoclemmys dahli
Mesoclemmys zuliae
Mesoclemmys gibba
Mesoclemmys nasuta
Platemys platycephala
Acanthochelys radiolata
Acanthochelys macrocephala
Acanthochelys pallidipectoris
Acanthochelys spixii
Mesoclemmys hogei
Phrynops geoffroanus
Phrynops williamsi
Phrynops hilarii
Mesoclemmys heliostemma
Mesoclemmys raniceps
Rhinemys rufipes
Mesoclemmys tuberculata
Mesoclemmys vanderhaegei
Pseudemydura umbrina
Macrochelodina parkeri
Macrodiremys colliei
Macrochelodina oblonga
Macrochelodina expansa
Chelodina mccordi
Chelodina longicollis
Chelodina pritchardi
Chelodina reimanni
Chelodina novaeguineae
Rheodytes leukops
Elusor macrurus
Myuchelys purvisi
Pelocomastes albagula
Pelocomastes lavarackorum
Pelocomastes irwini
Hanwarachelys novaeguineae
Elseya dentata
Elseya branderhorsti
Myuchelys georgesi
Myuchelys latisternum
Myuchelys bellii
Emydura victoriae
Emydura macquarii
Emydura tanybaraga
Emydura subglobosa
Carettochelys insculpta
Lissemys scutata
Lissemys punctata
Lissemys ceylonensis
Cyclanorbis senegalensis
Cyclanorbis elegans
Cycloderma frenatum
Cycloderma aubryi
Trionyx triunguis
Pelochelys cantorii
Pelochelys bibroni
Chitra vandijki
Chitra chitra
Chitra indica

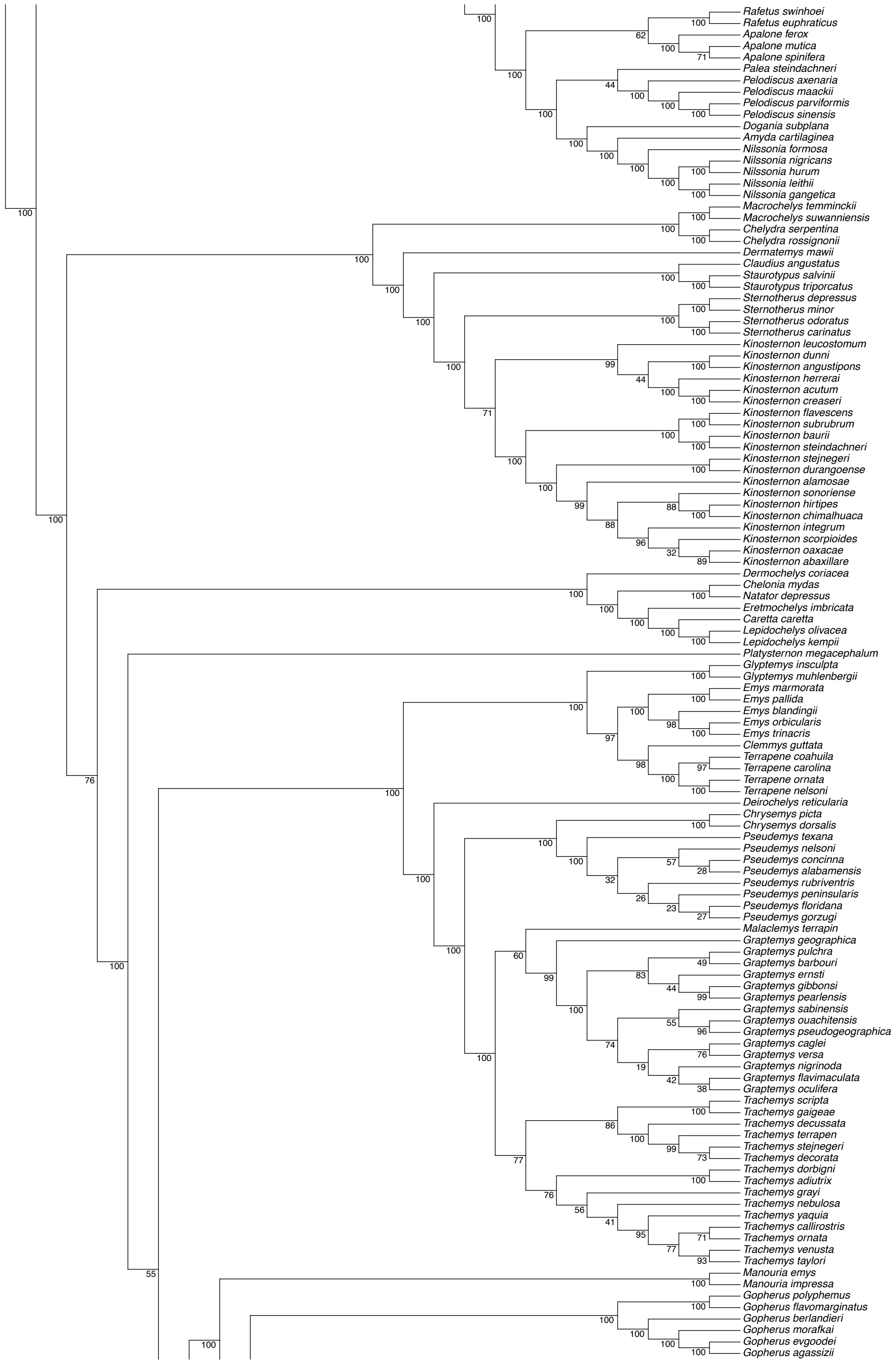

Rafetus swinhoei
Rafetus euphraticus
Apalone ferox
Apalone mutica
Apalone spinifera
Palea steindachneri
Pelodiscus axenaria
Pelodiscus maackii
Pelodiscus parviformis
Pelodiscus sinensis
Dogania subplana
Amyda cartilaginea
Nilssonia formosa
Nilssonia nigricans
Nilssonia hurum
Nilssonia leithii
Nilssonia gangetica
Macrochelys temminckii
Macrochelys suwanniensis
Chelydra serpentina
Chelydra rossignonii
Dermatemys mawii
Claudius angustatus
Staurotypus salvinii
Staurotypus triporcatus
Sternotherus depressus
Sternotherus minor
Sternotherus odoratus
Sternotherus carinatus
Kinosternon leucostomum
Kinosternon dunni
Kinosternon angustipons
Kinosternon herrerai
Kinosternon acutum
Kinosternon creaseri
Kinosternon flavescens
Kinosternon subrubrum
Kinosternon baurii
Kinosternon steindachneri
Kinosternon stejnegeri
Kinosternon durangoense
Kinosternon alamosae
Kinosternon sonoriense
Kinosternon hirtipes
Kinosternon chimalhuaca
Kinosternon integrum
Kinosternon scorpioides
Kinosternon oaxacae
Kinosternon abaxillare
Dermochelys coriacea
Chelonia mydas
Natator depressus
Eretmochelys imbricata
Caretta caretta
Lepidochelys olivacea
Lepidochelys kempii
Platysternon megacephalum
Glyptemys insculpta
Glyptemys muhlenbergii
Emys marmorata
Emys pallida
Emys blandingii
Emys orbicularis
Emys trinacris
Clemmys guttata
Terrapene coahuila
Terrapene carolina
Terrapene ornata
Terrapene nelsoni
Deirochelys reticularia
Chrysemys picta
Chrysemys dorsalis
Pseudemys texana
Pseudemys nelsoni
Pseudemys concinna
Pseudemys alabamensis
Pseudemys rubriventris
Pseudemys peninsularis
Pseudemys floridana
Pseudemys gorzugi
Malaclemys terrapin
Graptemys geographica
Graptemys pulchra
Graptemys barbouri
Graptemys ernsti
Graptemys gibbonsi
Graptemys pearlensis
Graptemys sabinensis
Graptemys ouachitensis
Graptemys pseudogeographica
Graptemys caglei
Graptemys versa
Graptemys nigrinoda
Graptemys flavimaculata
Graptemys oculifera
Trachemys scripta
Trachemys gaigeae
Trachemys decussata
Trachemys terrapen
Trachemys stejnegeri
Trachemys decorata
Trachemys dorbigni
Trachemys adiutrix
Trachemys grayi
Trachemys nebulosa
Trachemys yaquia
Trachemys callirostris
Trachemys ornata
Trachemys venusta
Trachemys taylori
Manouria emys
Manouria impressa
Gopherus polyphemus
Gopherus flavomarginatus
Gopherus berlandieri
Gopherus morafkai
Gopherus evgoodei
Gopherus agassizii

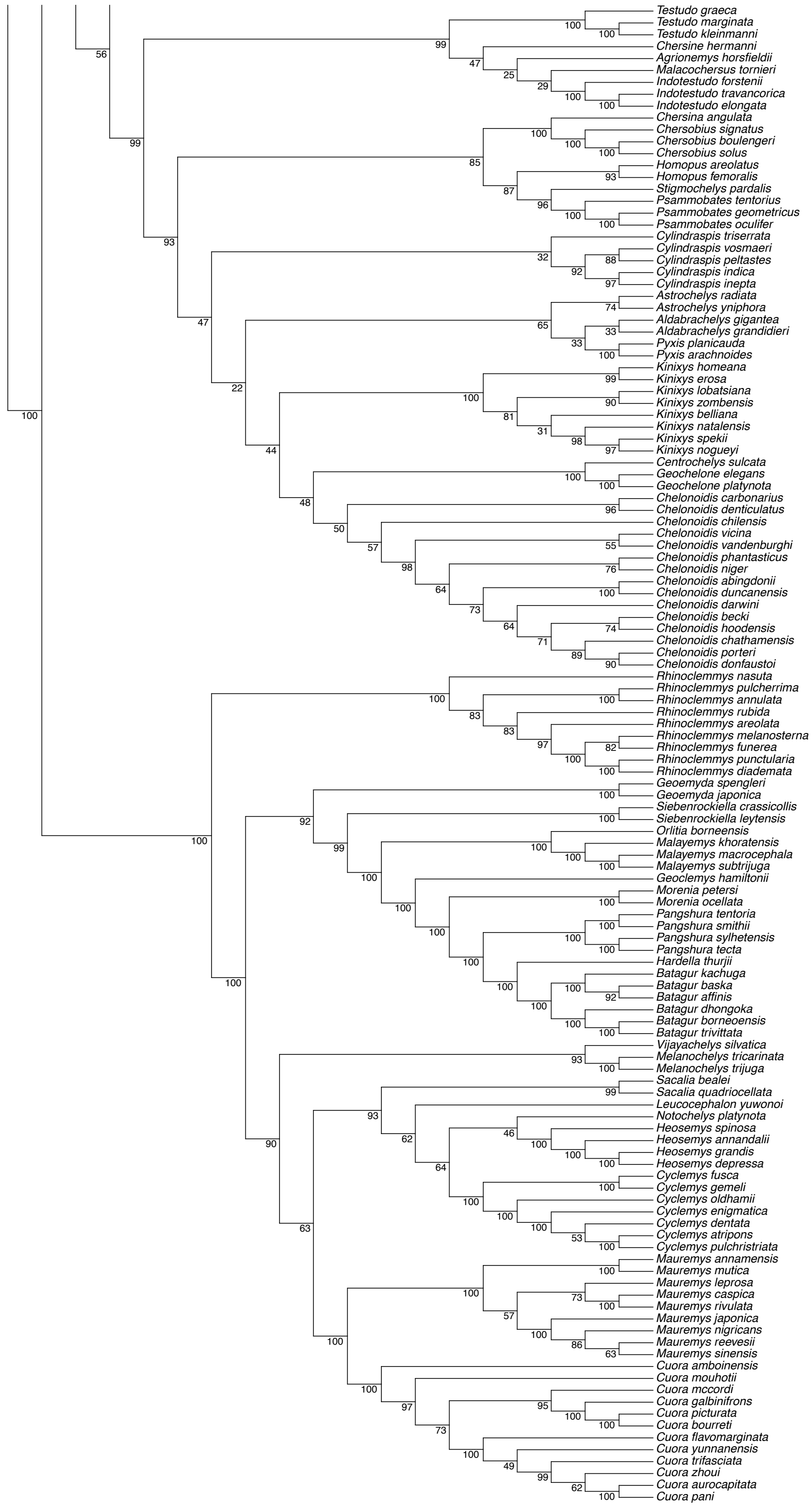
Testudo graeca
Testudo marginata
Testudo kleinmanni
Chersine hermanni
Agrionemys horsfieldii
Malacochersus tornieri
Indotestudo forstenii
Indotestudo travancorica
Indotestudo elongata
Chersina angulata
Chersobius signatus
Chersobius boulengeri
Chersobius solus
Homopus areolatus
Homopus femoralis
Stigmochelys pardalis
Psammobates tentorius
Psammobates geometricus
Psammobates oculifer
Cylindraspis triserrata
Cylindraspis vosmaeri
Cylindraspis peltastes
Cylindraspis indica
Cylindraspis inepta
Astrochelys radiata
Astrochelys yniphora
Aldabrachelys gigantea
Aldabrachelys grandidieri
Pyxis planicauda
Pyxis arachnoides
Kinixys homeana
Kinixys erosa
Kinixys lobatsiana
Kinixys zombensis
Kinixys belliana
Kinixys natalensis
Kinixys spekii
Kinixys nogueyi
Centrochelys sulcata
Geochelone elegans
Geochelone platynota
Chelonoidis carbonarius
Chelonoidis denticulatus
Chelonoidis chilensis
Chelonoidis vicina
Chelonoidis vandenburghi
Chelonoidis phantasticus
Chelonoidis niger
Chelonoidis abingdonii
Chelonoidis duncanensis
Chelonoidis darwini
Chelonoidis becki
Chelonoidis hoodensis
Chelonoidis chathamensis
Chelonoidis porteri
Chelonoidis donfaustoi
Rhinoclemmys nasuta
Rhinoclemmys pulcherrima
Rhinoclemmys annulata
Rhinoclemmys rubida
Rhinoclemmys areolata
Rhinoclemmys melanosterna
Rhinoclemmys funerea
Rhinoclemmys punctularia
Rhinoclemmys diademata
Geoemyda spengleri
Geoemyda japonica
Siebenrockiella crassicollis
Siebenrockiella leytensis
Orlitia borneensis
Malayemys khoratensis
Malayemys macrocephala
Malayemys subtrijuga
Geoclemys hamiltonii
Morenia petersi
Morenia ocellata
Pangshura tentoria
Pangshura smithii
Pangshura sylhetensis
Pangshura tecta
Hardella thurjii
Batagur kachuga
Batagur baska
Batagur affinis
Batagur dhongoka
Batagur borneoensis
Batagur trivittata
Vijayachelys silvatica
Melanochelys tricarinata
Melanochelys trijuga
Sacalia bealei
Sacalia quadriocellata
Leucocephalon yuwonoi
Notochelys platynota
Heosemys spinosa
Heosemys annandalii
Heosemys grandis
Heosemys depressa
Cyclemys fusca
Cyclemys gemeli
Cyclemys oldhamii
Cyclemys enigmatica
Cyclemys dentata
Cyclemys atripons
Cyclemys pulchristriata
Mauremys annamensis
Mauremys mutica
Mauremys leprosa
Mauremys caspica
Mauremys rivulata
Mauremys japonica
Mauremys nigricans
Mauremys reevesii
Mauremys sinensis
Cuora amboinensis
Cuora mouhotii
Cuora mccordi
Cuora galbinifrons
Cuora picturata
Cuora bourreti
Cuora flavomarginata
Cuora yunnanensis
Cuora trifasciata
Cuora zhoui
Cuora aurocapitata
Cuora pani

**Appendix S2.** Imputed body-lengths in cm (log) using the Rphylopars method (Goolsby et al. 2017).

| Species | Log(TL [cm]) |
|---|---|
| *Osteolaemus afzelli* | 5.28885877 |

**Appendix S3.** Imputed body-masses in g (log) using the Rphylopars method (Goolsby et al. 2017).

| Species | Log(mass[g] |
|---|---|
| *Acanthochelys radiolata* | 6.56116394 |
| *Aldabrachelys abrupta* | 11.8734733 |
| *Aldabrachelys grandidieri* | 11.6059184 |
| *Apalone mutica* | 7.71810422 |
| *Batagur trivittata* | 9.44901942 |
| *Chelodina canni* | 7.27479964 |
| *Chelodina gunaleni* | 7.12265037 |
| *Chelodina novaeguineae* | 7.48521924 |
| *Chelodina pritchardi* | 7.09316402 |
| *Chelodina steindachneri* | 6.91826542 |
| *Chelonoidis becki* | 11.6100879 |
| *Chelonoidis chathamensis* | 11.3416815 |
| *Chelonoidis darwini* | 11.4730349 |
| *Chelonoidis donfaustoi* | 11.9568293 |
| *Chelonoidis duncanensis* | 11.0726853 |
| *Chelonoidis guntheri* | 11.6925347 |
| *Chelonoidis hoodensis* | 11.1947687 |
| *Chelonoidis microphyes* | 11.5275682 |
| *Chelonoidis phantasticus* | 11.6705988 |
| *Chelonoidis porteri* | 11.9568293 |
| *Chelonoidis vandenburghi* | 11.8620628 |
| *Chelonoidis vicina* | 11.8620628 |
| *Chrysemys dorsalis* | 6.01095758 |
| *Claudius angustatus* | 6.5262252 |
| *Crocodylus suchus* | 11.7245643 |
| *Cuora aurocapitata* | 6.54185467 |

| | |
|---|---|
| *Cuora bourreti* | 6.90745227 |
| *Cuora cyclornata* | 7.66930615 |
| *Cuora pani* | 6.58295859 |
| *Cuora trifasciata* | 7.60351608 |
| *Cuora yunnanensis* | 6.36616797 |
| *Cuora zhoui* | 6.72844555 |
| *Cyclanorbis senegalensis* | 9.73123879 |
| *Cyclemys atripons* | 7.46684014 |
| *Cyclemys dentata* | 7.27733618 |
| *Cyclemys enigmatica* | 7.45994621 |
| *Cyclemys fusca* | 7.53806296 |
| *Cyclemys oldhamii* | 7.58617271 |
| *Cyclemys pulchristriata* | 7.40371486 |
| *Cylindraspis indica* | 10.0964931 |
| *Cylindraspis inepta* | 9.66321035 |
| *Cylindraspis peltastes* | 8.51865995 |
| *Cylindraspis triserrata* | 9.66321035 |
| *Cylindraspis vosmaeri* | 9.66321035 |
| *Dogania subplana* | 9.3045638 |
| *Elseya flaviventralis* | 7.724444 |
| *Emydura tanybaraga* | 7.05407521 |
| *Geoemyda japonica* | 6.24341896 |
| *Gopherus evgoodei* | 7.05645409 |
| *Gopherus flavomarginatus* | 8.30296414 |
| *Gopherus morafkai* | 7.5907716 |
| *Graptemys caglei* | 7.50392121 |
| *Graptemys ernsti* | 7.94381032 |
| *Graptemys nigrinoda* | 7.635763 |
| *Graptemys oculifera* | 7.68547324 |
| *Graptemys pearlensis* | 8.02952156 |
| *Graptemys pseudogeographica* | 7.78636869 |
| *Graptemys pulchra* | 7.82479212 |
| *Graptemys versa* | 7.53455453 |
| *Hanwarachelys rhodini* | 7.05610454 |
| *Hanwarachelys schultzei* | 7.41446824 |

| | |
|---|---|
| *Heosemys annandalii* | 9.14690364 |
| *Heosemys depressa* | 7.83025572 |
| *Kinixys lobatsiana* | 6.72714227 |
| *Kinixys natalensis* | 6.29934693 |
| *Kinixys nogueyi* | 6.88853054 |
| *Kinixys spekii* | 6.81636231 |
| *Kinixys zombensis* | 6.9540483 |
| *Kinosternon abaxillare* | 6.84740362 |
| *Kinosternon alamosae* | 5.69568153 |
| *Kinosternon angustipons* | 5.92501999 |
| *Kinosternon chimalhuaca* | 5.93001821 |
| *Kinosternon durangoense* | 5.80437346 |
| *Kinosternon flavescens* | 5.78319961 |
| *Kinosternon herrerai* | 6.05924643 |
| *Kinosternon oaxacae* | 6.14338778 |
| *Kinosternon steindachneri* | 5.27106296 |
| *Kinosternon stejnegeri* | 5.89156317 |
| *Lissemys ceylonensis* | 9.14210516 |
| *Lissemys scutata* | 8.31005757 |
| *Macrochelodina burrungandjii* | 7.55830688 |
| *Macrochelodina kuchlingi* | 7.95667987 |
| *Macrochelodina walloyarrina* | 7.27547819 |
| *Macrodiremys colliei* | 8.0694821 |
| *Malayemys khoratensis* | 7.51462704 |
| *Malayemys macrocephala* | 8.08318699 |
| *Malayemys subtrijuga* | 8.20402965 |
| *Mauremys japonica* | 6.37517762 |
| *Mauremys mutica* | 6.87314838 |
| *Mecistops leptorhynchus* | 10.9553276 |
| *Mesoclemmys heliostemma* | 7.45514403 |
| *Mesoclemmys nasuta* | 6.7492238 |
| *Mesoclemmys perplexa* | 5.96811058 |
| *Mesoclemmys raniceps* | 7.55664696 |
| *Mesoclemmys zuliae* | 6.41855981 |
| *Myuchelys purvisi* | 7.34181995 |

| | |
|---|---|
| *Nilssonia formosa* | 10.4069658 |
| *Nilssonia leithii* | 10.2915504 |
| *Notochelys platynota* | 8.16957675 |
| *Osteolaemus afzelli* | 10.29218 |
| *Osteolaemus osborni* | 9.44537514 |
| *Pelochelys signifera* | 10.0624217 |
| *Pelocomastes irwini* | 8.07317468 |
| *Pelodiscus axenaria* | 9.05959027 |
| *Pelodiscus maackii* | 10.0221656 |
| *Pelodiscus parviformis* | 7.89812604 |
| *Pelomedusa barbata* | 7.3046227 |
| *Pelomedusa galeata* | 7.97219865 |
| *Pelomedusa gehafie* | 6.99048031 |
| *Pelomedusa kobe* | 6.80721866 |
| *Pelomedusa neumanni* | 7.13022399 |
| *Pelomedusa olivacea* | 6.8966092 |
| *Pelomedusa schweinfurthi* | 6.78666752 |
| *Pelomedusa somalica* | 6.78666752 |
| *Pelomedusa variabilis* | 7.52891227 |
| *Peltocephalus dumerilianus* | 8.65542994 |
| *Pelusios bechuanicus* | 7.9776699 |
| *Pelusios broadleyi* | 6.70849628 |
| *Pelusios carinatus* | 7.80541959 |
| *Pelusios castaneus* | 7.70817418 |
| *Pelusios castanoides* | 7.37404571 |
| *Pelusios chapini* | 8.17523121 |
| *Pelusios cupulatta* | 7.38505227 |
| *Pelusios gabonensis* | 7.9660448 |
| *Pelusios marani* | 7.68737527 |
| *Pelusios nanus* | 6.32881163 |
| *Pelusios niger* | 8.06669255 |
| *Pelusios rhodesianus* | 7.54156718 |
| *Pelusios subniger* | 7.16465219 |
| *Pelusios upembae* | 7.39155822 |
| *Pelusios williamsi* | 7.50941725 |

| | |
|---|---|
| *Phrynops tuberosus* | 8.03379823 |
| *Phrynops williamsi* | 7.95315907 |
| *Pseudemys alabamensis* | 8.3675354 |
| *Pseudemys peninsularis* | 8.39336272 |
| *Pseudemys texana* | 8.06818659 |
| *Rafetus euphraticus* | 10.3024715 |
| *Rhinemys rufipes* | 7.19158605 |
| *Rhinoclemmys annulata* | 7.38558594 |
| *Rhinoclemmys diademata* | 8.03429284 |
| *Rhinoclemmys melanosterna* | 8.38266532 |
| *Rhinoclemmys punctularia* | 8.01522979 |
| *Rhinoclemmys rubida* | 7.38443562 |
| *Sacalia bealei* | 6.65465956 |
| *Siebenrockiella crassicollis* | 7.39242286 |
| *Staurotypus salvinii* | 7.20082201 |
| *Staurotypus triporcatus* | 7.88060579 |
| *Terrapene coahuila* | 6.53942711 |
| *Terrapene nelsoni* | 6.45943463 |
| *Trachemys decorata* | 8.1293629 |
| *Trachemys decussata* | 8.34734335 |
| *Trachemys gaigeae* | 8.17243308 |
| *Trachemys grayi* | 8.86352007 |
| *Trachemys nebulosa* | 7.98159522 |
| *Trachemys ornata* | 7.92904272 |
| *Trachemys stejnegeri* | 7.76827515 |
| *Trachemys taylori* | 6.94066228 |
| *Trachemys terrapen* | 8.02616969 |
| *Trachemys yaquia* | 7.60614277 |

**Appendix S4.** Imputed climate variables using the Rphylopars method (Goolsby et al. 2017).

| **Species** | **BIO36** | **BIO37** | **BIO38** | **BIO39** | **BIO40** |
|---|---|---|---|---|---|
| *Aldabrachelys_gigantea* | 1.84648008 | 2.61016789 | 0.86300135 | 0.17724099 | -1.3567541 |
| *Chelonoidis_abingdonii* | 1.79377432 | 0.1168676 | 1.94038523 | -0.3537702 | -1.7896385 |

| | | | | | |
|---|---|---|---|---|---|
| *Chelonoidis_duncanensis* | 1.68478942 | -0.3089224 | 2.29119698 | -0.1088011 | -2.0738076 |
| *Chelonoidis_hoodensis* | 1.69175342 | 0.52644942 | 0.83289899 | -0.6580248 | -0.9751086 |

**Appendix S5.** Imputed threat statuses the random forest (RF), neural network (NN), and PGLM models.

| Species | RF | NN | PGLM |
|---|---|---|---|
| *Chelodina canni* | LC | VU | NT |
| *Chelodina gunaleni* | EN | NT | NT |
| *Chelodina longicollis* | LC | LC | NT |
| *Chelodina steindachneri* | LC | VU | NT |
| *Chelonoidis carbonarius* | VU | VU | NT |
| *Chelus fimbriata* | VU | VU | NT |
| *Chelydra acutirostris* | VU | VU | NT |
| *Chitra vandijki* | CR | CR | CR |
| *Chrysemys dorsalis* | LC | LC | NT |
| *Crocodylus suchus* | CR | LC | VU |
| *Cuora cyclornata* | CR | EN | CR |
| *Cyclemys atripons* | CR | VU | EN |
| *Cyclemys enigmatica* | CR | EN | EN |
| *Cyclemys fusca* | CR | CR | CR |
| *Cyclemys gemeli* | CR | EN | EN |
| *Cyclemys oldhamii* | VU | CR | VU |
| *Cyclemys pulchristriata* | CR | CR | CR |
| *Deirochelys reticularia* | LC | LC | NT |
| *Elseya dentata* | LC | VU | NT |
| *Elseya flaviventralis* | LC | VU | VU |
| *Emydura macquarii* | LC | LC | NT |

| | | | |
|---|---|---|---|
| *Emydura tanybaraga* | LC | VU | NT |
| *Emydura victoriae* | LC | VU | VU |
| *Emys pallida* | VU | CR | EN |
| *Emys trinacris* | CR | CR | EN |
| *Gopherus morafkai* | VU | LC | NT |
| *Graptemys sabinensis* | VU | EN | NT |
| *Hanwarachelys rhodini* | LC | LC | LC |
| *Hanwarachelys schultzei* | VU | LC | NT |
| *Hydromedusa tectifera* | LC | VU | NT |
| *Kinixys belliana* | VU | VU | NT |
| *Kinixys erosa* | VU | VU | VU |
| *Kinixys nogueyi* | VU | VU | VU |
| *Kinixys spekii* | LC | VU | NT |
| *Kinixys zombensis* | VU | VU | VU |
| *Kinosternon abaxillare* | VU | CR | VU |
| *Kinosternon alamosae* | LC | NT | NT |
| *Kinosternon durangoense* | LC | CR | VU |
| *Kinosternon leucostomum* | NT | NT | NT |
| *Kinosternon oaxacae* | CR | NT | VU |
| *Kinosternon scorpioides* | NT | NT | NT |
| *Kinosternon steindachneri* | LC | NT | LC |
| *Lissemys ceylonensis* | VU | VU | VU |
| *Lissemys scutata* | VU | EN | EN |
| *Macrochelodina burrungandjii* | VU | VU | VU |
| *Macrochelodina expansa* | LC | LC | VU |
| *Macrochelodina kuchlingi* | VU | EN | EN |

| | | | |
|---|---|---|---|
| *Macrochelodina walloyarrina* | NT | EN | VU |
| *Macrochelys suwanniensis* | EN | EN | VU |
| *Macrodiremys colliei* | VU | EN | EN |
| *Malayemys khoratensis* | CR | CR | EN |
| *Malayemys macrocephala* | VU | VU | EN |
| *Mauremys caspica* | VU | VU | VU |
| *Mauremys leprosa* | VU | NT | NT |
| *Mauremys rivulata* | VU | LC | NT |
| *Mecistops leptorhynchus* | LC | LC | VU |
| *Mesoclemmys gibba* | VU | VU | NT |
| *Mesoclemmys heliostemma* | VU | VU | NT |
| *Mesoclemmys nasuta* | VU | VU | VU |
| *Mesoclemmys perplexa* | CR | CR | EX |
| *Mesoclemmys raniceps* | VU | VU | NT |
| *Mesoclemmys tuberculata* | VU | VU | NT |
| *Myuchelys georgesi* | EN | EN | EN |
| *Myuchelys latisternum* | LC | LC | NT |
| *Myuchelys purvisi* | EN | EN | EN |
| *Natator depressus* | VU | VU | VU |
| *Osteolaemus afzelli* | CR | LC | VU |
| *Osteolaemus osborni* | LC | LC | NT |
| *Pelocomastes albagula* | LC | EN | VU |
| *Pelocomastes irwini* | VU | EN | VU |
| *Pelocomastes lavarackorum* | VU | EN | VU |
| *Pelodiscus axenaria* | EN | EN | CR |
| *Pelodiscus maackii* | LC | CR | VU |

| | | | |
|---|---|---|---|
| *Pelodiscus parviformis* | EN | EN | CR |
| *Pelomedusa barbata* | VU | VU | VU |
| *Pelomedusa gehafie* | VU | VU | NT |
| *Pelomedusa kobe* | VU | LC | VU |
| *Pelomedusa neumanni* | VU | LC | NT |
| *Pelomedusa olivacea* | LC | LC | NT |
| *Pelomedusa schweinfurthi* | LC | LC | NT |
| *Pelomedusa somalica* | VU | LC | NT |
| *Pelomedusa subrufa* | LC | LC | NT |
| *Pelomedusa variabilis* | VU | VU | EN |
| *Pelusios adansonii* | LC | LC | NT |
| *Pelusios bechuanicus* | LC | LC | NT |
| *Pelusios carinatus* | VU | LC | VU |
| *Pelusios castaneus* | VU | LC | VU |
| *Pelusios chapini* | VU | LC | VU |
| *Pelusios cupulatta* | VU | VU | EN |
| *Pelusios gabonensis* | LC | LC | NT |
| *Pelusios marani* | VU | VU | VU |
| *Pelusios nanus* | LC | LC | NT |
| *Pelusios niger* | VU | VU | EN |
| *Pelusios sinuatus* | LC | LC | NT |
| *Pelusios upembae* | CR | CR | EN |
| *Pelusios williamsi* | VU | VU | VU |
| *Phrynops geoffroanus* | VU | VU | LC |
| *Phrynops hilarii* | VU | VU | NT |
| *Phrynops tuberosus* | VU | VU | NT |

| | | | |
|---|---|---|---|
| *Platemys platycephala* | VU | VU | NT |
| *Podocnemis vogli* | VU | VU | EN |
| *Psammobates oculifer* | LC | LC | NT |
| *Pseudemys floridana* | NT | LC | NT |
| *Rhinoclemmys diademata* | VU | CR | VU |
| *Rhinoclemmys melanosterna* | NT | NT | VU |
| *Rhinoclemmys pulcherrima* | NT | NT | NT |
| *Rhinoclemmys punctularia* | VU | VU | NT |
| *Terrapene nelsoni* | VU | VU | VU |
| *Trachemys callirostris* | VU | VU | VU |
| *Trachemys decussata* | VU | VU | NT |
| *Trachemys dorbigni* | VU | VU | VU |
| *Trachemys grayi* | VU | VU | VU |
| *Trachemys nebulosa* | VU | VU | NT |
| *Trachemys venusta* | VU | VU | NT |
